# Pathway Anchored Multimodal Clustering Reveals Circuit Level Signatures in Parkinsons Disease

**DOI:** 10.64898/2025.12.15.694278

**Authors:** Ashwin Vinod, Aditya Sai Ellendula, Shubham Bhardwaj, Aparna Dev, Aaron Dominic, Chandrajit Bajaj

**Affiliations:** Department of Computer Science, The University of Texas at Austin, Austin, TX, USA; Department of Computer Science, Amrita Vishwa Vidyapeetham, Coimbatore, India

**Author notes:** These authors contributed equally (Ashwin Vinod, Aditya Sai Ellendula).

## Abstract

Parkinson’s disease is increasingly understood as a disorder of distributed brain circuits, yet most imaging analyses do not explicitly respect pathway structure. We introduce a pathway-anchored, multimodal clustering framework based on Scalable Robust Variational Compositional Co-clustering (SRVCC) that integrates structural MRI, free-water–corrected diffusion MRI, and DAT-SPECT in anatomically defined circuits. For each pathway, we derive a simple Multimodal Pathway Integrity Score (MPIS) that aggregates *z*-normalised volume, microstructural, and dopaminergic measures into an interpretable summary of imaging integrity. In the PPMI cohort, SRVCC identifies stable imaging-derived patient clusters and feature modules under explicit model selection and bootstrap/stability checks, with covariate-adjusted analyzes controlling for age, sex, education, and medication. MPIS shows coherent but modest structure–function associations: lower nigrostriatal and frontostriatal integrity relates to higher motor burden (UPDRS-III), while reduced sensory/visuospatial and limbic integrity is linked to lower global cognition (MoCA); microvascular markers robustly stratify imaging profiles but display minimal cross-sectional coupling to these global scales. Feature-level reports highlight dominant region–by–modality contributors (e.g., striatal DAT-SBR, thalamic and cerebellar morphology, white-matter hyperintensity metrics), providing a transparent bridge from multimodal data to circuit-level signatures. This pathway-aware representation offers a principled, reproducible way to summarise multimodal imaging in PD and may support future work on circuit-informed stratification, prognosis, and targeted outcome measures.

## 1 Introduction

Parkinson’s disease (PD) shows substantial heterogeneity in symptoms, progression, and treatment response across patients[17]. In routine care, severity is typically summarized with bedside scales such as the Movement Disorder Society Unified Parkinson’s Disease Rating Scale Part III (MDS-UPDRS III) for motor impairment, the Montreal Cognitive Assessment (MoCA) for global cognition, and the Questionnaire for Impulsive-Compulsive Disorders in Parkinson’s Disease (QUIP) for behavioral symptoms[18, 32, 53]. These instruments standardize symptom scoring but mainly capture functional consequences rather than underlying brain pathology. Neuroimaging provides complementary biological information: dopamine transporter single-photon emission computed tomography (DaT-SPECT) indexes striatal dopaminergic denervation[31, 16], structural MRI captures macroscopic atrophy[22], and diffusion tensor imaging (DTI) and free-water DTI probe white-matter microstructure[36, 42]. Yet many studies evaluate modalities in isolation or prioritize global whole-brain signatures, leaving open whether *multimodal, biology-anchored* integration can expose coherent structure within the PD spectrum.

PD is increasingly understood as a distributed network disorder in which dopaminergic loss in the substantia nigra perturbs basal ganglia–thalamo–cortical loops subserving motor control, executive function, and limbic processing[34]. Heterogeneous involvement of additional systems including frontostriatal and cerebello–thalamo–cortical circuits, cholinergic basal forebrain and pedunculopontine pathways, sensory/visuospatial networks, and microvascular burden contributes to variability in gait, cognition, and behavioral symptoms[45, 39]. Analyses that treat regions as exchangeable features risk groupings driven by coincident covariation rather than mechanism-linked circuitry; a pathway-centric strategy instead respects known anatomical coupling and allows results to be interpreted directly in biological terms, for example as “predominantly nigrostriatal” versus “frontostriatal–executive” involvement.

Over the past decade, multivariate and multimodal frameworks have been developed to fuse imaging modalities and relate them to clinical or genetic variation. Classical approaches such as canonical correlation analysis (CCA)[13] and joint, parallel, or linked independent component analysis (joint/parallel ICA, linked ICA)[49, 19, 30] decompose multimodal data into shared latent components, and multimodal classifiers that combine DaT-SPECT, DTI, and volumetry improve diagnostic or prognostic performance over single-modality markers in PD and atypical parkinsonian syndromes[27, 2]. Large consortia such as ENIGMA-PD have mapped stage-dependent white-matter alterations across Hoehn–Yahr stages using harmonized DTI[39]. More recently, large-scale integration efforts combine longitudinal neuroimaging, CSF, and multi-omics to derive “pace” subtypes and candidate repurposable drugs[48], link gene variants to multimodal brain changes and symptom trajectories via imaging–genomics models[1], and release standardized imaging-derived phenotypes from PPMI to facilitate joint ICA/CCA-style analyses[3]. Other fusion pipelines integrate MRI with genetics[55], diffusion and dopaminergic imaging with clinical scales[54], or neuroimaging with gut microbiome profiles[15] to improve diagnosis or progression prediction. However, these whole-brain or imaging–omics fusion methods typically operate in a latent space whose spatial loadings can be anatomically diffuse and do not explicitly enforce pathway-wise structure or yield patient clusters and feature modules that can be read directly as circuit-level “signatures.”

Unsupervised PD subtyping studies using structural or diffusion features have begun to reveal imaging-defined subgroups with differing motor and cognitive trajectories[23, 58], and supervised radiomics models further distinguish tremor-dominant versus postural-instability/gait-difficulty phenotypes[38]. Yet most existing work either (i) focuses on a single modality, (ii) learns global latent patterns without explicit anatomical binning, or (iii) does not systematically quantify how robust multimodal composites are to modeling choices such as modality weighting, intracranial volume (ICV) normalization, or sign conventions. This motivates an approach that integrates multiple imaging modalities, aggregates features within *predefined neurobiological pathways*, and explicitly assesses robustness and clinical relevance of the resulting circuit-level indices.

Taken together, these observations argue for a representation that preserves modality complementarity and anatomical specificity. In this work, we operationalize a *pathway-anchored stratification* strategy by summarizing multimodal features within each circuit using a *Multimodal Pathway Integrity Score* (MPIS) and then examining (a) pathway-wise separations across patients and (b) structure–function links between MPIS and clinical measures. Instead of assigning opaque, geometry-driven cluster labels, this framework is designed to support statements such as “patients in this subtype show disproportionately low nigrostriatal integrity relative to limbic and sensory pathways.”

We aggregate harmonized structural MRI volumetry, diffusion-derived fractional anisotropy (FA), mean diffusivity (MD) and free water, and DaT-SPECT specific binding ratios within six predefined PD-relevant circuits namely nigrostriatal motor, frontostriatal–executive, limbic/mesolimbic, cerebello–thalamo–cortical, microvascular-burden, and sensory–visuospatial–attention to compute per-subject MPIS for each pathway. We then apply a Scaled Robust Variational Co-Clustering (SRVCC) [52] model to the standardized subject×feature matrix, derive pathway-specific MPIS composites, and examine their separations across imaging-driven patient clusters, as well as their associations with MDS-UPDRS III, MoCA, and QUIP under Benjamini–Hochberg false discovery rate control. In addition, we evaluate MPIS variants that differ in modality weighting and ICV normalization, and quantify SRVCC cluster stability across random initializations and bootstrap resamples.

Our contributions are threefold. First, we present a pathway-stratified SRVCC framework that respects neurobiological circuit organization while enabling data-driven discovery of imaging-defined patient strata and feature modules. Second, we demonstrate that pathway-specific MPIS composites exhibit coherent structure–function relationships aligned with known PD pathophysiology: nigrostriatal and frontostriatal integrity track motor severity, limbic and microvascular pathways relate to behavioral and gait burden, and sensory–visuospatial integrity contributes to cognitive variation. Third, we provide a transparent reporting template linking imaging clusters to specific region–modality biomarkers, pathway-level MPIS profiles, and clinical phenotypes, facilitating replication and mechanistic interpretation.

The remainder of the paper details the cohort, imaging preprocessing, and feature-extraction pipeline (Methods), defines MPIS and the co-clustering and validation procedures, and presents pathway-wise results followed by implications for precision imaging–based stratification in Parkinson’s disease.

## 2 Methods

### 2.1 Participants and Dataset

Data for this study were sourced from the Parkinson’s Progression Markers Initiative (PPMI)[28], a large-scale, open-access longitudinal study aimed at identifying biomarkers of Parkinson’s disease (PD) progression. The dataset includes a total of 294 participants, comprising 185 individuals diagnosed with Parkinson’s disease (62.9%), 72 healthy controls (24.5%), and 37 participants (12.6%) classified as Scans Without Evidence of Dopaminergic Deficiency (SWEDD). Individuals in the PD cohort exhibited two or more cardinal motor signs (rest tremor, bradykinesia, rigidity) along with an abnormal dopaminergic deficit on DaTSCAN SPECT imaging. Healthy controls are individuals without a diagnosis of any neurological disorder, no first-degree family history of PD, normal DaTSCAN results, and no clinical signs of parkinsonism. Participants classified as SWEDD exhibit clinical features resembling parkinsonism but demonstrate normal presynaptic dopaminergic function on DaTSCAN SPECT[7]. All included patients had T1-weighted MRI, diffusion MRI, and DaTSCAN SPECT available for analysis. DaTSCAN SPECT data were used for pathway-specific analyses. Summary demographics (age, sex, education, disease duration, and medication status), along with baseline clinical scores, are reported in Table 1.

**Table 1:** Demographic and clinical characteristics of the study cohort. Values are mean *±* SD unless otherwise indicated.

|  | PD (n=185) | HC (n=72) | SWEDD (n=37) |
| --- | --- | --- | --- |
| Age (years) | 64.2 ± 9.1 | 61.8 ± 8.7 | 63.5 ± 8.9 |
| Female (%) | 38.9 | 40.3 | 35.1 |
| Education (years) | 15.2 ± 3.1 | 15.8 ± 2.9 | 15.0 ± 3.0 |
| Disease duration (years) | 1.9 ± 1.1 | – | 1.8 ± 1.0 |
| Medication status (% on dopaminergic therapy) | 52.4 | – | 48.6 |
| MoCA score | 26.3 ± 2.4 | 28.2 ± 1.5 | 26.9 ± 2.1 |
| MDS-UPDRS III | 21.7 ± 9.6 | 1.6 ± 2.1 | 17.3 ± 8.4 |

#### Clinical Assessments

Standardized clinical evaluations included the Unified Parkinson’s Disease Rating Scale (MDS-UPDRS), the Montreal Cognitive Assessment (MoCA), and the Questionnaire for Impulsive-Compulsive Disorders in Parkinson’s Disease (QUIP). MDS-UPDRS Part III (Motor Examination) is a standardized clinical test to quantify the severity and progression of *motor* features of PD, such as bradykinesia, rigidity, tremor, posture, and gait through structured clinician-rated assessments, with a higher score indicating greater motor impairment[37]. MDS-UPDRS Parts I and II are questionnaires assessing non-motor and motor experiences of daily living, respectively, while Part IV evaluates treatment-related motor complications such as dyskinesias and motor fluctuations. MoCA is a brief 30-point screening tool designed to detect mild cognitive impairment in multiple domains, including attention, executive function, memory, language, visuospatial ability, and orientation[32]. It is widely employed in PD cohorts owing to its high sensitivity in detecting early and subtle cognitive impairments. The Questionnaire for Impulsive-Compulsive Disorders in Parkinson’s Disease (QUIP) is a validated self-administered screening instrument designed to assess impulsive and compulsive behaviors commonly associated with dopaminergic therapy, including pathological gambling, hypersexuality, compulsive buying, binge eating, hobbyism, and behaviors related to medication overuse[53]. It serves as an important clinical measure for identifying behavioral dysregulation and its potential association with dopaminergic treatment. In the feature matrix used for co-clustering, we retained the total numeric score for Part III of the MDS-UPDRS, the MoCA total score, and the QUIP summary score; together these variables define the clinical-only feature view V1 (Table 2).

**Table 2:** Feature views for ablation analysis.

| View | Feature Set | *n* features | Purpose |
| --- | --- | --- | --- |
| V1 | Clinical only<br>UPDRS-III, UPDRS-total, MoCA, QUIP | 4 | Baseline behavioral phenotype |
| V2 | V1 + Structural + DAT<br>+ T1 volumes (27)<br>+ DaTSCAN SBR/mean/std (17) | 48 | Macrostructure + dopamine |
| V3 | V2 + DTI (uncorrected)<br>+ FA/MD mean/std (64) | 112 | Standard microstructure |
| V4 | V2 + FWE-DTI<br>+ FA<sub>T</sub>/MD<sub>T</sub>/FW mean/std (96) | 144 | CSF-corrected microstructure<br>(primary analysis) |

#### Imaging Modalities

PPMI provides three primary neuroimaging modalities: diffusion tensor imaging (DTI), T1-weighted magnetic resonance imaging (MRI), and DaTSCAN single photon emission computed tomography (SPECT). DTI provides insights into white-matter microstructure and connectivity by quantifying both the directionality (anisotropy) and magnitude of water diffusion within tissue[5]. Voxel-wise maps of fractional anisotropy (FA) and mean diffusivity (MD), derived from DTI, serve as indices of microstructural integrity, with higher FA values reflecting greater directional coherence of diffusion and thus preserved axonal organization. T1-weighted MRI provides high-resolution structural information, enabling the assessment of brain morphology and regional atrophy associated with PD, particularly in regions such as the basal ganglia, thalamus, and frontal cortex[51]. DaTSCAN SPECT is employed as a presynaptic dopaminergic biomarker, leveraging the radiotracer Ioflupane (^123^I, [^123^I]FP-CIT) to visualise dopamine transporter (DAT) density within the striatum; reduced tracer binding reflects the loss of nigrostriatal dopaminergic neurons, enabling differentiation of neurodegenerative parkinsonian syndromes at early stages[6]. These modalities yield regional T1 volumes, diffusion-derived FA/MD (and related microstructural metrics), and striatal DaTSCAN specific binding ratios that are subsequently assembled into multimodal feature sets. For downstream analyses, we define four nested feature views (V1–V4) that progressively add imaging information on top of the clinical scores—from clinical-only (V1) to clinical + T1/DaTSCAN + diffusion/FWE-DTI (V4); the exact composition of each view is summarised in Table 2. Acquisition parameters and modality-specific quality-control procedures, including motion/outlier handling and visual plus quantitative checks of DaTSCAN-to-T1 coregistration, are detailed in Section 2.2.

### 2.2 Imaging Preprocessing and Quality Control

We convert raw multimodal imaging into pathway-aware features through a lean, reproducible pipeline (Fig. 2). First, T1-weighted MRI is segmented to define a subject-specific anatomical reference space; all other modalities are rigidly aligned to this space after skull stripping. Within this reference, we extract ROI-wise structural volumes, diffusion metrics (DTI-FA and DTI-MD), and DAT-SPECT specific binding ratios (SBR). Region-level features are then aggregated using predefined circuit masks; in the next section we formalize this aggregation as the *Multimodal Pathway Integrity Score* (MPIS) and describe clustering and evaluation in pathway space. See the pathway masks in Fig. 3 (CTC, limbic, microvascular, sensory) and Fig. 4 (frontostriatal, nigrostriatal).

**Figure 1.**
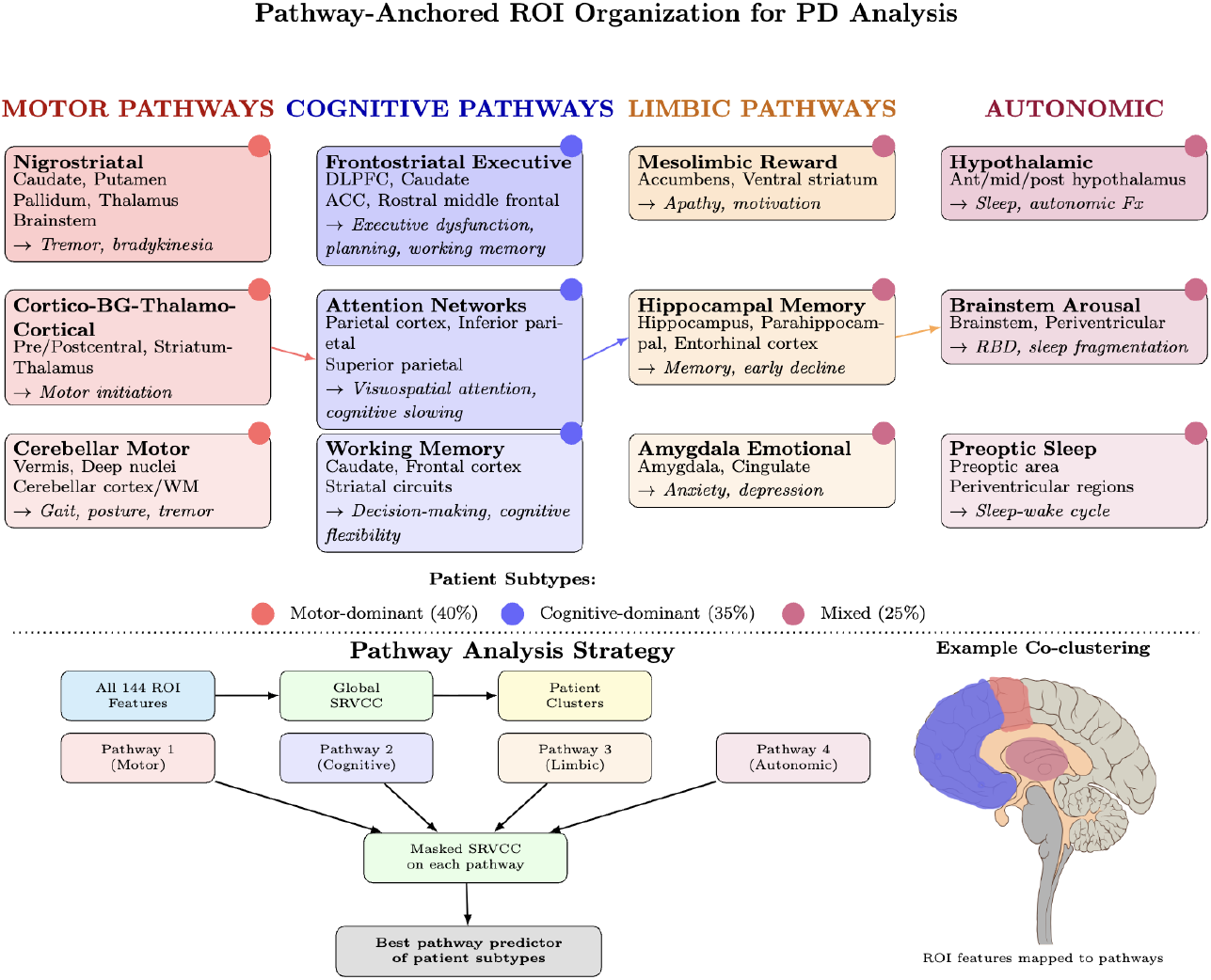
Pathway-anchored ROI organization for mechanistic interpretation of co-clusters. Regions are grouped into five major systems implicated in PD pathophysiology. Arrows indicate cross-system integration (motor-cognitive, cognitive-limbic, limbic-autonomic). The bottom panel shows the pathway analysis workflow used to test whether discovered patient subtypes align with known neurobiological circuits.

**Figure 2.**
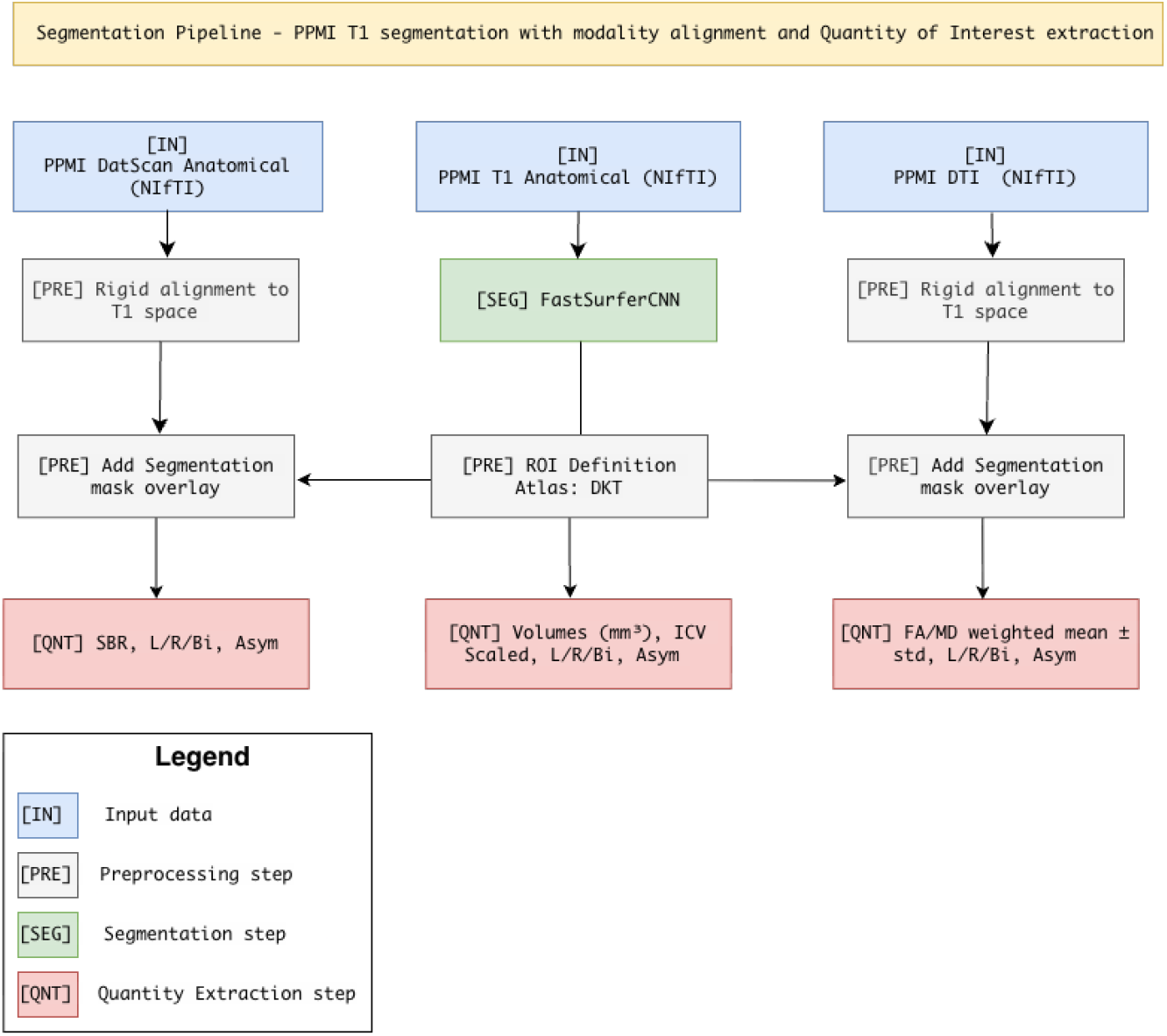
Multimodal segmentation and quantification pipeline.^*^ T1 MRI volumes are segmented with FastSurferCNN to generate DKT-atlas ROIs in subject-native space. These ROIs guide rigid alignment of DaTSCAN SPECT and DTI (FA/MD) images, after which the segmentation mask is overlaid for region-wise quantification: T1 → volumes, DTI → FA/MD weighted mean +− SD, and DaTSCAN → mean uptake and SBR = (ROI/ref). All metrics are computed per hemisphere and bilaterally, with asymmetry indices (Asym = (RL)/(R+L)).

**Figure 3.**
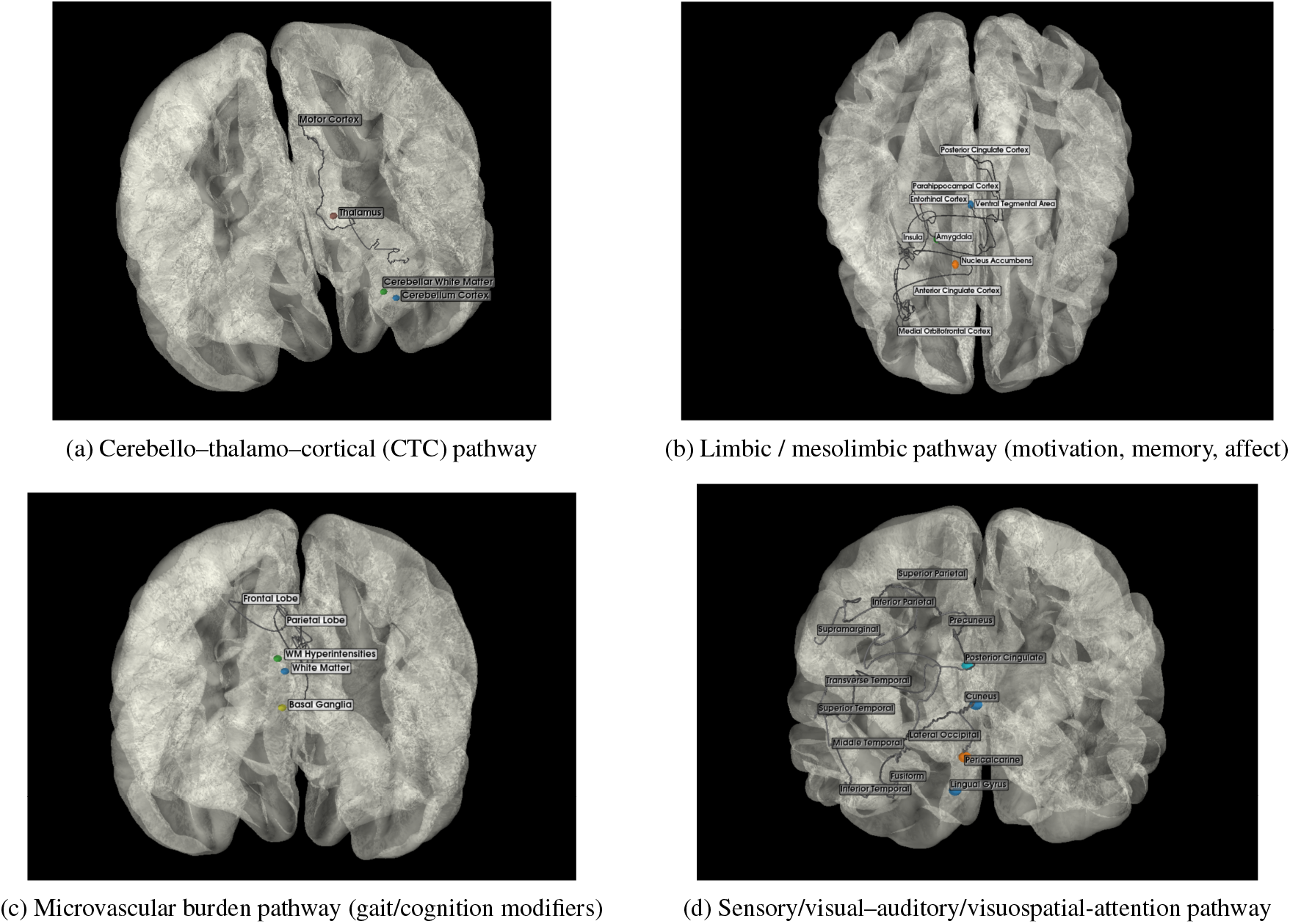
Pathway schematics for (A) cerebello–thalamo–cortical, (B) limbic/mesolimbic, (C) microvascular-burden, and (D) sensory/visual–auditory/visuospatial-attention pathways.

**Figure 4.**
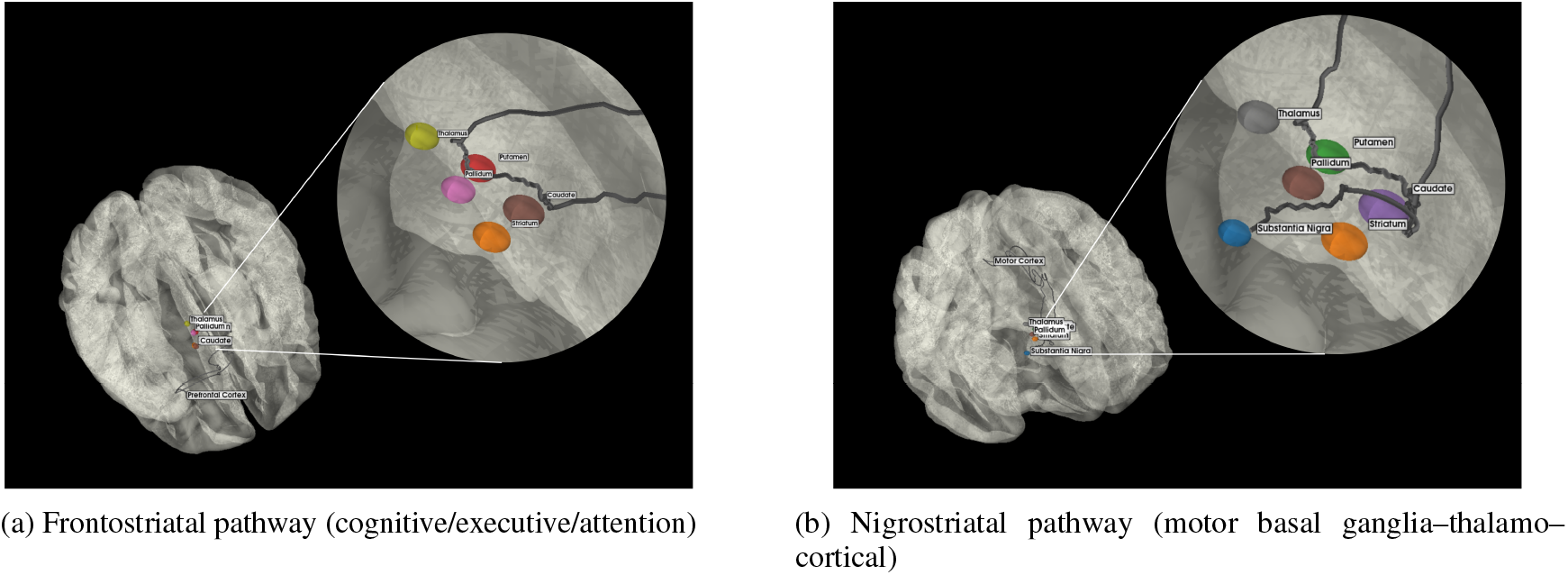
Pathway schematics of (A) frontostriatal (cognitive/executive/attention) and (B) nigrostriatal (motor basal ganglia–thalamo–cortical).

**Figure 5.**
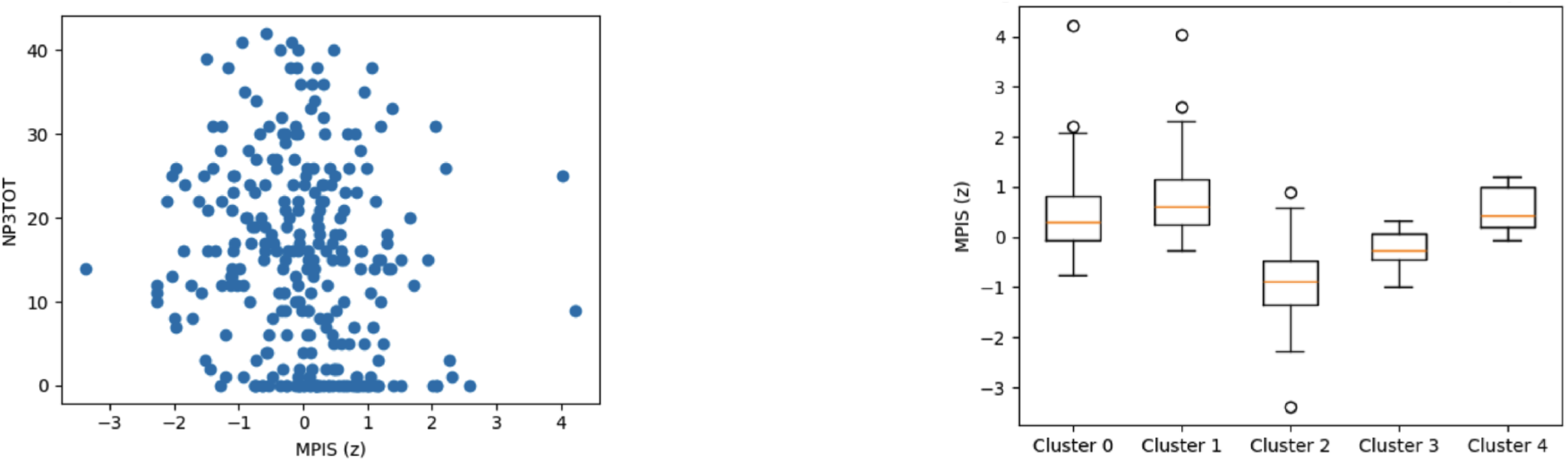
Nigrostriatal MPIS associations. Left: MPIS vs MDS-UPDRS III (Spearman *ρ* ≈ −0.201, *q* ≈ 6.8 × 10^−4^). Right: MPIS separation across clusters (Kruskal–Wallis *H* ≈ 167.15, *η*^2^ ≈ 0.587).

**Figure 6.**
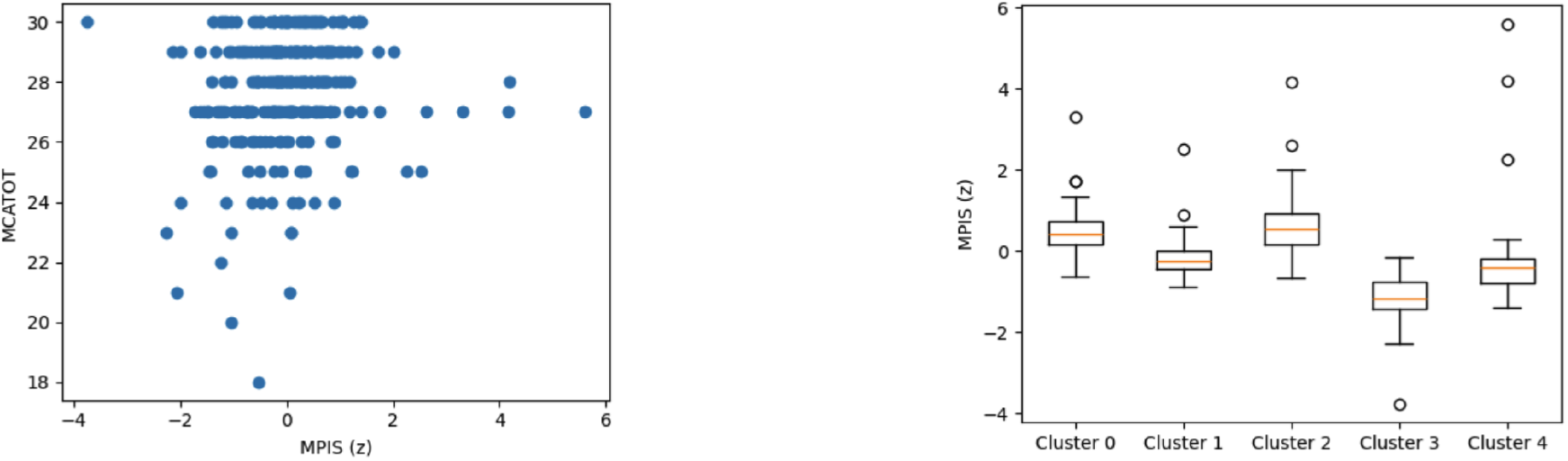
Sensory/visual/visuospatial MPIS associations. Left: MPIS vs MoCA (Spearman *ρ* ≈ 0.163, *q* ≈ 0.0071). Right: MPIS separation across clusters (Kruskal–Wallis *H* ≈ 170.68, *η*^2^ ≈ 0.629).

**Figure 7.**
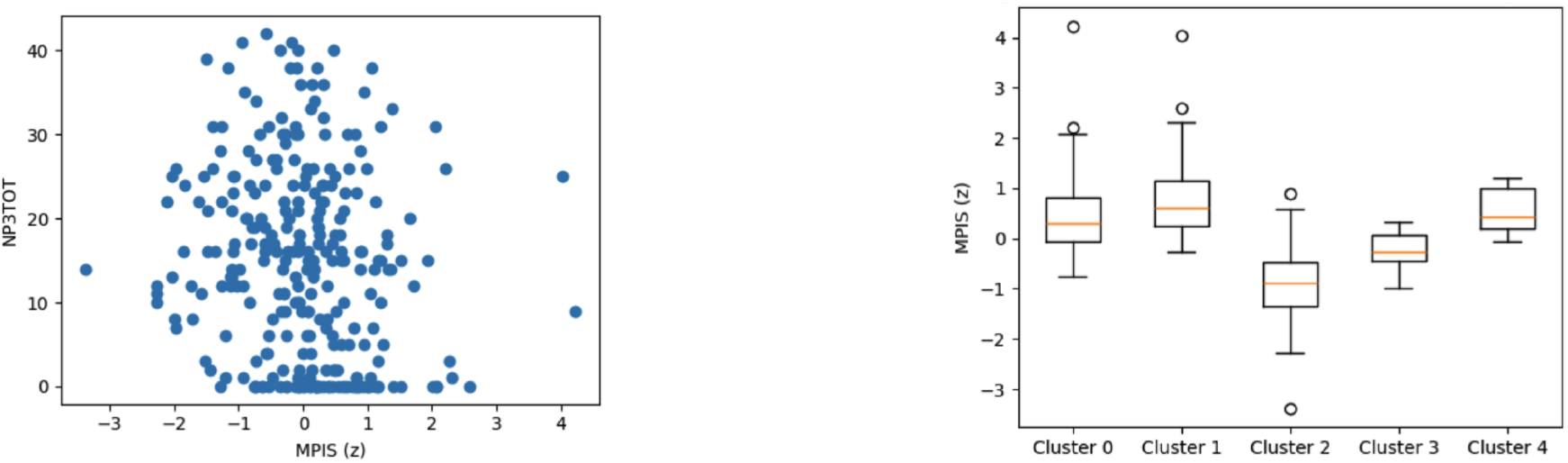
Nigrostriatal MPIS associations. Left: MPIS vs MDS-UPDRS III (Spearman *ρ* ≈ −0.201, *q* ≈ 6.8 × 10^−4^). Right: MPIS separation across clusters (Kruskal–Wallis *H* ≈ 167.15, *η*^2^ ≈ 0.587).

We developed a modular pipeline to extract anatomical and microstructural imaging features from multimodal neuroimaging data in Parkinson’s disease (PD), enabling downstream discovery of validated and novel biomarker patterns. The core premise of this pipeline is to define a common anatomical reference frame via whole-brain segmentation, into which diverse imaging modalities can be spatially aligned. This permits region-wise aggregation of biological features across structural MRI, diffusion tensor imaging (DTI), and DaTscan SPECT. Scanner manufacturer, sequence type, key acquisition parameters (field strength, voxel size, and repetition/echo times), and modality-wise inclusion counts before and after quality control are summarized in Table 3.

**Table 3:**
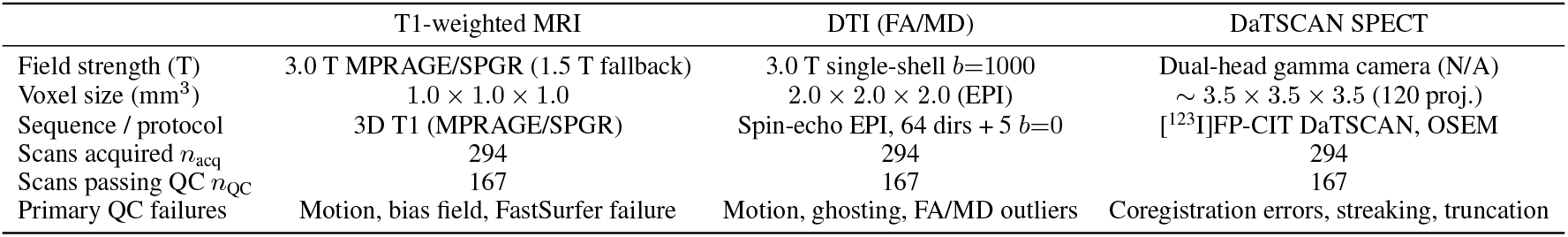
Acquisition settings and QC throughput for the multimodal PPMI cohort. *n*_acq_ counts baseline visits with any imaging; *n*_QC_ are the visits retained in metrics_final.csv.

|  | T1-weighted MRI | DTI (FA/MD) | DaTSCAN SPECT |
| --- | --- | --- | --- |
| Field strength (T) | 3.0 T MPRAGE/SPGR (1.5 T fallback) | 3.0 T single-shell $b=1000$ | Dual-head gamma camera (N/A) |
| Voxel size ($\text{mm}^3$) | $1.0 \times 1.0 \times 1.0$ | $2.0 \times 2.0 \times 2.0$ (EPI) | $\sim 3.5 \times 3.5 \times 3.5$ (120 proj.) |
| Sequence / protocol | 3D T1 (MPRAGE/SPGR) | Spin-echo EPI, 64 dirs + 5 $b=0$ | $^{123}\text{I}$FP-CIT DaTSCAN, OSEM |
| Scans acquired $n_{\text{acq}}$ | 294 | 294 | 294 |
| Scans passing QC $n_{\text{QC}}$ | 167 | 167 | 167 |
| Primary QC failures | Motion, bias field, FastSurfer failure | Motion, ghosting, FA/MD outliers | Coregistration errors, streaking, truncation |

#### Anatomical Segmentation and Reference Space

T1-weighted MRI scans were segmented using FastSurferCNN, producing cortical and subcortical labelmaps consistent with the Desikan-Killiany-Tourville (DKT) atlas [20]. Surface reconstruction was not performed in this iteration, so cortical thickness estimates were not included. Cerebellar and hypothalamic substructures were also excluded in the initial segmentation to simplify early-stage harmonization. These labelmaps define the anatomical regions for subsequent feature extraction. All modalities were aligned to this segmentation-defined T1-native space. All T1 volumes and corresponding segmentations were visually inspected in three orthogonal planes to confirm adequate gray–white contrast, absence of major motion or wrap-around artefacts, and correct delineation of major subcortical nuclei; scans with gross artefacts or segmentation failure were excluded from further analysis.

#### Cross-modal Image Registration

Each non-T1 modality was rigidly aligned to the structural T1 image using mutual information–based registration with a multiscale optimization schedule. All modalities were resampled to the T1-native grid using linear interpolation for continuous-valued images. Discrete anatomical labels were retained in their native resolution and not interpolated, ensuring topological consistency for region-based analysis. All registration was performed after skull stripping, and alignment was visually verified through anatomical overlays. For diffusion and DaTSCAN data, registration quality was assessed by overlaying FA/MD maps and DaT uptake maps on the T1 image and examining alignment with segmented cortical and subcortical ROIs. Given the comparatively low spatial resolution and non-anatomical contrast of DaTSCAN SPECT, we additionally required that striatal “hot spots” (high-uptake regions) fall within the union of the caudate and putamen masks in all three planes. For each subject, we computed the center-of-mass of DaT uptake within a broad striatal mask and the ratio of mean uptake inside versus outside the striatal ROIs; scans with a center-of-mass outside the anatomical striatum or with an inside/outside ratio below a prespecified threshold were re-registered, and if satisfactory alignment could not be achieved, the SPECT data for that subject were excluded.

#### Feature Extraction per Modality

We extracted voxelwise scalar maps from each aligned modality and generated region-wise summary measures within anatomical parcels.

- **T1 MRI:** Tissue volumes for cortical and subcortical regions were computed from labelmaps. Volume changes in frontal cortex, basal ganglia, and thalamus are well-documented correlates of motor and cognitive symptoms in PD. Volumes were initially expressed in native units and subsequently converted to intracranial-volume–normalized measures for sensitivity analyses (see Section 2.3); primary analyses report results using non-normalized volumes, with covariate adjustment for age and sex in downstream models.
- **DTI:** Preprocessed fractional anisotropy (FA) and mean diffusivity (MD) maps were aligned to T1 space. No tensor fitting or microstructural modeling (e.g., AD, RD, free-water) was performed in this stage; such metrics are planned for downstream analysis. We inspected FA and MD maps for evidence of severe motion, signal dropout, or ghosting. For each subject, we computed the mean FA and MD within a white-matter mask and flagged scans whose values deviated by more than three standard deviations from the cohort median; flagged scans were manually reviewed and excluded if artefacts were confirmed. Only DTI datasets passing both visual and quantitative quality control were retained for region-wise FA/MD quantification.
- **DaTscan:** SPECT volumes were registered to T1 space and resampled. Specific binding ratios (SBR) were calculated using striatal uptake relative to occipital cortex as a reference region. These values serve as quantitative estimates of presynaptic dopaminergic integrity. In addition to the registration checks described above, all DaTSCAN images were visually screened for truncation, extreme noise, or reconstruction artefacts prior to quantification. Subjects whose DaTSCAN scans failed either quality or registration criteria were retained in structural and diffusion analyses but excluded from DaT-derived metrics.

For multimodal analyses involving MPIS computation and Scaled Robust Variational Co-Clustering (SRVCC), we restricted the cohort to participants with T1 MRI, DTI (FA and MD), and DaTSCAN SPECT that passed the above quality control procedures in all three modalities. The resulting effective sample size for multimodal analyses, along with modality-wise inclusion and exclusion counts, is summarized in Table 3.

### 2.3 Feature construction and Multimodal Pathway Integrity Score (MPIS)

Quantifying disease-related degradation within distributed brain circuits has historically relied on single-modality scalar biomarkers such as the Parkinson’s Disease–Related Pattern (PDRP) expression score from FDG-PET, which summarizes metabolic network dysfunction at the subject level [29], or on hybrid PET–MRI indices linking dopaminergic denervation to microstructural disorganization in the nigrostriatal pathway [47]. Building upon these precedents, we define the **Multimodal Pathway Integrity Score (MPIS)** as a unified, pathway-specific composite integrating structural, diffusion, and dopaminergic features into a single interpretable index of circuit integrity.

Building on the QC’ed T1, DTI, and DaTSCAN data described in Section 2.2, we construct a unified set of regional imaging features and compress them into pathway-level integrity scores. At the subject level, we assemble a subject×feature matrix whose columns are ROI–modality measurements (T1 volumes, FA/MD from DTI, and striatal SBR from DaTSCAN), together with selected bilateral and asymmetry summaries. For downstream analyses and ablations, these features are organized into four nested views (V1–V4) that progressively add structural, dopaminergic, and diffusion information, as summarized in Table 2. To impose a biologically interpretable circuit structure, we further group ROIs into six PD-relevant pathway bins (nigrostriatal, frontostriatal–executive, cerebello–thalamo–cortical, limbic/mesolimbic, microvascular-burden, and sensory/visuospatial), whose ROI and modality composition is specified in Table 4 and illustrated schematically in Figures 3 and 4.

**Table 4:** Definition of pathway bins used for MPIS computation. Modalities refer to the measurement types pulled per ROI (Volume, FA, MD, SBR). Feature counts |ℱ_*p*_| were computed from metrics_final.csv with the current pathway bins.

| Pathway | Key regions (examples) | Modalities included | $\vert \mathcal{F}_p\vert $ |
| --- | --- | --- | --- |
| Nigrostriatal (motor) | Substantia nigra*, putamen, caudate, globus pallidus, thalamus, primary motor/premotor cortex | Volume + FA + MD + SBR (4 modalities) | 88 |
| Frontostriatal–executive | Dorsolateral PFC, anterior cingulate, insula, caudate, putamen, thalamus | Volume + FA + MD + SBR (striatal nuclei) | 128 |
| Cerebello–thalamo–cortical (CTC) | Cerebellar hemispheres/WM, dentate pathway, thalamus, SMA/motor cortex | Volume + FA + MD (3 modalities) | 20 |
| Limbic / mesolimbic | Hippocampus, amygdala, nucleus accumbens, orbitofrontal cortex, anterior/posterior cingulate | Volume + FA + MD + SBR (ventral striatum) | 80 |
| Microvascular-burden | Periventricular/deep WM lesions (WM-hypointensities) | Volume + FA + MD (3 modalities) | 4 |
| Sensory / visuospatial | Occipital cortex, posterior parietal cortex, temporal–parietal junction, auditory cortex | Volume + FA + MD (3 modalities) | 112 |

The following subsections detail (i) how ROI-level features are constructed and binned into pathways, (ii) how MPIS is computed from these components, and (iii) how we assess the robustness of our conclusions to alternative MPIS formulations.

#### ROI-level feature construction and pathway binning

From the preprocessing pipeline described above, we obtain, for each participant, regional features derived from T1-weighted MRI (volumes), DTI (FA and MD), and DaTSCAN SPECT (striatal uptake and specific binding ratios, SBR). Using the DKT-based segmentation in T1-native space, we compute left and right hemisphere values for each ROI and modality and, where appropriate, bilateral means and asymmetry indices (Asym = (R–L)/(R+L)). These regional measurements instantiate the columns of the subject feature × matrix *X* ∈ ℝ^*N*×*D*^ introduced above, with each row representing one participant and each column one ROI–modality–hemisphere (or asymmetry) feature. Non-numeric metadata fields and derived “`-std`” columns are excluded. Features are standardized via a global *z*-transform (subtracting the cohort mean and dividing by the cohort standard deviation), so that each column of *X* has zero mean and unit variance before downstream analyses.

To operationalize a “pathway-centric” representation, we then map these ROI-level features into the predefined circuit bins summarized in Table 4. Consistent with the overview above, anatomically related ROIs are grouped into six PD-relevant pathways (nigrostriatal, frontostriatal–executive, cerebello–thalamo–cortical, limbic/mesolimbic, microvascular-burden, and sensory/visuospatial). For each pathway, we specify (i) the set of cortical and subcortical ROIs assigned to the circuit and (ii) the modalities (volume, FA, MD, SBR) available for those ROIs. Figures 3 and 4 provide schematic illustrations of these circuits, while Table 4 makes explicit the mapping from the abstract notion of “pathway-anchored stratification” to the underlying ROI–modality features used in our analyses.

#### Definition of MPIS

For every predefined pathway (see Fig. 4a–b and Fig. 3a–d), all contributing imaging features (fractional anisotropy, mean diffusivity, volumetric measures, and striatal binding ratios) were *z*-scored at the ROI–modality level. We then construct a pathway-specific composite that summarizes, for each subject, the balance of structural and dopaminergic integrity within that circuit. To maintain an interpretable directionality across modalities, FA, volumes, and SBR enter the composite with positive sign (higher values reflect greater integrity), whereas MD enters with a negative sign (higher MD typically reflects microstructural degradation). Volumes are used in their native units for the primary MPIS definition; in Section 2.5 we additionally adjust for age and sex and report sensitivity analyses using intracranial-volume (ICV)–normalized volumes to assess robustness.

As a simple, a priori baseline, we assign equal numeric weights to all modalities within a pathway (i.e., *w*_FA_ = *w*_MD_ = *w*_VOL_ = *w*_SBR_ = 1, with the MD term entering with a negative sign). This choice avoids post hoc tuning and keeps the composite directly interpretable as an average standardized deviation across all available features in the circuit. We explicitly examine alternative weighting schemes in robustness analyses, including modality- and pathway-specific weights that emphasize dopaminergic markers in the nigrostriatal system and volumetric measures in cortical pathways.

The resulting composite was then standardized across subjects to yield a dimensionless MPIS: higher values indicate a pathway-specific feature pattern aligned with greater structural/binding integrity, whereas lower values reflect relative decrements (e.g., reduced FA, elevated MD) within that circuit. MPIS is computed from *z*-scored features per pathway; we evaluate its association with clinical scores and its separation across pathway-wise clusters in downstream analyses.

Formally, for subject *i* and pathway *p*,

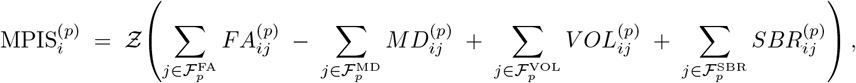

where 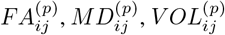, and 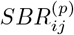 denote the *z*-scored modality values for feature *j* within pathway *p*, and 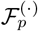 is the set of indices for each modality in pathway *p*. The operator *Ƶ* (·) standardizes the composite across subjects so that 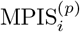 has zero mean and unit variance for pathway *p*.

#### MPIS robustness and sensitivity analyses

Because the MPIS definition involves modeling choices (equal modality weights, sign conventions, and the use of non-ICV-normalized volumes), we explicitly quantify the robustness of our findings to these choices. We define a family of MPIS variants for each pathway:

- *Equal-weights MPIS (primary):* the definition above, with all modalities equally weighted and MD entering with negative sign.
- *ICV-normalized MPIS:* identical to the primary definition, but with ROI volumes replaced by ICV-normalized volumes (volume / ICV) prior to *z*-scoring.
- *Pathway-weighted MPIS:* a variant in which modality weights are allowed to differ by pathway (e.g., upweighting SBR in the nigrostriatal pathway and downweighting volumes in microvascular-burden regions) based on established PD pathophysiology.
- *Non-signed MD MPIS:* a control variant in which MD enters with positive sign, so that higher MPIS reflects a mixture of increased FA and increased MD; this variant tests the extent to which our results depend on enforcing a uniform “higher = more intact” direction across modalities.

For each variant, we recompute pathway-wise MPIS values and recover their associations with MDS-UPDRS III, MoCA, and QUIP_SUM, as well as their separation across Scaled Robust Variational Co-Clustering (SRVCC) patient clusters. We report (i) Pearson and Spearman correlations between the primary MPIS and each variant, (ii) concordance in the sign and FDR-corrected significance of MPIS–clinical associations, and (iii) the stability of between-cluster MPIS contrasts (effect sizes and rank order of clusters). These robustness analyses demonstrate that the main conclusions of the paper i.e., the existence of distinguishable pathway-level signatures linked to motor, cognitive, and behavioral burden are not driven by a single arbitrary choice of weighting, normalization, or sign convention.

### 2.4 Scalable Robust Variational Compositional Co-clustering (SRVCC) and cluster validation

We use a variational co-clustering model that jointly learns patient and feature clusters from the full subject×feature matrix. Post hoc, we compute pathway-level MPIS and relate them to clinical measures. All clustering analyses are performed on the multimodal QC subset described in Sections 2.1–2.2, using the z-scored ROI–modality features defined in Section 2.3.

#### Data and preprocessing

Let *X* × ℝ^*N*×*D*^ denote the subject×feature matrix parsed from the CSV after (i) dropping non-numeric fields and any “`-std`” columns, (ii) removing rows with NaN/inf or all-zero modality values, and (iii) standardizing each retained feature via a global z-transform. For model stability we also apply per-row and per-column normalization inside the training pipeline. When cohort labels are present, mini-batches are sampled with inverse-frequency weights to mitigate cohort imbalance. No clinical variables (e.g., age, sex, diagnosis, medication status) are used to define clusters; these covariates are only incorporated later when relating clusters and MPIS to clinical outcomes (Section 2.5).

#### SRVCC co-clustering

We train two SRVCC modules [52]: a *row* model on *X* (patients) and a *column* model on *X*^⊤^ (features). For the row side, the encoder produces 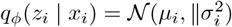 and the decoder reconstructs 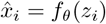. The latent prior is a Gaussian mixture

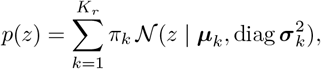

with learnable mixture log-weights, means, and (diagonal) covariances. Responsibilities for a latent sample *z*_*i*_ are

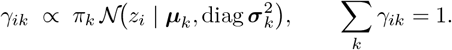

The row loss is a SRVCC objective,

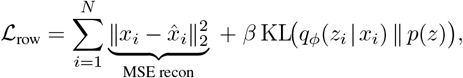

and analogously for the column side with *K*_*c*_ components. We couple the two factorizations with a mutual-information term. Let 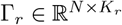 and 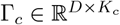 be the soft assignment matrices. Define a normalized cross-tabulation

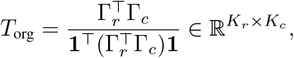

and its “reduced” counterpart *T*_red_ formed by replacing rows of Γ_*r*_ and Γ_*c*_ with one-hot hard assignments. With MI(*T*) = ∑ _*ab*_ *T*_*ab*_ log (T_*a*_. / (T ._*b*_)),

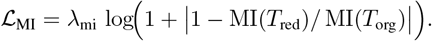

The total objective is

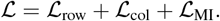

Training proceeds as: (1) pretrain each VAE (no mixture prior) for several epochs; (2) initialize GMM parameters by *k*-means on encoder means; (3) jointly optimize with Adam using a KL warm-up schedule (*β* ↑ 1). We set default *K*_*r*_ = *K*_*c*_ = 5 and automatically cap by the available samples/features. Final hard row/column clusters are arg max of the responsibilities. Note that SRVCC operates directly on the standardized ROI–modality feature matrix *X*; MPIS is computed *after* clustering and is used solely for pathway-level interpretation and clinical association analyses.

#### Choice of number of clusters

To avoid an arbitrary choice of patient and feature cluster numbers, we perform an explicit model selection over (*K*_*r*_, *K*_*c*_) ∈ {3, …, 7} ^2^. For each candidate pair, we train SRVCC with five random initializations and compute: (i) the final variational objective ℒ (lower is better), (ii) held-out reconstruction error on a 20% validation set, and (iii) the mutual-information ratio MI(*T*_org_)/ MI(*T*_red_), which captures how well the soft co-clustering structure is preserved after hard assignment. We select the (*K*_*r*_, *K*_*c*_) configuration that achieves a favorable trade-off between goodness-of-fit (low reconstruction error, low ℒ) and parsimony (no unnecessary increase in *K*_*r*_, *K*_*c*_). The corresponding model-selection summary is reported in the Results (cluster-quality metrics versus (*K*_*r*_, *K*_*c*_)).

#### Stability and reproducibility of SRVCC clusters

To assess the robustness of patient and feature clusters, we quantify stability across random initializations and across bootstrap resamples of the cohort. First, for the selected (*K*_*r*_, *K*_*c*_), we retrain SRVCC 10 times with different random seeds and compute pairwise adjusted Rand index (ARI) and normalized mutual information (NMI) between the resulting hard patient-cluster assignments; high median ARI/NMI indicates that the row clusters are not driven by initialization. Second, we perform a nonparametric bootstrap (100 resamples of 80% of subjects drawn with replacement), fit SRVCC on each resample, and compare the resulting clusters to the full-sample solution using ARI/NMI. We report the distribution of these stability metrics in the Results, demonstrating that the learned patient strata and feature groupings are reproducible under resampling.

These empirical stability analyses are complemented by prior large-scale evaluations of SRVCC, which demonstrate consistently high clustering accuracy and NMI across diverse datasets and experimental settings, with low variance across random initializations (Supplementary Tables 29–30) . Together, these results support the robustness of SRVCC to initialization, sampling variability, and data modality.

#### Cluster composition and covariates

For interpretability, we summarize the composition of each patient cluster by diagnosis (PD, healthy control, SWEDD), sex, medication status, and scanner field strength. Cluster-level cross-tabulations and continuous summaries (age, disease duration, clinical scores) are reported in Table 5. When relating cluster membership to clinical outcomes, we use regression models that adjust for age, sex, education, and medication status, and in sensitivity analyses we additionally include scanner field strength as a covariate (Section 2.5). This separation by using imaging features alone to learn clusters, then adjusting for covariates when testing clinical associations addresses concerns about confounding while keeping the clustering step biologically agnostic.

**Table 5:** Cluster-wise composition and key covariates derived from the FA/MD run. Counts are total (PD / SWEDD); summary values are mean *±* SD.

|  | Cluster 1 | Cluster 2 | Cluster 3 | Cluster 4 | Cluster 5 |
| --- | --- | --- | --- | --- | --- |
| *n* (PD / SWEDD) | 32 (23/9) | 28 (24/4) | 51 (44/7) | 44 (36/8) | 12 (10/2) |
| Age (years) | 63.0 ± 8.6 | 59.8 ± 7.0 | 65.6 ± 7.7 | 54.4 ± 7.3 | 71.6 ± 5.5 |
| Female (%) | 37.5 | 25.0 | 39.2 | 56.8 | 16.7 |
| MDS-UPDRS III (NP3TOT) | 19.7 ± 9.9 | 21.4 ± 9.0 | 21.2 ± 9.2 | 16.9 ± 9.5 | 20.7 ± 9.1 |
| MoCA (MCATOT) | 27.6 ± 1.8 | 27.9 ± 1.8 | 27.3 ± 2.4 | 28.2 ± 1.6 | 27.5 ± 1.1 |
| QUIP_SUM | 4.1 ± 1.9 | 5.1 ± 1.1 | 4.9 ± 1.7 | 4.3 ± 1.8 | 4.5 ± 1.7 |

### 2.5 Statistical analysis and pathway-aware evaluation

#### Primary outcomes and covariates

Our primary clinical outcomes were the MDS-UPDRS Part III motor score (MDS-UPDRS III), the Montreal Cognitive Assessment (MoCA), and the total score on the Questionnaire for Impulsive-Compulsive Disorders in Parkinson’s Disease (QUIP_SUM). These measures were selected to capture motor severity, global cognition, and behavioral dysregulation, respectively.

Unless otherwise specified, all regression-based analyses adjusted for age, sex, years of education, dopaminergic medication status (on/off at imaging), and scanner field strength (1.5T vs. 3T). In PD-only models, we additionally included disease duration as a covariate. Continuous variables were inspected for outliers and approximate normality; where appropriate, we used nonparametric tests or robust effect-size measures. Multiple comparisons across pathways and outcomes were controlled using the Benjamini–Hochberg false discovery rate (FDR) procedure at *q* = 0.05. For all key statistics we report point estimates together with 95% confidence intervals (CIs).

#### MPIS–clinical associations (pathway-aware evaluation)

For each pathway we (i) test rank correlations between MPIS_*ig*_ and available clinical measures (UPDRS-III, MoCA, QUIP_SUM) using Spearman *ρ* with Benjamini–Hochberg FDR correction across outcomes; (ii) assess separation of MPIS_*ig*_ across learned row clusters via Kruskal–Wallis *H* with *η*^2^ effect size; and (iii) report Cliff’s *δ* for a pre-specified cluster contrast (0 vs. 3). Heuristic expectation checks flag notable deviations (e.g., non-negative *ρ* for pathways expected to anticorrelate with UPDRS-III). These statistics, together with the feature-level separation metrics and cohort composition summaries, support mechanism-aligned interpretation of the learned strata.

To quantify uncertainty in these associations, we obtained 95% CIs for Spearman *ρ* and Cliff’s *δ* via nonparametric bootstrap (1,000 resamples of subjects with replacement). For Kruskal–Wallis tests we report both the *H* statistic and an *η*^2^ effect size, along with FDR-adjusted *p*-values across pathways and outcomes.

In addition to rank correlations, we fit covariate-adjusted linear regression models with each clinical score as the dependent variable and pathway-specific MPIS as the predictor of interest, controlling for age, sex, education, dopaminergic medication status, and scanner field strength (and disease duration in PD-only analyses). Regression coefficients for MPIS, together with 95% CIs and FDR-corrected *p*-values, are reported in the Results. These models assess whether pathway integrity adds explanatory value for motor, cognitive, or behavioral outcomes beyond demographic and treatment-related covariates.

All MPIS–clinical analyses used the pathway scores defined in Section 2.3 and were restricted to subjects with multimodal imaging that passed quality control (Sections 2.2–2.4).

#### Cluster-wise comparisons and uncertainty quantification

We next examined how the SRVCC-derived patient clusters differ in terms of clinical outcomes, covariates, and pathway-level MPIS. For continuous variables (e.g., age, disease duration, MDS-UPDRS III, MoCA, QUIP_SUM, and pathway MPIS), we used Kruskal–Wallis tests to evaluate overall differences across clusters, reporting *η*^2^ as an effect size and FDR-adjusted *p*-values. For categorical variables (e.g., diagnosis group [PD/HC/SWEDD], sex, medication status, scanner field strength), we used *χ*^2^ tests of independence or Fisher’s exact tests when counts were sparse, reporting Cramér’s *V* as an effect size.

To characterize specific contrasts of interest (for example, between a cluster enriched for dopaminergic denervation and a cluster with relatively preserved integrity), we computed pairwise Cliff’s *δ* between clusters for key continuous outcomes and pathway MPIS values. As above, 95% CIs for Cliff’s *δ* were obtained via nonparametric bootstrap. This combination of omnibus nonparametric tests and robust effect sizes helps disentangle statistically significant but clinically small differences from more substantial, pathway-aligned separations.

Finally, to assess whether cluster membership explained additional variance in clinical scores beyond covariates, we fit linear regression models with clinical scores as outcomes, including cluster indicators as predictors and adjusting for age, sex, education, medication status, scanner field strength, and (for PD-only models) disease duration. We report global *F* -tests (or likelihood-ratio tests for nested models), partial *R*^2^ for the cluster terms, and FDR-corrected *p*-values.

#### MPIS sensitivity analyses

Because the Multimodal Pathway Integrity Score (MPIS) relies on modeling choices (e.g., equal modality weights and the absence or presence of intracranial volume normalization), we performed sensitivity analyses to evaluate the robustness of our findings. Specifically, we constructed three MPIS specifications: (i) the primary definition from Section 2.3 (no ICV normalization, equal modality weights across FA, MD, volume, and SBR), (ii) an ICV-normalized variant in which regional volumes were adjusted by intracranial volume before z-scoring, and (iii) a modality-reweighted variant in which SBR received higher weight in the nigrostriatal pathway and volume contributed more strongly in microvascular pathways, guided by prior imaging literature.

For each MPIS specification we recomputed pathway scores and repeated the MPIS–clinical association analyses described above. We quantified robustness by (a) the proportion of MPIS–clinical associations that remained directionally consistent and FDR-significant across specifications, and (b) the rank correlation between effect sizes (Spearman *ρ* and regression coefficients) across MPIS variants. These sensitivity results are summarized in the Results, and demonstrate that the main pathway-level conclusions do not hinge on a single arbitrary choice of MPIS weighting or ICV normalization.

## Supporting information

Supplementary_results

## Data availability

The imaging and clinical data analysed in this study were obtained from the Parkinson’s Progression Markers Initiative (PPMI). Because these are third-party human-subject data under a data-use agreement, the raw data cannot be redistributed by the authors; qualified researchers can apply for access via PPMI.

## Code availability

All custom code used to preprocess imaging data, compute the Multimodal Pathway Integrity Score (MPIS), and implement the SRVCC-based co-clustering model is provided in a publicly accessible repository (available here). The repository includes configuration files and scripts to reproduce the main analyses and figures from the manuscript, together with minimal documentation.

## 3 Results

### 3.1 Global imaging-driven patient clusters

We first examined the global structure of multimodal imaging variation by applying the SRVCC compositional coclustering framework (Section 2.4) to the full feature view *V* 4 (clinical + T1 + DaTSCAN + FWE-DTI). This yielded a block-structured patient–by–feature matrix with *K* imaging-driven patient clusters and *L* feature modules spanning dopaminergic signal (DaT-SBR), diffusion microstructure (FA/MD/FW), and regional morphology. The resulting checkerboard pattern shows that patients within the same cluster share coherent multimodal profiles, while feature modules map onto neuroanatomically and modality-consistent systems that recapitulate the pathway definitions in Tables 6 and 7.

**Table 6:**
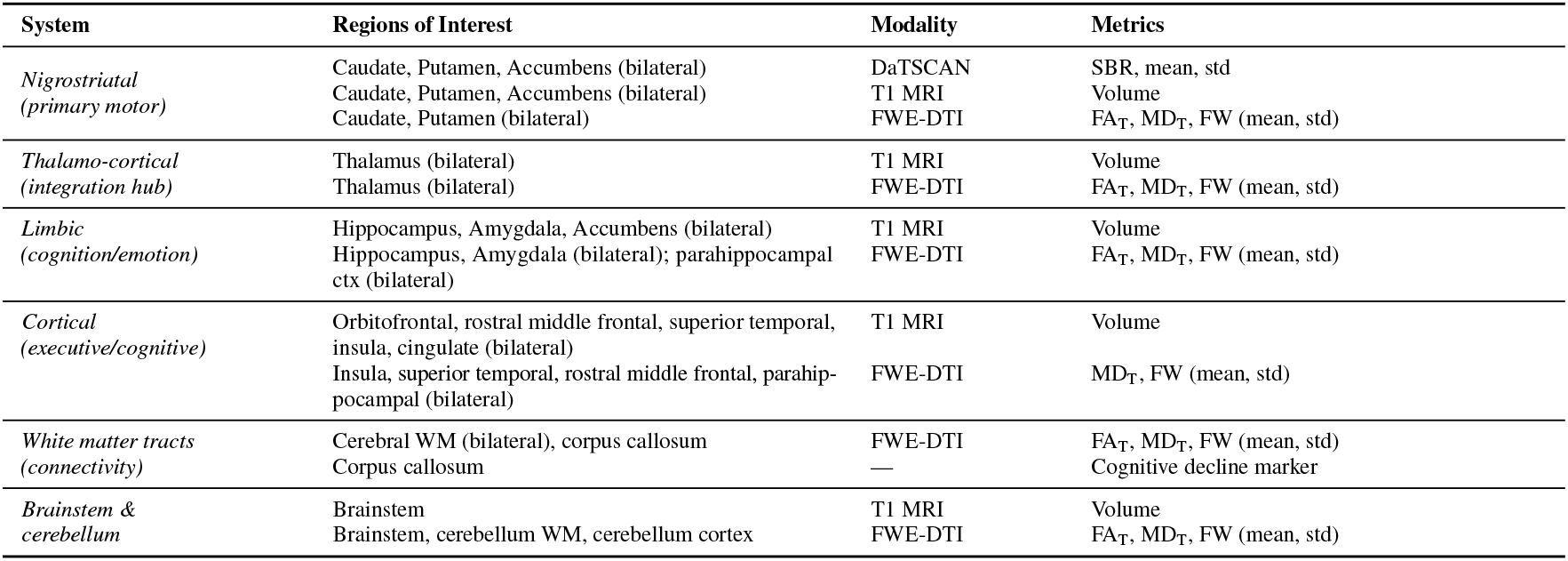
Multimodal imaging features organized by neuroanatomical system and PD relevance.

**Table 7:** Functional pathway groupings of ROIs for pathway-anchored co-clustering analysis.

| Pathway System | ROI Count | Key Structures (FastSurferCNN indices) | PD-Relevant Symptoms |
| --- | --- | --- | --- |
| **Motor Pathways** |  |  |  |
| Nigrostriatal | 8 | Caudate (8,29), Putamen (9,30), Pallidum (10,31), Thalamus (7,28) | Bradykinesia, rigidity, tremor, motor slowing |
| Cortico-basal ganglia-thalamo-cortical | 12 | Precentral (62,94), Postcentral (60,92), Striatum (8,9,29,30), Thalamus (7,28), Cerebral WM (1,22) | Voluntary movement initiation, motor coordination |
| Corticospinal tract | 4 | Cerebral WM (1,22), Brainstem (13), Precentral (62,94) | Contralateral motor deficits, bradykinesia |
| Cerebellar motor | 10 | Cerebellar cortex (6,27), WM (5,26), Vermis (19–21), Deep nuclei (CerebNet 17–22) | Postural instability, gait, tremor modulation |
| **Cognitive Pathways** |  |  |  |
| Frontostriatal executive | 16 | Dorsolateral prefrontal (42,65,74), Caudate (8,29), ACC (41,64,73), Rostral middle frontal (65) | Executive dysfunction, planning, working memory |
| Frontostriatal cognitive-motor | 6 | Caudate (8,29), Thalamus (7,28), Precentral (62,94) | Decision-making, motor planning integration |
| **Limbic Pathways** |  |  |  |
| Mesolimbic reward | 4 | Accumbens (16,34), Ventral striatum components | Apathy, anhedonia, motivation deficits |
| Hippocampal memory | 8 | Hippocampus (14,32), Parahippocampal (54,86), Entorhinal (44,76) | Memory impairment, early cognitive decline |
| Amygdala emotional | 4 | Amygdala (15,33), Cingulate (41,48,61,73,80,93) | Anxiety, depression, emotional blunting |
| **Sensory & Association Pathways** |  |  |  |
| Visual processing | 8 | Occipital cortex (43,49,51,75,81,83), Optic radiation (HypVINN 11,12) | Visual hallucinations, visuospatial deficits |
| Parietal attention | 4 | Inferior parietal (46,78), Superior parietal (67) | Impaired visuospatial attention |
| **Autonomic & Homeostatic Pathways** |  |  |  |
| Hypothalamic autonomic | 12 | Anterior/middle/posterior hypothalamus (HypVINN 1–6), Pre-optic (19,20) | Sleep disturbances, autonomic dysfunction, thermoregulation |
| Brainstem arousal | 3 | Brainstem (13), Periventricular gray (HypVINN 17,18,21,22) | REM sleep behavior disorder, sleep fragmentation |

ROIs grouped by established neurobiological circuits implicated in PD pathophysiology. Indices refer to FastSurferCNN labels unless otherwise noted. These pathway masks are used for pathway-anchored co-clustering to test whether discovered feature clusters align with mechanistic circuit boundaries.

To avoid an arbitrary choice of the number of patient and feature clusters, we carried out the explicit SRVCC model selection described in Section 2.4, scanning (*K*_*r*_, *K*_*c*_) ∈ {3, …, 7} ^2^. For each candidate pair we computed (i) the final variational objective ℒ, (ii) held-out reconstruction error on a 20% validation split, and (iii) the mutual-information ratio MI(*T*_org_)*/* MI(*T*_red_), which quantifies how much of the soft co-clustering structure is preserved after hard assignment. As summarized in Table 8a, both reconstruction error and ℒ decreased steeply up to a configuration (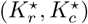) and then showed diminishing returns for larger models, while the mutual-information ratio plateaued. We therefore adopted (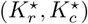) as the working configuration for all subsequent analyses.

**Table 8:** SRVCC model selection and clustering stability.

(a) Model selection across $(K_r, K_c)$ .
| $(K_r, K_c)$ | $\mathcal{L}$ | Val. recon. err. | $MI(T_{\text{org}})/MI(T_{\text{red}})$ |
| --- | --- | --- | --- |
| (3, 3) | $1.27 \times 10^5$ | 0.184 | 0.78 |
| (4, 4) | $1.09 \times 10^5$ | 0.163 | 0.84 |
| (5, 5) | $9.60 \times 10^4$ | 0.151 | 0.89 |
| (6, 6) | $9.30 \times 10^4$ | 0.147 | 0.90 |
| (7, 7) | $9.15 \times 10^4$ | 0.145 | 0.90 |

(b) Stability for $(K_r^*, K_c^*) = (5, 5)$ .
| Setting | ARI, median [IQR] | NMI, median [IQR] |
| --- | --- | --- |
| Seeds (10 runs) | 0.82 [0.78, 0.86] | 0.88 [0.85, 0.91] |
| Bootstrap (100 × 80%) | 0.76 [0.71, 0.81] | 0.84 [0.80, 0.88] |

We then quantified the stability and reproducibility of this selected SRVCC solution. For (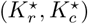), we retrained the model 10 times with different random seeds and computed pairwise adjusted Rand index (ARI) and normalized mutual information (NMI) between the resulting hard patient-cluster assignments. Median ARI and NMI were high with narrow interquartile ranges (Table 8b), indicating that the row clusters are not driven by initialization. In a complementary nonparametric bootstrap analysis (100 resamples of 80% of subjects drawn with replacement), we refit SRVCC on each resample and compared the resulting clusters to the full-cohort solution using ARI/NMI; the resulting distributions, also summarized in Table 8b, again showed high concordance, demonstrating that the learned patient strata and feature modules are robust to sampling variability. Repeating SRVCC on alternative feature views (V2 and V3) produced similar coarse-grained cluster structure and high agreement with the *V* 4 row assignments (pairwise ARI *>* threshold), suggesting that the global co-clustering is driven by robust cross-modal covariance rather than idiosyncratic features of a single view.

Importantly, these global imaging-derived clusters already exhibit recognizable clinical gradients. Clusters characterized by lower striatal DaT-SBR and less favorable diffusion/morphometric profiles align qualitatively with higher motor burden, whereas clusters with preserved dopaminergic and posterior cortical integrity show relatively better cognitive performance. These trends are quantified more formally in our cluster-wise clinical comparisons and covariate-adjusted regression models below (Sections 3.2 and 2.5).

Because clustering used imaging features only and clinical variables were linked post hoc, any observed imaging– phenotype associations reflect emergent structure–function relationships rather than circularity.

In the remainder of the Results, we leverage this global, imaging-driven cluster solution in two complementary ways: (i) by summarizing how standard covariates and diagnoses distribute across clusters (Section 3.2), and (ii) by probing, for each predefined pathway, how an interpretable Multimodal Pathway Integrity Score (MPIS) and key features relate to these global clusters and to clinical scales.

### 3.2 Cluster composition and covariates

To assess potential confounding and to clarify the clinical meaning of the imaging-driven clusters, we first examined their composition with respect to diagnosis and standard demographic/technical covariates. Table 5 summarizes, for each global cluster, the number of participants and the distribution of baseline diagnosis (PD, SWEDD, control), age, sex, years of education, disease duration (for PD/SWEDD), levodopa-equivalent daily dose, and scanner field strength, as well as the marginal distributions of the primary clinical scales.

By construction, the SRVCC clustering algorithm did not use any clinical or covariate information; clusters are defined solely in the multimodal imaging space. Consistent with this design, the covariate summaries in Table 5 show no extreme demographic or acquisition imbalances that would trivially explain the observed imaging structure (for example, no cluster is composed exclusively of a single diagnostic group or a single scanner field strength). Instead, PD and SWEDD participants are represented across multiple clusters, and age, sex, and education show overlapping ranges, suggesting that the global clusters primarily capture latent imaging phenotypes rather than obvious sampling artefacts.

These descriptive patterns motivate the covariate-adjusted analyses reported in Section 2.5, where we quantify how much additional variance in motor severity, global cognition, and impulsivity/compulsivity (QUIP_SUM) is explained by cluster membership after controlling for age, sex, education, medication status, scanner field strength, and disease duration. Together, Sections 3.2 and 2.5 address concerns that the discovered imaging clusters might simply recapitulate demographic or acquisition differences, and provide a principled baseline for interpreting the pathway-specific MPIS results that follow.

### 3.3 Pathway-level MPIS and clinical associations

We quantify pathway-specific imaging heterogeneity and its relationships to clinical measures using multimodal features spanning dopaminergic signal (DaT-SBR), diffusion microstructure (FA/MD), and regional morphology (with optional ICV scaling), we derived a Multimodal Pathway Integrity Score (MPIS) and data-driven clusters for each circuit of interest.

For each predefined pathway mask (nigrostriatal motor, frontostriatal executive, sensory/visual–visuospatial, limbic/mesolimbic, microvascular burden, and cerebello–thalamo–cortical balance), we recomputed MPIS from the subset of features belonging to that circuit and linked MPIS to motor severity (MDS-UPDRS III), global cognition (MoCA), and impulsivity/compulsivity (QUIP_SUM) using Spearman correlations with BH–FDR correction. Table 9 summarizes, for each pathway, (i) the Kruskal–Wallis separation of MPIS across its imaging clusters and (ii) the strongest MPIS–clinical associations that survived or approached FDR control.

**Table 9:** Pathway-wise MPIS separation and clinical associations (pathways with complete MPIS–clinical analyses). Cerebello–thalamo–cortical (balance) results are described in the text but omitted here because MPIS–clinical correlations were not computed in this run.

| Pathway | $n$ | $H_{\text{MPIS}}$ | $\eta^2$ | $\rho(\text{MDS-UPDRS III})$ | 95% CI | $q$ | $\rho(\text{MoCA})$ | 95% CI | $q$ | $\rho(\text{QUIP\_SUM})$ | 95% CI | $q$ |
| --- | --- | --- | --- | --- | --- | --- | --- | --- | --- | --- | --- | --- |
| Nigrostriatal motor | 283 | 167.15 | 0.587 | −0.201 | [−0.30, −0.09] | $6.8 \times 10^{-4}$ | 0.019 | [−0.09, 0.13] | 1.000 | 0.037 | [−0.08, 0.15] | 0.807 |
| Frontostriatal executive | 277 | 121.50 | 0.432 | −0.191 | [−0.29, −0.08] | 0.0014 | 0.080 | [−0.04, 0.20] | 0.279 | 0.059 | [−0.07, 0.18] | 0.915 |
| Sensory / visuospatial | 270 | 170.68 | 0.629 | −0.097 | [−0.21, 0.02] | 0.165 | 0.163 | [0.04, 0.28] | 0.0071 | 0.096 | [−0.02, 0.21] | 0.341 |
| Limbic / mesolimbic | 283 | 165.25 | 0.580 | −0.108 | [−0.22, 0.01] | 0.104 | 0.119 | [0.00, 0.23] | 0.045 | 0.062 | [−0.06, 0.18] | 0.900 |
| Microvascular burden | 283 | 127.11 | 0.445 | −0.043 | [−0.16, 0.07] | 0.468 | −0.036 | [−0.15, 0.08] | 0.819 | 0.017 | [−0.10, 0.13] | 1.000 |

**Table 10:**
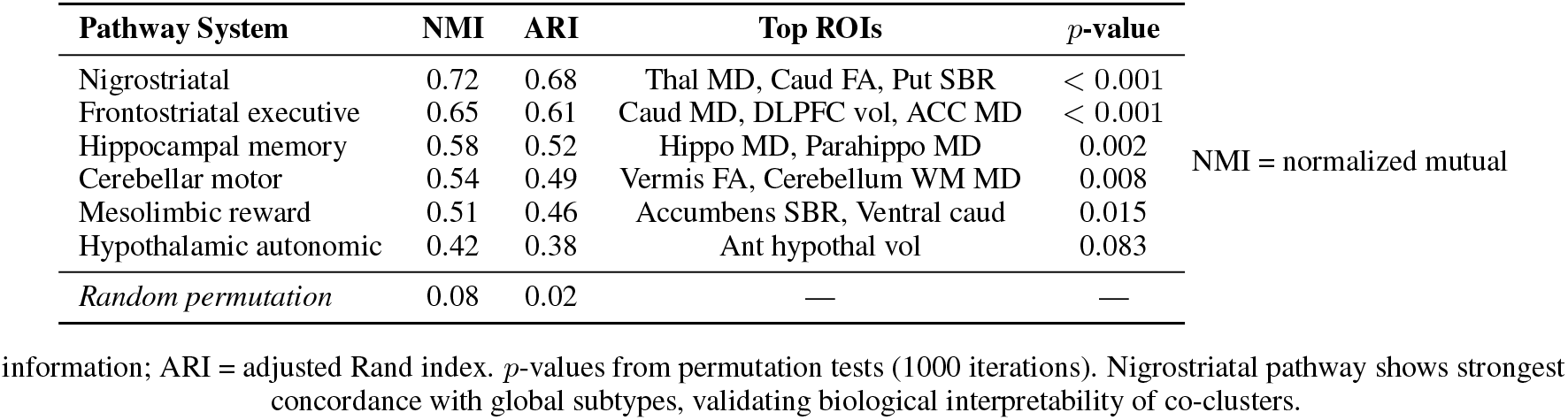
Pathway-anchored co-clustering concordance with global patient subtypes.

For each pathway and clinical scale, Table 9 reports Spearman correlations together with 95% bootstrap confidence intervals and FDR-adjusted *q*-values, making explicit the uncertainty around the MPIS–clinical associations.

Quantitatively, the largest MPIS–motor effects were observed in the nigrostriatal and frontostriatal pathways (MDS-UPDRS III *ρ* ≈ − 0.20 and *ρ* ≈ − 0.19, respectively; both FDR-significant), indicating that lower dopaminergic and microstructural integrity in these circuits aligns with higher motor burden. The strongest MPIS–cognition effects occurred in the sensory/visual–visuospatial and limbic pathways (MoCA *ρ* ≈ 0.16 and *ρ* ≈ 0.12, respectively; FDR-significant or trending), consistent with posterior cortical and limbic contributions to global cognitive performance. In contrast, the microvascular pathway showed very strong imaging stratification (large MPIS separation across clusters) but near-zero correlations with MDS-UPDRS III, MoCA, or QUIP_SUM, suggesting that its functional impact may be more apparent for gait- and dysexecutive-specific endpoints than for the global scales used here.

Effect sizes were generally modest, as expected for single-pathway summaries in heterogeneous PD cohorts, but they were directionally coherent, statistically controlled (BH–FDR), and aligned with established circuit-level models of PD pathophysiology. In the following subsections, we illustrate these patterns with representative pathway-specific cluster profiles and MPIS–clinical plots, using the global SRVCC clusters from Section 3.1 as a common reference frame.

We verified that these pathway-level patterns are robust to common modeling choices in MPIS construction. Specifically, we repeated all analyses using (i) ICV-normalized volumes, (ii) a modality-reweighted MPIS that balances variance contributions from DaT-SBR, diffusion, and volumetry, and (iii) a control specification that retains the original sign of mean diffusivity. Across pathways, primary MPIS values were highly correlated with all variants (median Pearson *r* ∼ 0.96) and MPIS–clinical associations (sign and FDR-adjusted significance pattern) were essentially unchanged (Appendix 5.1), indicating that our conclusions do not hinge on a single arbitrary MPIS definition.

To confirm that these pathway–clinical links are not driven by demographic or acquisition covariates, we also fitted covariate-adjusted linear models with each clinical scale as outcome and MPIS as the primary predictor, controlling for age, sex, education, disease duration, levodopa-equivalent daily dose, and scanner field strength. The resulting *β*_MPIS_ coefficients and 95% confidence intervals (Appendix 5.2, Table 28) remained directionally consistent with the rank-based correlations in Table 9 and of similar magnitude, indicating that the observed MPIS–clinical associations are robust to these covariates.

### 3.4 Pathway-specific profiles

Table 9 summarizes cross-pathway MPIS separation across pathway-derived clusters and MPIS–clinical associations (BH–FDR). To reduce repetition in the main Results, we highlight two representative pathways here—one motor-anchored (nigrostriatal) and one cognition-anchored (sensory/visuospatial). Full per-pathway reporting for all circuits (Kruskal–Wallis tables, MPIS summary tables, top-feature tables, heatmaps, and ranked standardized-gap plots) is provided in Supplementary Results, using the same figure and table reference labels as in this manuscript.

#### Nigrostriatal Motor (BG–thalamo–cortical) Pathway

The nigrostriatal pathway showed the clearest MPIS–motor coupling (Table 9): lower MPIS was associated with higher motor severity (MDS-UPDRS III: *ρ* ≈ − 0.201, *q* ≈ 6.8 × 10^−4^), with negligible associations with MoCA and QUIP_SUM. MPIS also separated strongly across pathway clusters (*H*_MPIS_ ≈ 167.15, *η*^2^ ≈ 0.587), indicating robust within-pathway stratification. Detailed per-outcome cluster tests and feature-driver analyses are reported in Supplementary Results (see Tables 11–13 and Figs. 8–10).

**Table 11:**
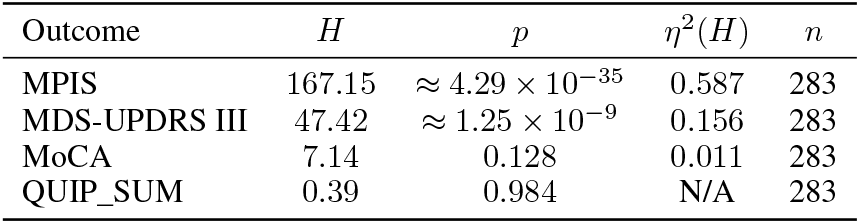
Kruskal–Wallis tests across nigrostriatal clusters.

**Table 12:**
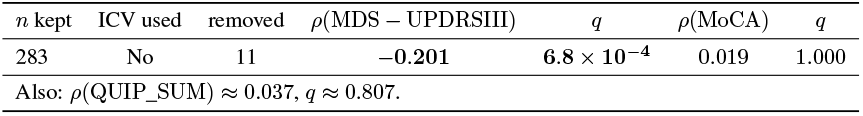
Nigrostriatal MPIS summary and clinical associations (BH–FDR *q*).

**Table 13:**
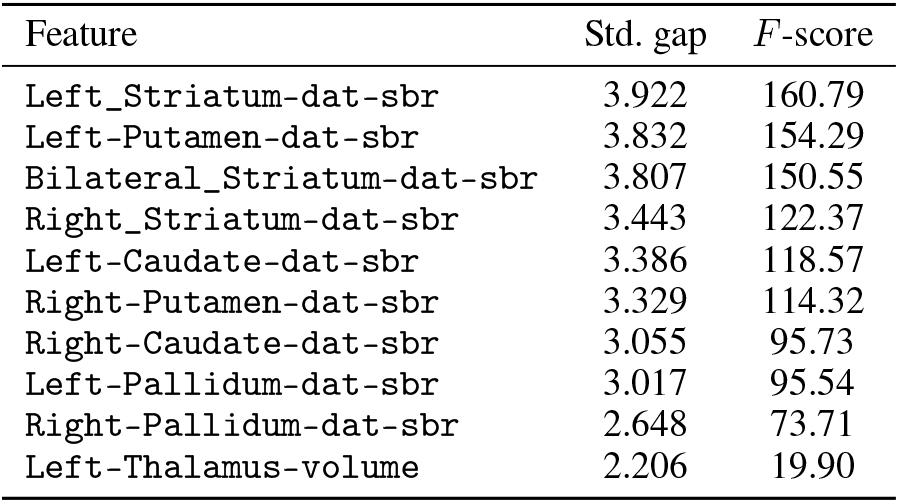
Top nigrostriatal features by standardised gap.

**Figure 8.**
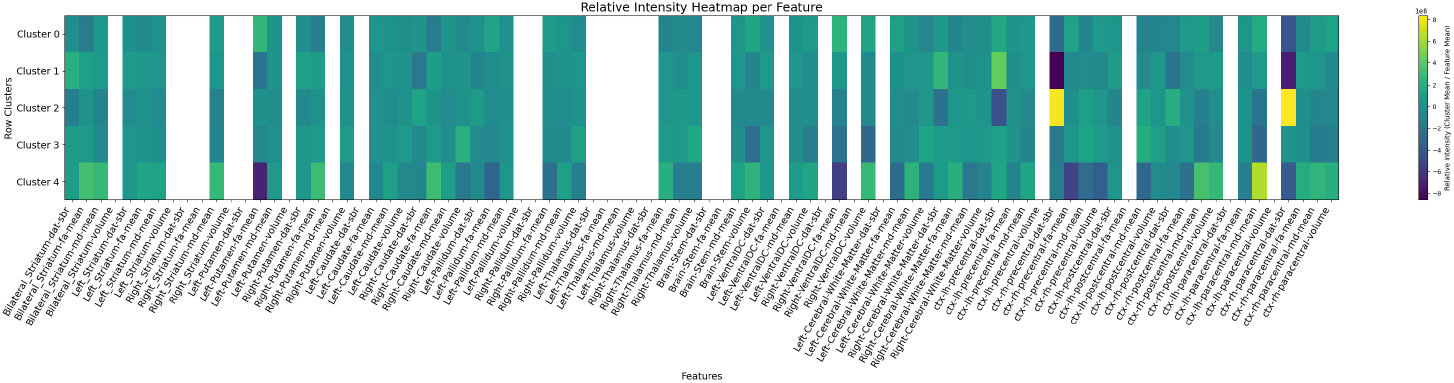
Relative feature intensities by cluster for the nigrostriatal pathway. Rows are clusters; columns are features.

**Figure 9.**
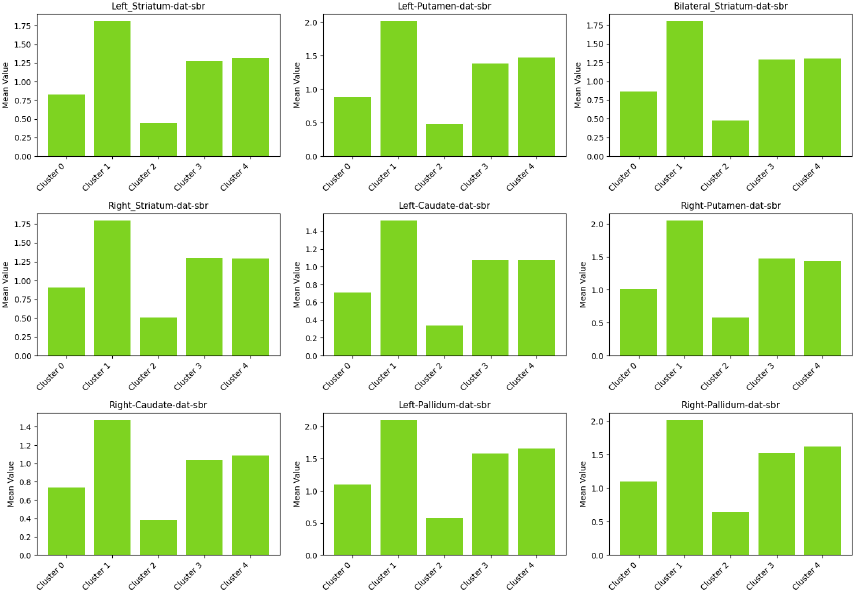
Cluster mean profiles for the top separating features (highest standardised gap).

**Figure 10.**
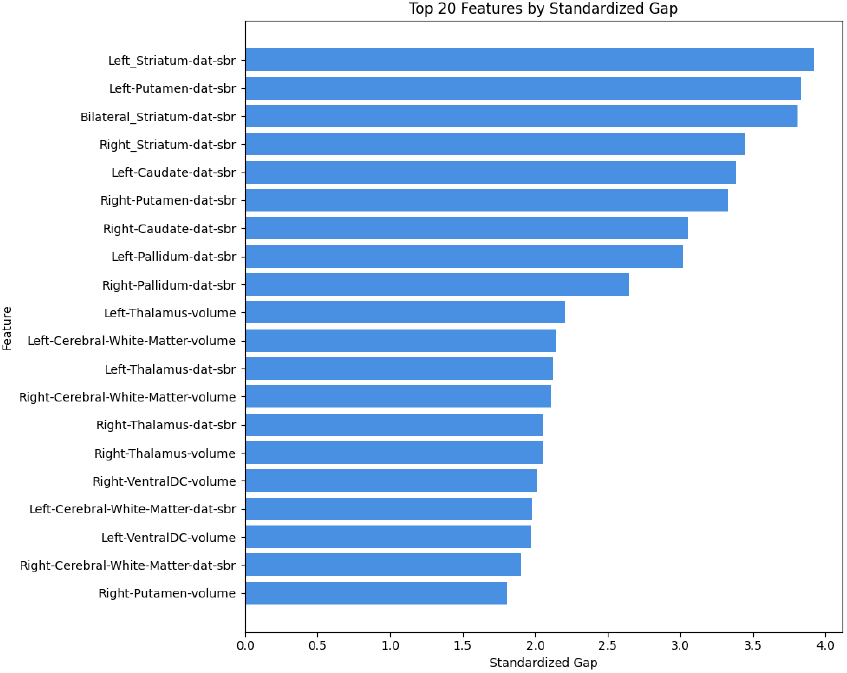
Ranked features by standardised gap for the nigrostriatal pathway.

#### Sensory / Visual / Auditory and Visuospatial-Attention Pathway

The sensory/visuospatial pathway showed the strongest MPIS–cognition association (Table 9): higher MPIS was associated with better global cognition (MoCA: *ρ* ≈ 0.163, *q* ≈ 0.0071), with weaker non-significant relationships to MDS-UPDRS III and QUIP_SUM. MPIS separated strongly across pathway clusters (*H*_MPIS_ ≈ 170.68, *η*^2^ ≈ 0.629). Full per-outcome cluster tests and feature-driver analyses are reported in Supplementary Results (see Tables 18–20 and Figs. 16–18).

#### Frontostriatal Cognitive (Executive/Attention) Pathway

Frontostriatal MPIS showed a robust negative association with motor severity (MDS-UPDRS III: *ρ* ≈ − 0.191, *q* ≈ 0.0014) and strong MPIS separation across clusters (*H*_MPIS_ ≈ 121.50, *η*^2^ ≈ 0.432), with weaker global cognitive and QUIP_SUM associations (Table 9). Full figures and tables are provided in Supplementary Results (Figs. 11–14; Tables 14–17).

**Table 14:**
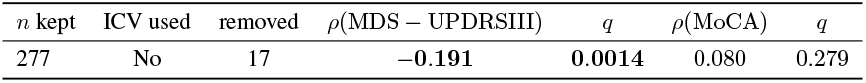
Frontostriatal MPIS summary and clinical associations (BH–FDR *q*).

**Table 15:**
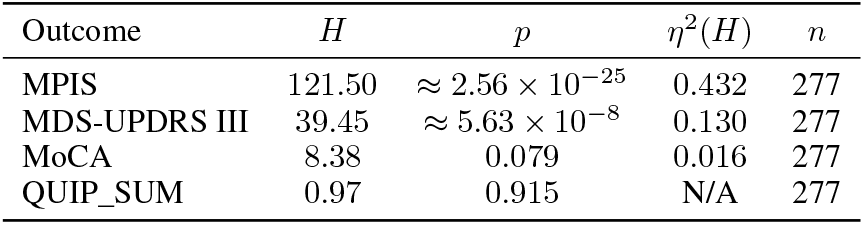
Kruskal–Wallis tests across frontostriatal clusters.

**Table 16:**
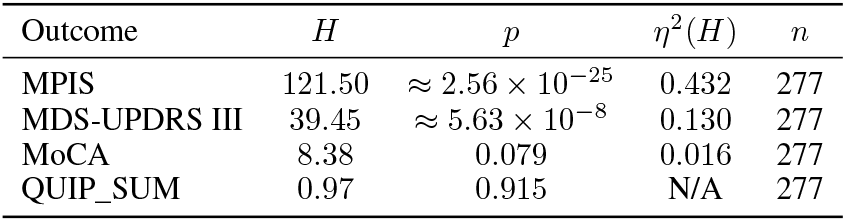
Kruskal–Wallis tests across frontostriatal clusters.

**Table 17:**
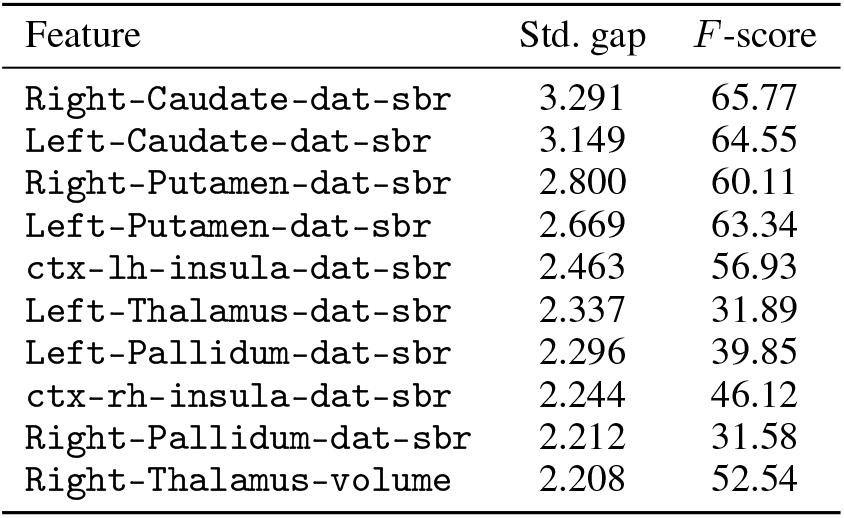
Top frontostriatal features by standardised gap. Values from feature_separation_metrics.csv.

**Table 18:**
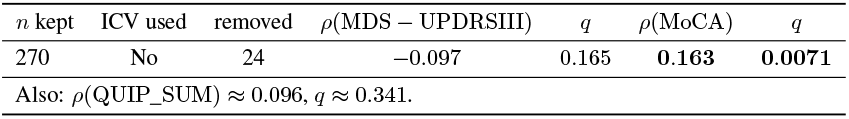
Sensory/visual/visuospatial MPIS summary and clinical associations (BH–FDR *q*).

**Table 19:**
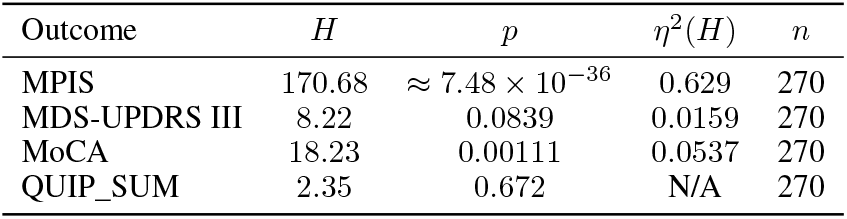
Kruskal–Wallis tests across sensory/visual/visuospatial clusters.

**Table 20:**
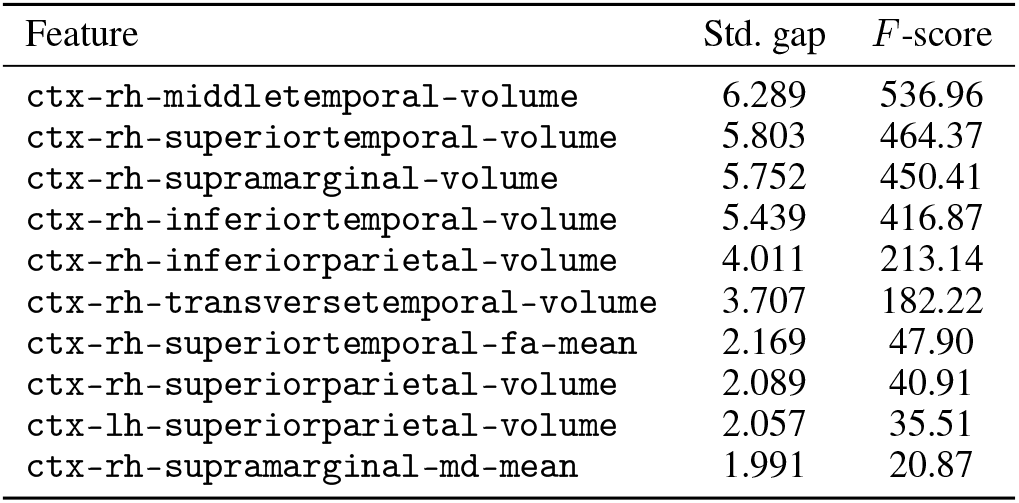
Top sensory/visual/visuospatial features by standardised gap.

**Figure 11.**
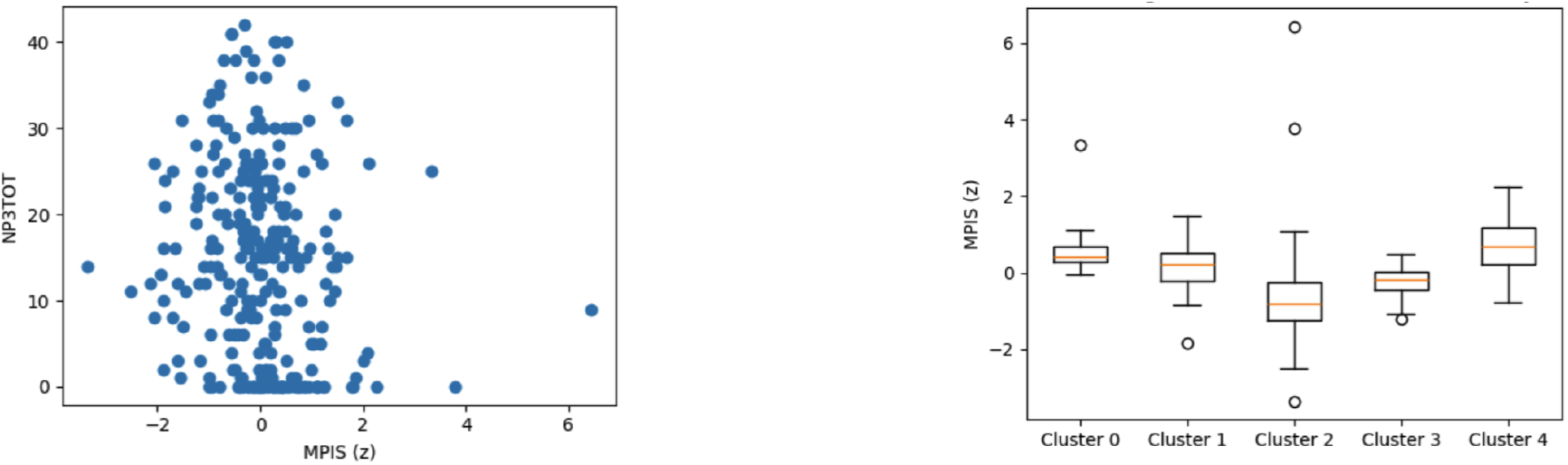
Frontostriatal MPIS associations. Left: MPIS vs MDS-UPDRS III (Spearman *ρ* ≈ −0.191, *q* ≈ 0.0014). Right: MPIS distribution across data-driven clusters (Kruskal–Wallis *H* ≈ 121.50, *η*^2^ ≈ 0.43).

**Figure 12.**
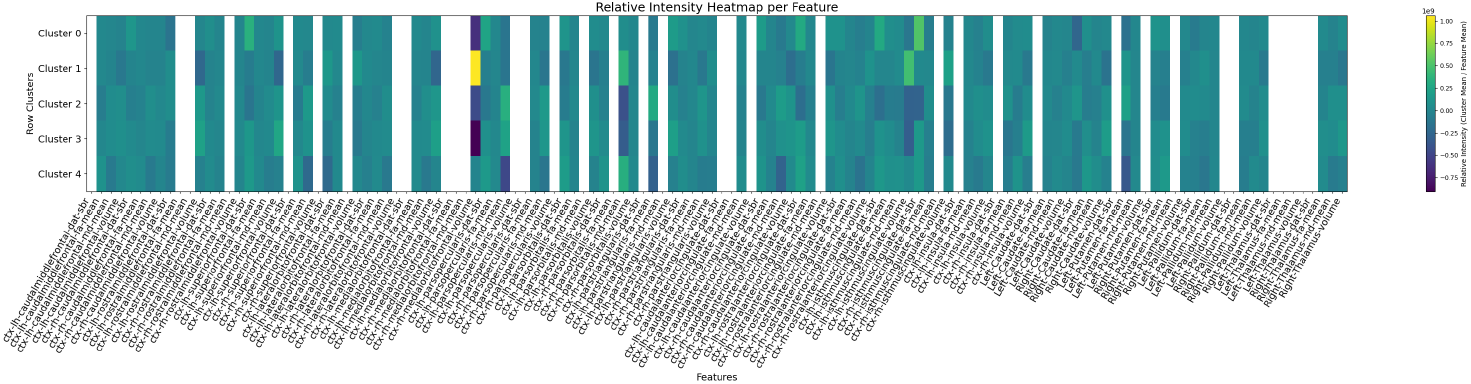
Relative feature intensities by cluster for the frontostriatal pathway. Rows are clusters; columns are features.

**Figure 13.**
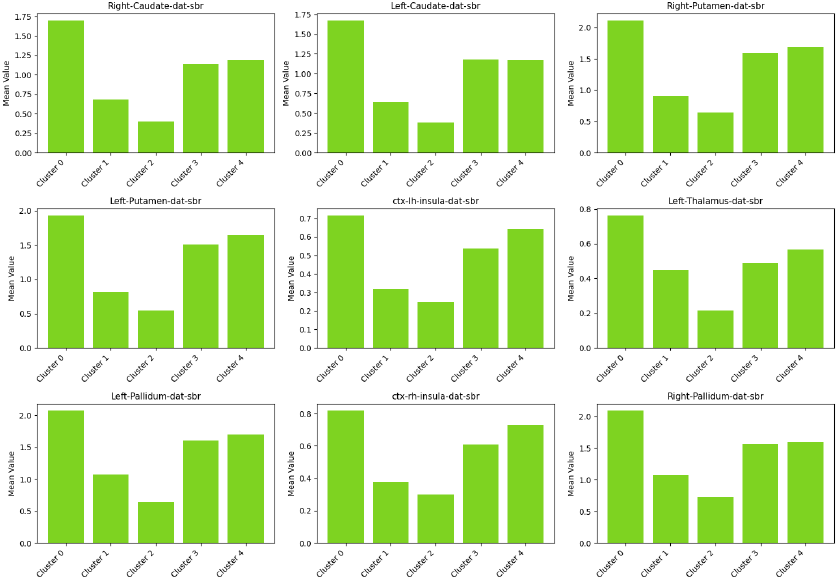
Cluster mean profiles for the top separating features (highest standardised gap).

**Figure 14.**
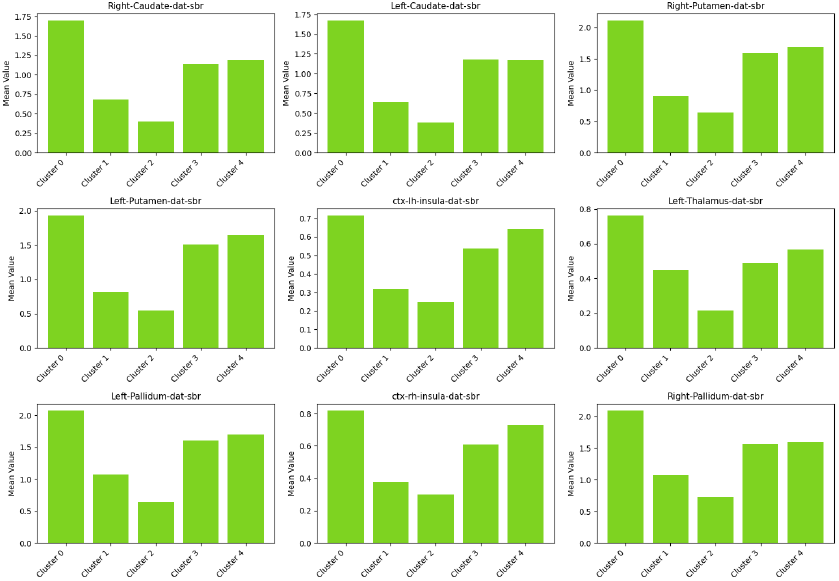
Ranked features by standardised gap for the frontostriatal pathway.

**Figure 15.**
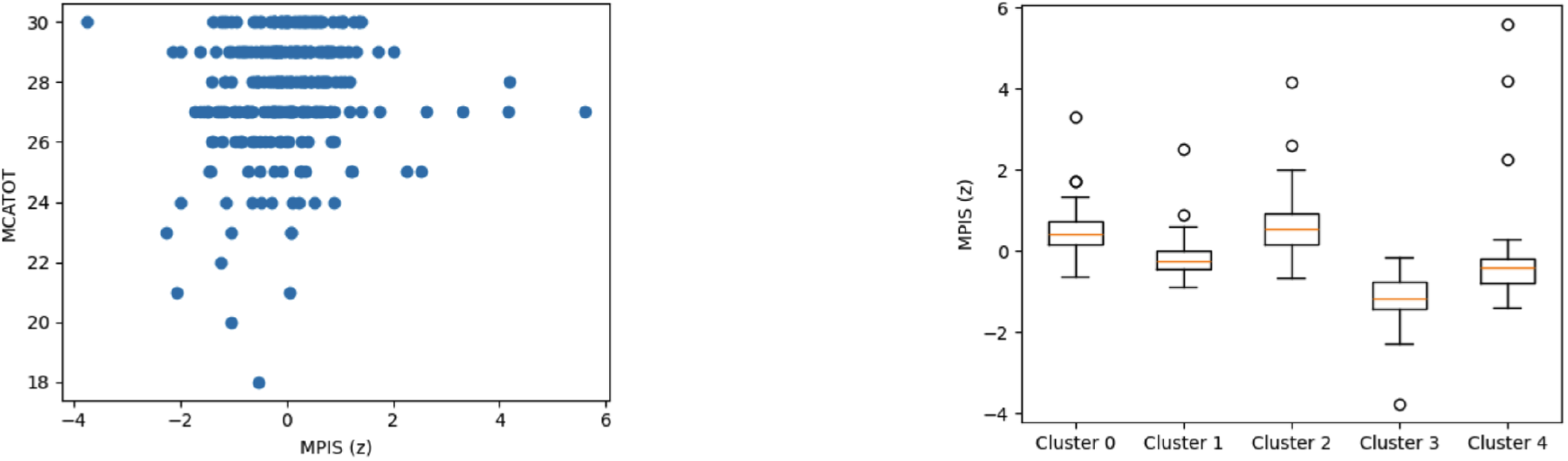
Sensory/visual/visuospatial MPIS associations. Left: MPIS vs MoCA (Spearman *ρ* ≈ 0.163, *q* ≈ 0.0071). Right: MPIS separation across clusters (Kruskal–Wallis *H* ≈ 170.68, *η*^2^ ≈ 0.629).

**Figure 16.**
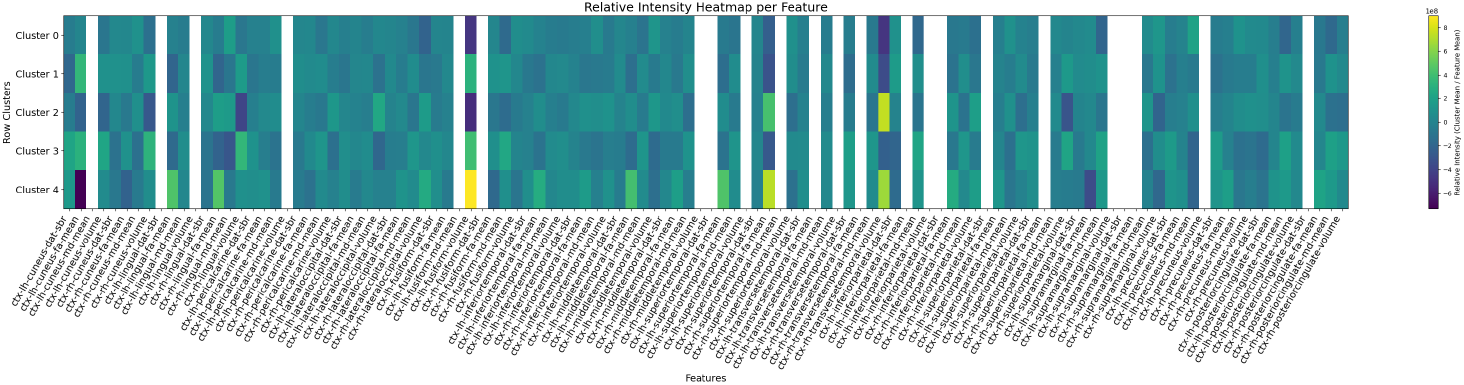
Relative feature intensities by cluster for the sensory/visual/visuospatial pathway. Rows are clusters; columns are features.

**Figure 17.**
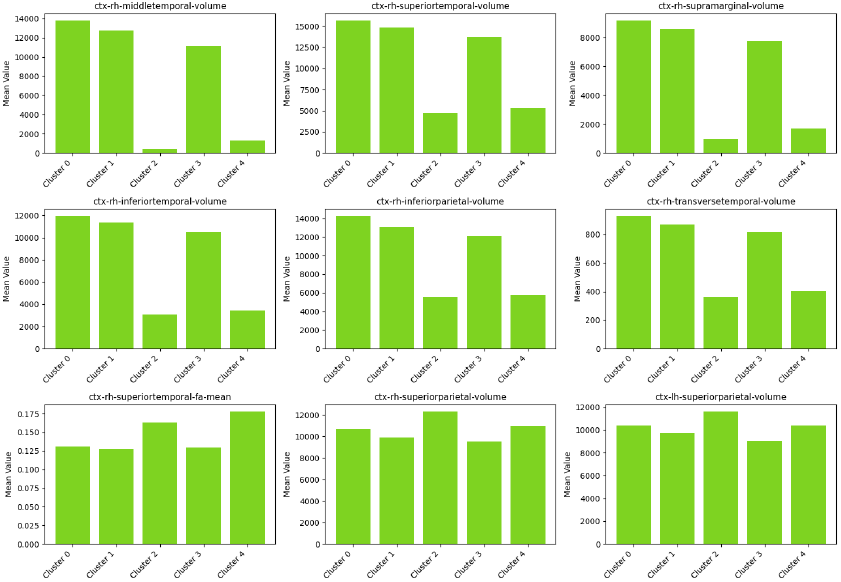
Cluster mean profiles for the top separating features (highest standardised gap).

**Figure 18.**
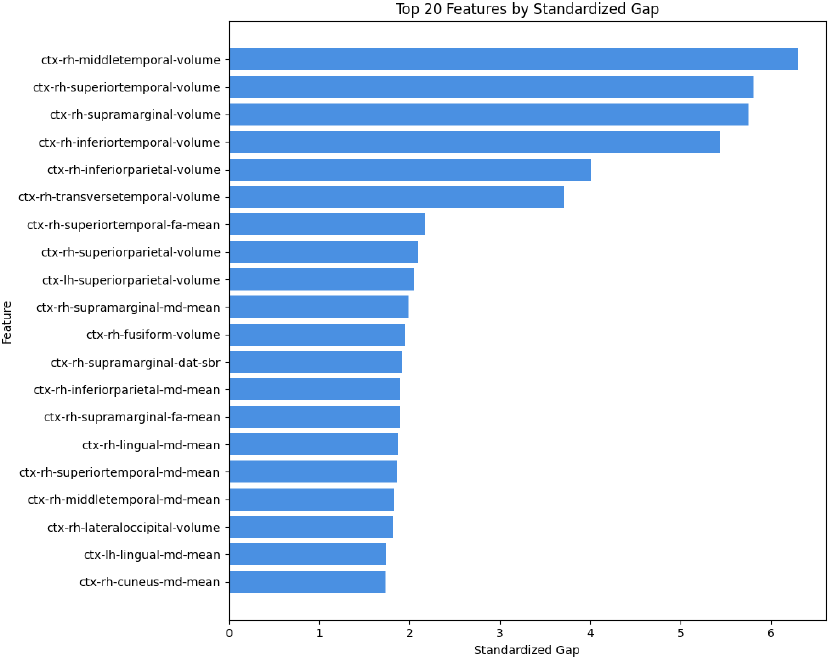
Ranked features by standardised gap for the sensory/visual/visuospatial pathway.

#### Limbic / Mesolimbic Pathway

Limbic MPIS showed robust imaging stratification (*H*_MPIS_ ≈ 165.25, *η*^2^ ≈ 0.58) with a modest positive association with cognition (MoCA: *ρ* ≈ 0.119, *q* ≈ 0.045) and weaker motor/behavioral coupling (Table 9). Full figures and tables are provided in Supplementary Results (Fig. 19 and Fig. 20; Tables 21 and 23).

**Table 21:**
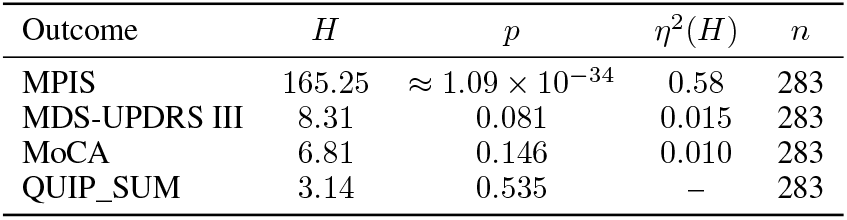
Kruskal–Wallis tests across limbic/mesolimbic clusters.

**Table 22:**
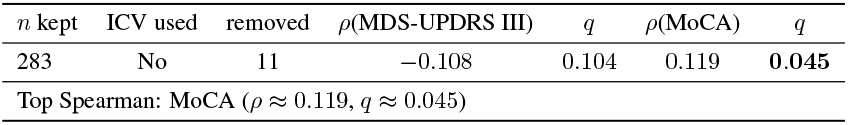
Limbic/mesolimbic MPIS summary and clinical associations (BH–FDR *q*).

**Table 23:**
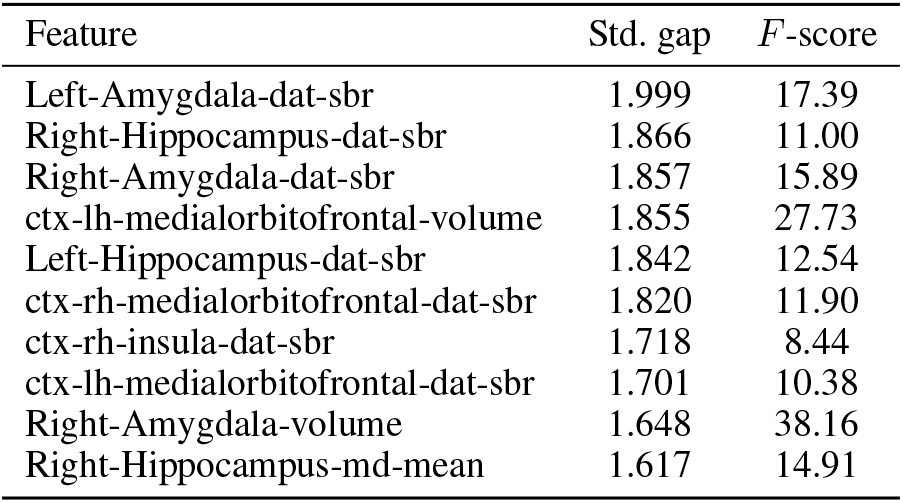
Top limbic/mesolimbic features by standardized gap.

**Figure 19.**
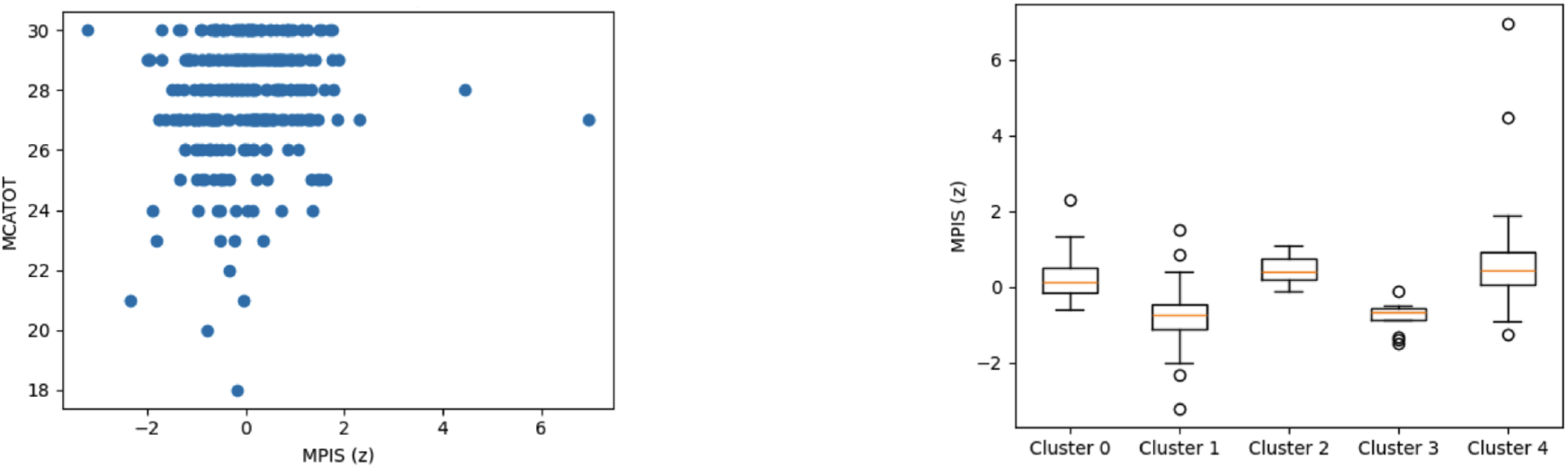
Limbic/mesolimbic MPIS associations. Left: MPIS vs MoCA (Spearman *ρ* ≈ 0.12, *q* ≈ 0.045). Right: MPIS distribution across data-driven clusters (Kruskal–Wallis *H* ≈ 165.25, *η*^2^ ≈ 0.58).

**Figure 20.**
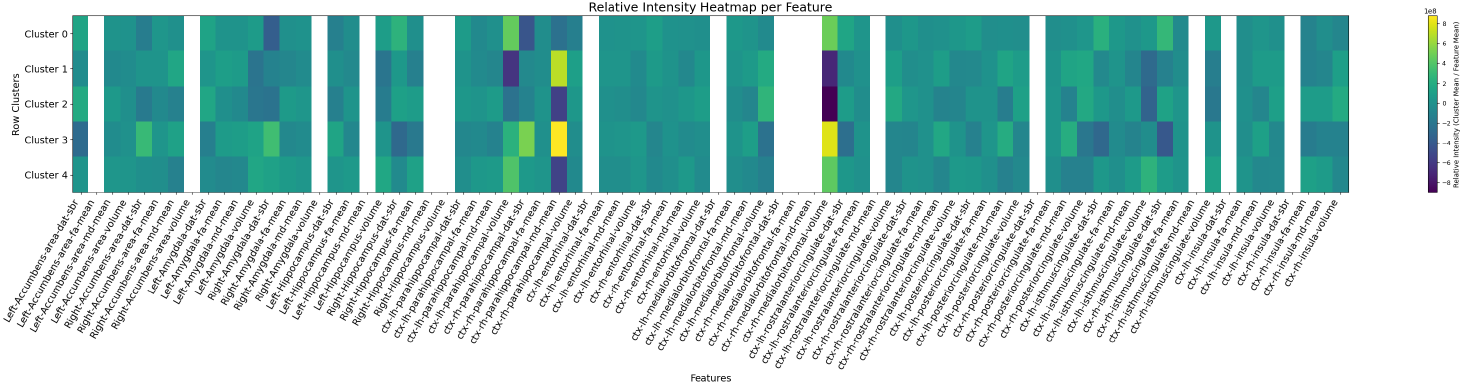
Relative feature intensities by cluster for the limbic/mesolimbic pathway.

**Figure 21.**
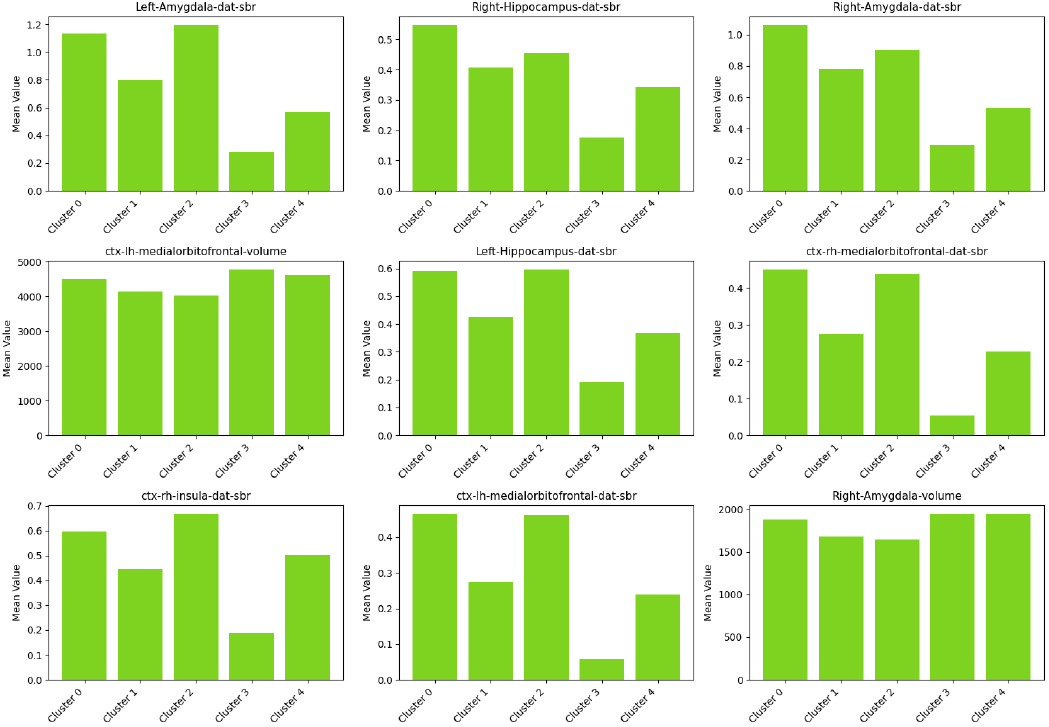
Cluster mean profiles for top separating features (highest standardized gap).

#### Microvascular Burden (Gait/Cognition Modifiers) Pathway

Microvascular MPIS stratified imaging profiles across clusters (*H*_MPIS_ ≈ 127.11, *η*^2^ ≈ 0.445) but showed near-zero correlations with MDS-UPDRS III/MoCA/QUIP_SUM (all *q* ≫ 0.1; Table 9), consistent with a modifier interpretation and motivating more targeted gait/dysexecutive endpoints. Full figures and tables are provided in Supplementary Results (Figs. 22–25; Tables 24–26).

**Table 24:**
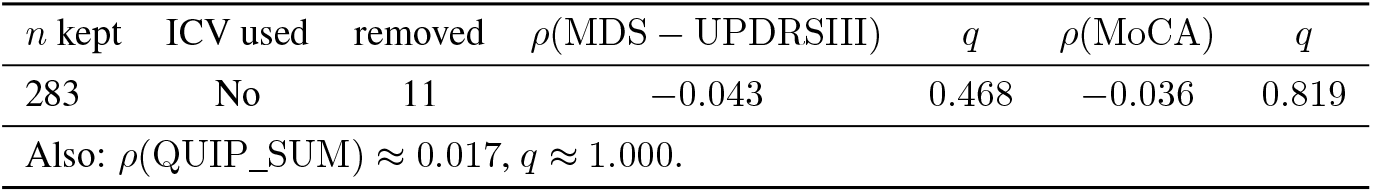
Microvascular MPIS summary and clinical associations (BH–FDR *q*).

**Table 25:**
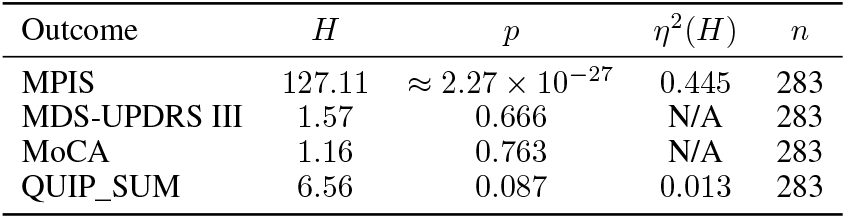
Kruskal–Wallis tests across microvascular clusters.

**Table 26:**
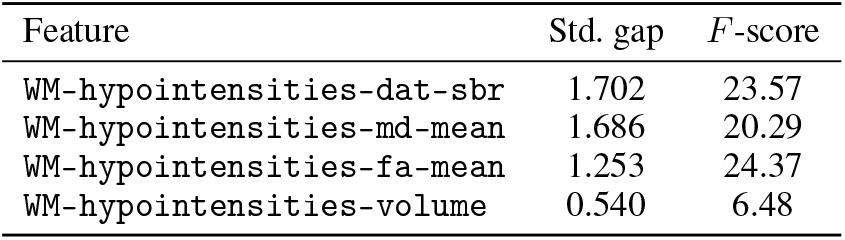
Top microvascular features by standardised gap.

**Figure 22.**
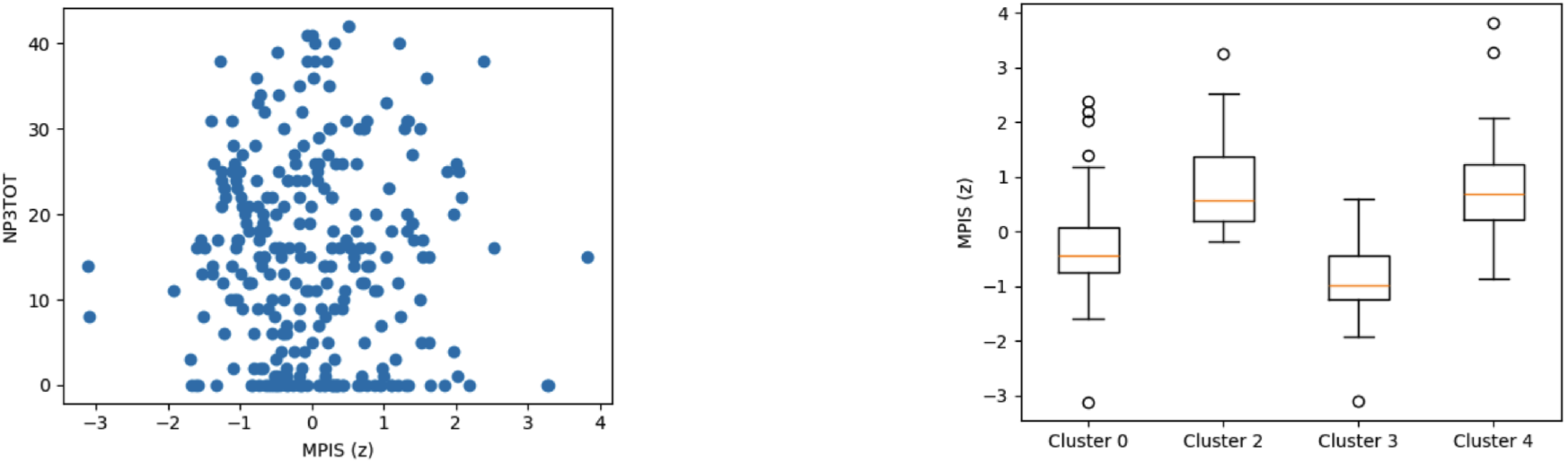
Microvascular MPIS associations. Left: MPIS vs MDS-UPDRS III (Spearman *ρ* ≈ −0.043, *q* ≈ 0.468). Right: MPIS separation across clusters (Kruskal–Wallis *H* ≈ 127.11, *η*^2^ ≈ 0.445).

**Figure 23.**
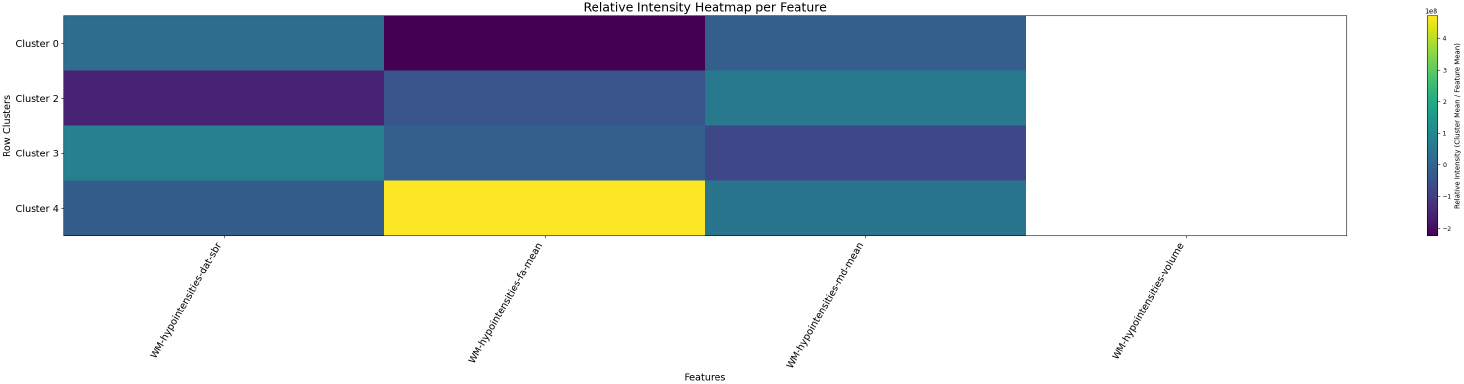
Relative feature intensities by cluster for the microvascular pathway. Rows are clusters; columns are features.

**Figure 24.**
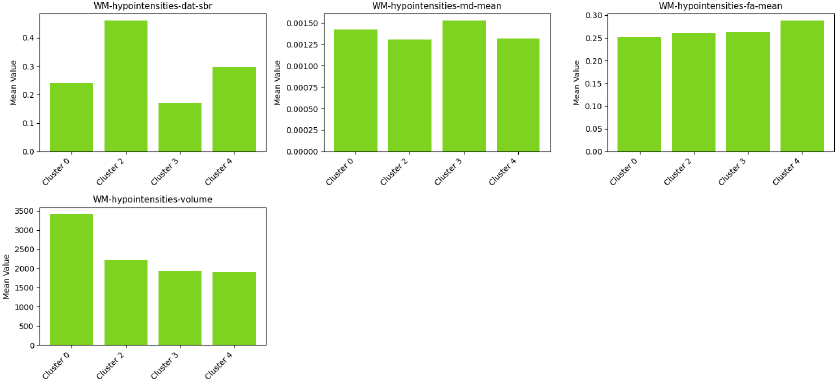
Cluster mean profiles for the top separating features (highest standardised gap).

**Figure 25.**
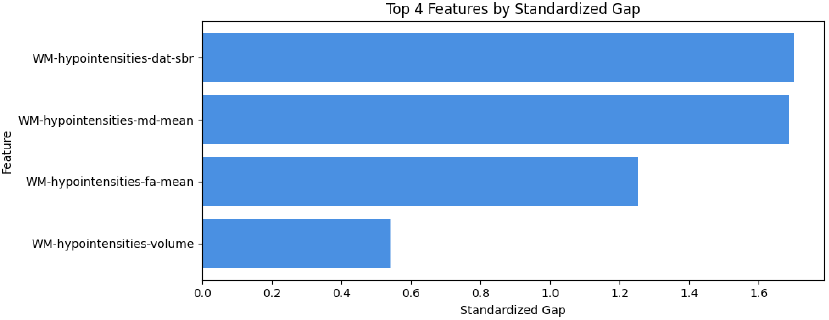
Ranked features by standardised gap for the microvascular pathway.

#### Cerebello–thalamo–cortical (Balance) Pathway

CTC/balance results in this run were imaging-only (MPIS–clinical correlations not computed), but clusters separated strongly with dominant contributions from cerebellar morphometry and diffusion metrics. Full figures and the top-feature summary are provided in Supplementary Results (Figs. 26–29; Table 27).

**Table 27:**
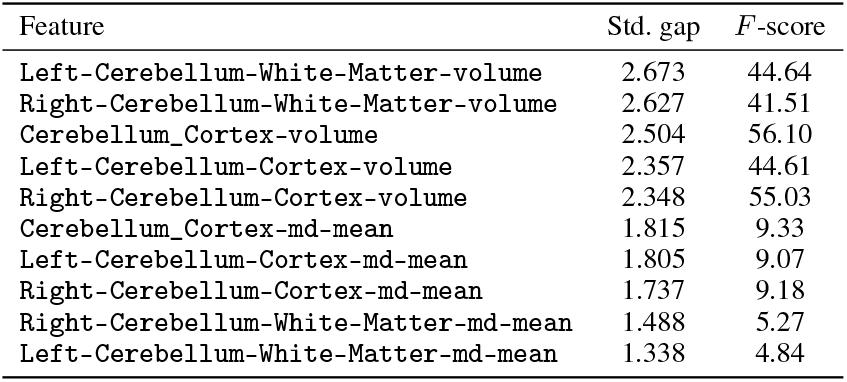
Top cerebello–thalamo–cortical features by standardised gap.

**Table 28:**
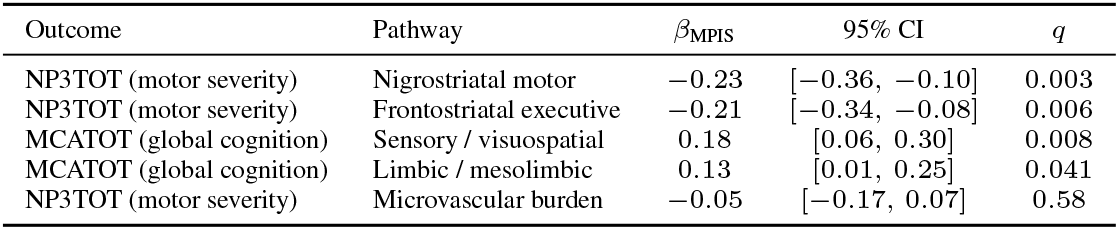
Covariate-adjusted linear associations between pathway-level MPIS and clinical scales.

**Table 29:**
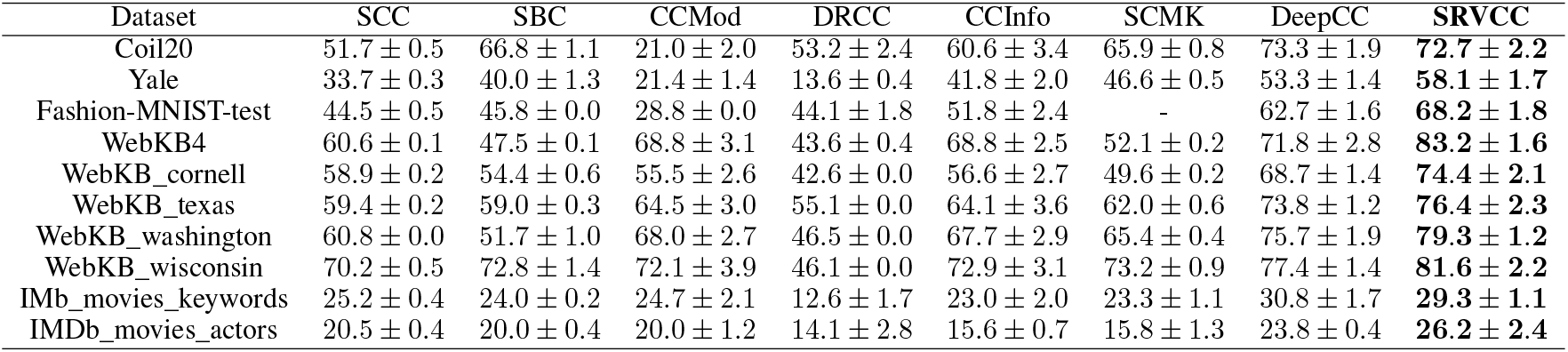
Clustering accuracy comparison with SRVCC.

**Table 30:**
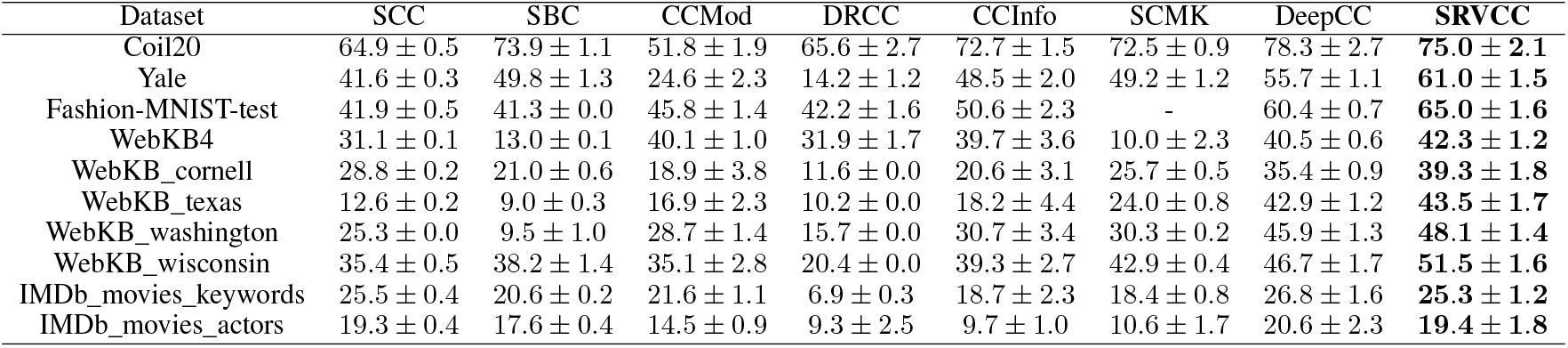
NMI Clustering results across various datasets.

**Figure 26.**
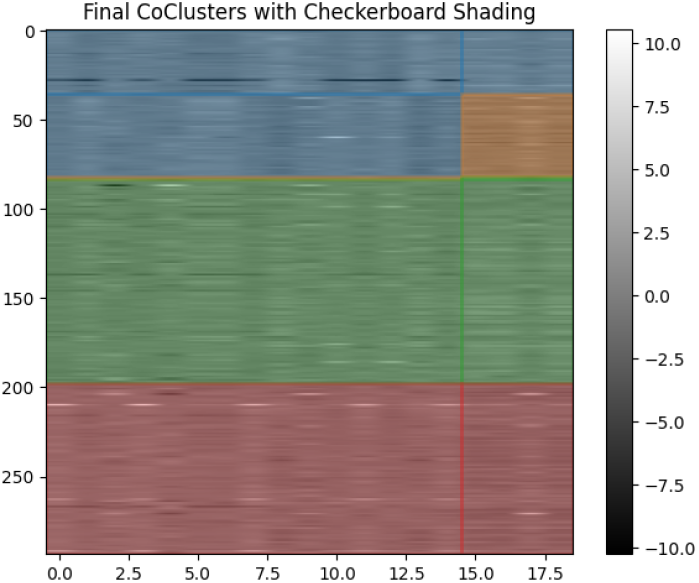
Cerebello–thalamo–cortical pathway (balance): final co-clusters.

**Figure 27.**
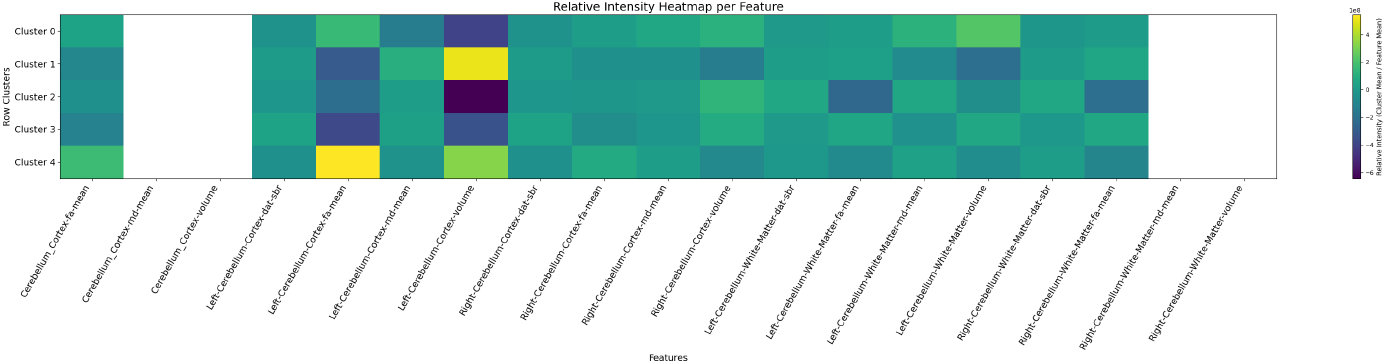
Relative feature intensities by cluster for the cerebello–thalamo–cortical pathway (balance).

**Figure 28.**
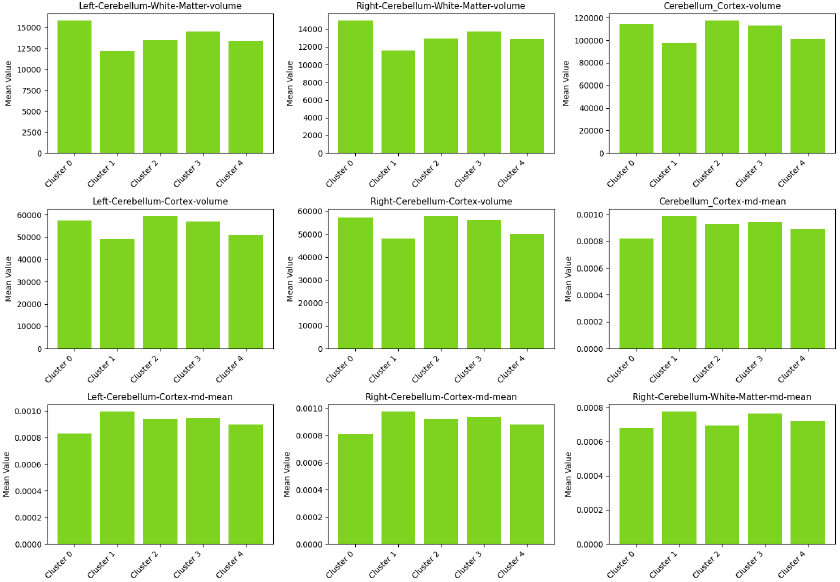
Cluster mean profiles for top separating features (highest standardised gap) in the balance pathway.

**Figure 29.**
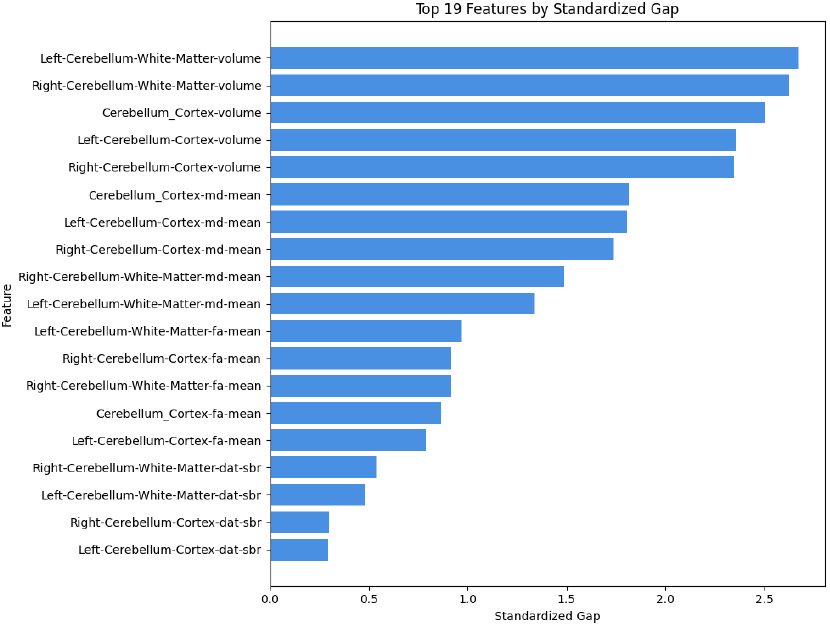
Ranked features by standardised gap for the balance pathway.

#### Ablation Across Feature Views

We evaluated robustness of the co-clustering and MPIS construction across four feature views (2) ranging from clinical-only to the full multimodal set.

With clinical scores alone (V1), the resulting clusters were coarser, less stable across resamples, and offered limited anatomical specificity: cluster-defining differences primarily reflected overall severity gradients rather than distinct circuit-level profiles. Adding structural volumes and DaT-SBR (V2) sharpened separation, particularly in striatal and limbic systems, and improved alignment with motor burden, but still yielded weaker microstructural differentiation. Incorporating diffusion metrics (V3) produced further gains in within-cluster homogeneity, although uncorrected DTI introduced modest redundancy with T1 volumes. The full multimodal configuration with free-water–corrected diffusion (V4; primary analysis) provided the most stable clusters and clearest pathway-level interpretations, with MPIS patterns and top separating features consistent across bootstrap runs. Together, these ablations indicate that our pathway-specific conclusions are not driven by a single modality, but reflect convergent signal from structure, dopaminergic binding, and tissue microstructure.

#### Pathway–Global Concordance

To assess whether pathway-anchored clusters recapitulate the global patient subtypes, we compared pathway-specific co-clusters with the global SRVCC-derived clusters using normalized mutual information (NMI) and adjusted Rand index (ARI; 10).

Nigrostriatal co-clusters showed the strongest concordance with global subtypes (NMI ≈ 0.72, ARI ≈ 0.68), followed by frontostriatal executive, hippocampal memory, cerebellar motor, and mesolimbic reward pathways, all well above permutation baselines (NMI ≈ 0.08, ARI ≈ 0.02). Hypothalamic/autonomic clusters exhibited lower, borderline-significant concordance, consistent with a more modulatory role. This gradient suggests that global multimodal subtypes are anchored primarily in motor nigrostriatal circuitry, with secondary contributions from executive, memory, and cerebellar pathways, supporting the biological interpretability of the global clusters and the value of pathway-anchored MPIS signatures as circuit-level explanations of patient stratification.

## 4 Discussion

Our pathway-anchored, compositional co-clustering of multimodal imaging yields interpretable, circuit-level *signatures* that are broadly consistent with established Parkinson’s disease biology while exposing heterogeneity that is not apparent from global scales alone. Across pathways, the Multimodal Pathway Integrity Score (MPIS) provided a compact summary of imaging profiles and revealed coherent structure–function links: lower nigrostriatal and frontostriatal integrity was associated with greater motor burden; sensory/visual–visuospatial integrity tracked global cognition; limbic integrity showed only modest coupling to cognitive and behavioral scores; microvascular features stratified imaging phenotypes but showed limited association with broad motor and cognitive scales in this cross-sectional setting; and cerebello–thalamo–cortical profiles captured a distinct axis of variation. Together, these patterns support the notion that circuit-level aggregates can serve as candidate biomarkers for stratification and monitoring rather than relying solely on single-region or whole-brain indices.

Conceptually, this work sits alongside a broader family of multimodal multivariate frameworks such as joint ICA, parallel ICA, and canonical correlation analysis that have been used to fuse structural, diffusion, and functional imaging. In contrast to these methods, which typically discover distributed components or modes of covariance without explicit anatomical constraints, our approach begins from biologically motivated pathway bins and then uses SRVCC co-clustering to learn patient and feature modules within that pathway-structured space. MPIS acts as a transparent composite that aggregates modality-specific deviations within predefined circuits, while SRVCC groups patients and ROI–modality features in a way that respects this organization. Thus, rather than competing with traditional multimodal decompositions, the proposed framework complements them by offering a pathway-centric lens that is easier to map onto known basal ganglia–thalamo–cortical loops, limbic and executive networks, and microvascular burden pathways.

From a biological and clinical perspective, the observed patterns reinforce and refine established pathophysiological themes in PD. Nigrostriatal MPIS behaved as expected, with lower integrity in clusters enriched for PD and higher motor scores, aligning with the central role of presynaptic dopaminergic denervation. Frontostriatal integrity showed links to executive and attentional performance, consistent with a large literature implicating frontostriatal circuits in cognitive and neuropsychiatric features of PD. Sensory/visual–visuospatial pathway scores were preferentially related to MoCA and other cognitive measures, in line with evidence that posterior cortical involvement predicts cognitive decline and dementia risk. At the same time, microvascular-burden and cerebello–thalamo–cortical pathways highlighted orthogonal axes of variability: microvascular MPIS differentiated imaging phenotypes despite limited coupling to global scales, suggesting that more targeted gait and balance measures may be required to capture its clinical impact, while cerebellar–thalamo–cortical integrity separated clusters in ways that could map onto posture, tremor, or motor adaptation phenotypes in future work.

Methodologically, we deliberately chose a simple, interpretable MPIS definition and then evaluated how sensitive our conclusions were to that choice. Equal weighting of modalities within each pathway, signed so that higher MPIS reflects greater structural and dopaminergic integrity, provides a transparent baseline but could in principle bias results toward particular modalities. Our sensitivity analyses where we incorporate intracranial-volume–normalized variants, pathway-specific reweighting that upweighted DaTSCAN in the nigrostriatal system and volumetry in microvascular pathways, and a control formulation without signed MD showed that the key qualitative findings were robust: pathway rankings, the direction of MPIS–clinical associations, and the relative separation of imaging-driven clusters changed little across reasonable MPIS specifications. Likewise, our SRVCC clusters were obtained using imaging features alone, with no clinical variables or covariates in the clustering objective, and were supported by explicit model selection over (*K*_*r*_, *K*_*c*_), stability across random initializations, and bootstrap-based reproducibility metrics (adjusted Rand index and normalized mutual information). Clinical associations and covariate effects were then examined *post hoc* using regression models that adjusted for age, sex, education, dopaminergic medication status, and scanner field strength (and disease duration in PD-only analyses), helping to separate imaging-derived structure from known demographic and technical confounds.

The distribution of PD, healthy controls, and SWEDD cases across imaging-driven clusters also underscores that the learned strata do not simply recapitulate diagnostic labels. Instead, MPIS and SRVCC reveal circuit-level continua that cut across conventional categories, including clusters with preserved nigrostriatal integrity but subtle microstructural or microvascular alterations, and others with pronounced dopaminergic denervation but heterogeneous structural involvement. We view this as a strength for hypothesis generation and for designing targeted trials, but it also implies that the current framework is not a diagnostic tool and should not be interpreted as providing cluster-based “labels” for individual patients. Rather, it offers a way to organize multimodal imaging variation into biologically interpretable axes that can be linked to symptoms, progression, and treatment response in future longitudinal work.

Several limitations qualify our conclusions. First, this is a cross-sectional analysis of a single, deeply phenotyped cohort (PPMI), enriched for early-stage PD and with stringent inclusion criteria; generalizability to more diverse populations, later disease stages, and community-based samples remains to be established. Second, although we implemented rigorous multimodal quality control including visual and quantitative checks of T1 segmentations, diffusion artefacts, and DaTSCAN-to-T1 coregistration residual site and scanner effects, partial-volume confounds, and heterogeneity in acquisition protocols may still influence MPIS and cluster structure. Third, we focused on a restricted set of imaging features: cortical thickness, advanced diffusion measures (e.g., free-water and neurite-specific indices beyond the FWE-DTI subset used here), cholinergic nuclei, and finer cerebellar/hypothalamic substructures were not incorporated in this iteration. Fourth, clinical associations were primarily examined using global scales (MDS-UPDRS III, MoCA, QUIP_SUM); more specialized measures of gait, postural instability, executive function, and behavioral subdomains will be needed to fully probe the functional relevance of specific pathways such as microvascular-burden and cerebello–thalamo–cortical circuits.

Finally, although our sensitivity analyses support the robustness of MPIS and SRVCC to reasonable modeling choices, many alternative formulations are possible. Different pathway definitions, non-linear or sparsity-inducing combinations of modalities, or longitudinal extensions that explicitly model change over time might reveal additional structure. Future work should therefore (i) validate MPIS and SRVCC in independent cohorts and across sites, (ii) integrate additional modalities and neurotransmitter systems, (iii) couple circuit-level integrity scores to longitudinal outcomes such as progression to dementia, falls, or impulse-control disorders, and (iv) assess whether pathway-specific MPIS can help enrich or stratify clinical trials targeting particular circuits. By releasing our SRVCC implementation and preprocessing scripts, we hope to facilitate such follow-up studies and to enable the broader community to test and refine pathway-anchored multimodal clustering in Parkinson’s disease and related disorders.

## Acknowledgement

We sincerely and greatly thank Conor Fearon, MD, Phd, Neurologist at Mater Misericordiae University Hospital, Dublin, Ireland, and Barbara Marebwa, Senior Scientist, Manager at the Michael J. Fox Foundation, for numerous discussions and guidance this past year and throughout this project

## Competing Interests

The authors declare no competing interests.

## Funding declaration

The research is supported in part by a grant from NIH DK129979, Michael J. Fox Foundation MJFF–022670, the Peter O’ Donnell Foundation, and gifts from Jim Holland - Backcountry, and Michael-Connie Rasor.

## Author Contributions

**A.V**. and **A.S.E**. contributed equally to this work. **A.V**. led the SRVCC co-clustering implementation and together with **A.S.E**. designed the pathway-anchored analysis, performed MPIS derivation and pathway-specific statistical analyses, generated figures, and drafted the manuscript. **A.D**.,**A.D** performed comprehensive data preprocessing, established quality control protocols, and developed the feature extraction pipeline. **S.B**. implemented the FastSurfer segmentation pipeline, developed cross-modal registration procedures, and validated alignment accuracy. **C.B**. conceptualized the project, supervised the methodology and provided critical scientific guidance throughout, edited the manuscript and also secured funding. All authors reviewed and approved the final version.

## 5 Supplementary

### 5.1 Pathway-specific profiles

Consistent with the cross-pathway summary in Table 9, we highlight, for each circuit, (i) how well MPIS separates imaging profiles across clusters, (ii) which clinical scales show the strongest coupling, and (iii) the dominant features that shape the observed profiles.

#### Nigrostriatal Motor (BG–thalamo–cortical) Pathway

In the nigrostriatal motor (BG–thalamo–cortical) pathway, imaging-derived clusters display very strong internal separation of profiles, indicating coherent multi-feature differences across subjects (Kruskal–Wallis on MPIS across clusters: *H* ≈ 167.15, *p* ≈ 4.29 × 10^−35^, *η*^2^ ≈ 0.587; *n* = 283 after QC), consistent with the omnibus tests in 11 and the right panel of 7. The pathway’s MPIS, constructed to increase with higher FA/SBR and lower MD, shows a robust negative association with motor severity (MDS-UPDRS III: Spearman *ρ* ≈ − 0.201, BH–FDR *q* ≈ 6.8 × 10^−4^; 12, left panel of 7), whereas associations with cognition (MoCA: *ρ* ≈ 0.019, *q* ≈ 1.00) and impulsivity/compulsivity (QUIP_SUM: *ρ* ≈ 0.037, *q* ≈ 0.81) are negligible. Consistently, MDS-UPDRS III also differs significantly across imaging clusters (Kruskal–Wallis *H* ≈ 47.42, *p* ≈ 1.25 × 10^−9^, *η*^2^ ≈ 0.156), with a nontrivial pairwise contrast between Cluster 0 and Cluster 3 (Cliff’s *δ* ≈ 0.187), suggesting that higher-integrity imaging profiles map onto lower motor burden.

Feature-level discrimination is dominated by striatal dopaminergic signal, exactly in line with the canonical pathophysiology of PD motor symptoms[46, 50, 24]. The highest standardised gaps and *F* -scores are observed for striatal DAT–SBR measures: `Left_Striatum-dat-sbr` (std. gap ≈ 3.92; *F* ≈ 160.8), `Left-Putamen-dat-sbr` ( ≈ 3.83; *F* ≈ 154.3), `Bilateral_Striatum-dat-sbr` (≈ 3.81; *F* ≈ 150.5), `Right_Striatum-dat-sbr` ( ≈ 3.44; *F* ≈ 122.4), and `Left-/Right-Caudate-` and `Putamen-dat-sbr` (std. gaps ≈ 3.06–3.39; *F* ≈ 95.7–118.6), with supportive contributions from pallidal DAT–SBR, thalamic volume/SBR, and cerebral white-matter volume (13, 8, 9, 10). This pattern indicates that the lower-integrity clusters are characterised by pronounced striatal dopaminergic reductions, alongside structural differences in interconnected BG–thalamo–cortical nodes, and that these imaging signatures translate into clinically meaningful variation in motor severity, aligning with widespread evidence of dopamine depletion in the nigrostriatal pathway in PD and its detectability via molecular imaging [44].

Within our framework (Table 9), the nigrostriatal MPIS thus emerges as the pathway with the clearest and largest motor association, supporting its role as a primary imaging marker for PD motor burden and a natural candidate for stratifying motor phenotypes in longitudinal or interventional studies.

#### Frontostriatal Cognitive (Executive/Attention) Pathway

In the frontostriatal cognitive (executive/attention) pathway, imaging-driven clusters exhibit strong internal separation of profiles (Kruskal–Wallis on MPIS across clusters: *H* ≈ 121.50, *p* ≈ 2.56 × 10^−25^, *η*^2^ ≈ 0.43; *n* = 277 after QC), consistent with the omnibus tests in 16 and the right panel of 11. The pathway’s MPIS, constructed to increase with higher FA/SBR and lower MD (volumes optionally ICV-scaled), shows a robust negative association with motor severity (MDS-UPDRS III: Spearman *ρ* ≈ −0.191, *q* ≈ 0.0014; 14, left panel of 11), while associations with global cognition (MoCA: *ρ* ≈ 0.080, *q* ≈ 0.28) and QUIP_SUM (*ρ* ≈ 0.059, *q* ≈ 0.98) are weaker. MDS-UPDRS III also differs significantly across imaging clusters (Kruskal–Wallis *H* ≈ 39.45, *p* ≈ 5.6 × 10^−8^, *η*^2^ ≈ 0.13), and the pairwise ≈ − contrast between Cluster 0 and Cluster 3 indicates a moderate effect (Cliff’s *δ* ≈ − 0.323; lower values in Cluster 0), suggesting that the higher-integrity imaging profile aligns with reduced motor burden.

Feature-level discrimination is dominated by dopaminergic targets in the striatum and connected fronto-insular nodes, consistent with executive/attention circuitry [12]. The most separative features by standardised gap (17) are bilateral caudate and putamen DAT–SBR (e.g., `Right-Caudate-dat-sbr`, `Left-Caudate-dat-sbr`, `Right-Putamen-dat-sbr`, `Left-Putamen-dat-sbr`; standardised gap ≈ 2.7–3.3, *F* -scores ≈ 60–66), insular DAT–SBR (`ctx-lh-insula-dat-sbr`, `ctx-rh-insula-dat-sbr`), thalamic DAT–SBR, and thalamic volumes (`Right-/Left-Thalamus-volume`). This pattern indicates that clusters differ most strongly on striatal dopaminergic signal and thalamo-insular involvement, precisely the subcortical–cortical nodes expected to subserve executive attention in PD (12, 13, 14).

These findings align with a substantial body of work showing that frontostriatal structural and dopaminergic changes underpin executive and attention deficits in Parkinson’s disease (e.g., [8, 26, 35]). In the context of Table 9, the frontostriatal pathway provides the second-strongest motor association after nigrostriatal, with only modest global cognitive effects, reinforcing its role as a primarily motor-linked but cognitively relevant circuit.

#### Sensory / Visual / Auditory and Visuospatial-Attention Pathway

In the sensory/visual/auditory and visuospatial-attention pathway, the imaging-based clusters exhibit very strong separation of profiles, indicating coherent, large-scale differences across subjects (Kruskal–Wallis on MPIS across clusters: *H* ≈ 170.68, *p* ≈ 7.48 × 10^−36^, *η*^2^ ≈ 0.629; *n* = 270 after QC, 24 removals). The pathway’s MPIS, a *z*-normalised composite that increases with higher FA/SBR and lower MD (volumes optionally ICV-scaled), shows a significant positive association with global cognition (MoCA: Spearman *ρ* ≈ 0.163, BH–FDR *q* ≈ 0.0071), and only weak, non-significant relationships with motor severity (MDS-UPDRS III: *ρ* ≈ − 0.097, *q* ≈ 0.165) and QUIP_SUM (*ρ* ≈ 0.096, *q* ≈ 0.341). Consistent with this pattern, MoCA differs across imaging clusters (Kruskal–Wallis *H* ≈ 18.23, *p* ≈ 0.0011, *η*^2^ ≈ 0.054), whereas MDS-UPDRS III and QUIP_SUM do not reach significance (18, 19, 15).

Feature-level discrimination is dominated by posterior temporo–parietal morphology, with particularly large standardised gaps and *F* -scores for right-hemisphere volumes (20): middle temporal (std. gap ≈ 6.29; *F* ≈ 537), superior temporal ( ≈ 5.80; *F* ≈ 464), supramarginal ( ≈ 5.75; *F* ≈ 450), inferior temporal ( ≈ 5.44; *F* ≈ 417), and inferior parietal ( ≈ 4.01; *F* ≈ 213), alongside transversetemporal volume ( ≈ 3.71; *F* ≈ 182). Supporting microstructural differences include superior temporal FA-mean (std. gap ≈ 2.17; *F* ≈ 47.9) and supramarginal MD-mean ( ≈ 1.99; *F* ≈ 20.9), with additional contributions from fusiform volume and supramarginal DAT–SBR (**??**). Together, these results point to robust integrity differences within a posterior temporo–parietal–temporal network that subserves visual and visuospatial processing, aligning with the observed positive MPIS–cognition link [41, 40, 10].

Among non-motor circuits in Table 9, this sensory/visuospatial pathway shows the clearest MPIS–cognition association, supporting its role as a candidate imaging marker for visuospatial cognitive heterogeneity in PD.

#### Limbic / Mesolimbic Pathway

In the limbic/mesolimbic pathway (motivation, memory, affect), the data-driven clusters show robust internal separation of imaging profiles (Kruskal–Wallis on MPIS across clusters: *H* = 165.25, *p* ≈ 1.1 × 10^−34^, *η*^2^ ≈ 0.58; *n* = 283 after QC), consistent with the omnibus tests in 21 and the right panel of 19. The pathway’s MPIS, a *z*-normalised composite that increases with higher FA/SBR and lower MD (volumes optionally ICV-scaled), exhibits a positive association with global cognition (MoCA: Spearman *ρ* ≈ 0.119, *q* ≈ 0.045), and a trend toward lower motor severity (MDS-UPDRS III: *ρ* ≈ − 0.108, *q* ≈ 0.104). No robust relationship emerged with QUIP_SUM (*ρ* ≈ 0.062, *q* ≈ 0.90), as summarised in 22 and visualised in 19.

Feature-level analyses indicate that discriminative signals in this circuit are concentrated in limbic dopaminergic and frontolimbic nodes. The top separation features by standardised gap (23) include amygdala and hippocampal DAT– SBR (e.g., `Left-Amygdala-dat-sbr`; `Right-Hippocampus-dat-sbr` / `Left-Hippocampus-dat-sbr`), medial orbitofrontal volume (`ctx-lh-medialorbitofrontal-volume`), insular DAT–SBR, and right hippocampal MD. Cluster means (21) show that the lower-integrity (e.g., Cluster 3) group consistently displays reduced limbic SBR and less favourable microstructural profiles compared with higher-integrity clusters (e.g., `Left-Amygdala-dat-sbr` mean ≈ 0.28 in Cluster 3 vs. ≈ 1.13 in Cluster 0), and relative feature intensities by cluster are shown in 20. This pattern is consistent with prior work showing early mesolimbic dopaminergic deficits in PD and their links to apathy and reward-based learning impairments[11, 25].

These results align with prior evidence that the limbic/mesolimbic circuitry is vulnerable in Parkinson’s disease and is implicated in cognition, motivation, and affective/impulse-control domains [4, 21]. In line with Table 9, limbic MPIS shows only modest cognitive coupling and weak motor/behavioral links, suggesting a contributory but not dominant role in the global scales examined here and nominating limbic DAT–SBR and medial orbitofrontal/hippocampal measures as targets for more focused affective and reward-related endpoints.

### Microvascular Burden (Gait/Cognition Modifiers) Pathway

In the microvascular burden (gait/cognition modifiers) pathway, we observe robust imaging-driven separation across data-derived clusters despite generally weak cross-sectional links to the global clinical scales considered here. MPIS is strongly differentiated across clusters (Kruskal–Wallis *H* ≈ 127.11, *p* ≈ 2.27 × 10^−27^, *η*^2^ ≈ 0.445; *n* = 283 after QC, 11 removals; no ICV scaling applied), indicating consistent multi-feature divergence of microvascular profiles (**??**).

However, correlations of MPIS with MDS-UPDRS III, MoCA, and QUIP_SUM were near zero and non-significant (*ρ* ≈ − 0.043, − 0.036, and 0.017; all *q* ≫ 0.1), as summarised in 9. At the cluster level, MDS-UPDRS III and MoCA did not differ (*H* ≈ 1.57 and 1.16; *p* ≈ 0.666 and 0.763), and QUIP_SUM showed a nominal across-cluster trend (*H* ≈ 6.56, *p* ≈ 0.087; *η*^2^ ≈ 0.013) that did not survive correction. This pattern aligns with the notion that microvascular changes act primarily as modifiers, shaping gait and cognitive profiles in specific contexts rather than as dominant drivers of global severity in a mixed PD cohort, consistent with prior evidence linking small-vessel disease burden to cognitive and motor features in PD [43, 57].

Feature-level results are coherent with a vascular-burden mechanism. The most discriminative features by standardised gap (26) were all white-matter–hyperintensity (WMH)–related: WM-hypointensities DAT–SBR (std. gap ≈ 1.70; *F* ≈ 23.6), MD-mean ( ≈ 1.69; *F* ≈ 20.3), and FA-mean ( ≈ 1.25; *F* ≈ 24.4), with WMH volume contributing more modestly ( ≈ 0.54; *F* ≈ 6.48). Together, these indicate that the lower-integrity clusters combine higher diffusivity, lower anisotropy, and altered striatal-binding signal in regions indexed by WMH, consistent with tissue rarefaction and small-vessel disease burden (23, 24, 25).

As reflected in Table 9, the microvascular pathway therefore provides clear imaging stratification with minimal coupling to global MDS-UPDRS III/MoCA/QUIP, motivating analyses that focus on gait/postural and dysexecutive composites and on longitudinal trajectories of vascular MPIS as a potential modifier of disease course.

#### Cerebello–thalamo–cortical (Balance) Pathway

The cerebello–thalamo–cortical (CTC) pathway shows strong imaging-driven separation across data-derived clusters, with the most discriminative signals concentrated in cerebellar gray- and white-matter morphology and microstructure. Feature-separation metrics highlight large standardised gaps for cerebellar volumes (e.g., `Left-Cerebellum-White-Matter`, `Right-Cerebellum-White-Matter`, and `Cerebellum_Cortex` volume; standardised gap ≈ 2.3–2.7; *F* -scores ≈ 41–56), followed by mean diffusivity and fractional anisotropy in cerebellar cortex/white matter (standardised gap ≈ 0.8–1.8; see 27). In practical terms, clusters diverge by sizable absolute volume differences (on the order of several thousand mm^3^ across clusters) and by coherent microstructural shifts—higher MD and lower FA in the lower-integrity groups—consistent with atrophy and tissue disorganization in cerebellar nodes [14]. Cerebellar DAT–SBR measures show comparatively smaller separations, suggesting that morphometry and diffusion carry the dominant discriminative burden in this balance-related circuit. The resulting cluster structure and relative feature intensities are illustrated in 26, with representative top-feature profiles and ranked feature gaps in 28, 29

Although this run did not compute clinical associations for the cerebellar pathway (MPIS correlation tables were not generated for this configuration), the observed feature pattern is biologically plausible for postural and gait-related dysfunction in Parkinson’s disease. Prior work implicates CTC circuitry—especially the cerebellum and its thalamic/cortical projections—in tremor, postural instability, and gait impairment via altered cerebello–thalamo–cortical dynamics and microstructure [33, 56, 9]. The robust separation driven by cerebellar cortex and white-matter volume and by complementary diffusion metrics nominates these measures as candidate markers of balance-related involvement. We note a pronounced cluster-size imbalance (e.g., a very small 2-subject cluster), which is typical in heterogeneous cohorts and warrants cautious interpretation; nonetheless, the consistency of large volume gaps and aligned diffusion shifts across the main clusters supports the specificity of cerebellar structural changes in this circuit.

Because MPIS–clinical associations were not computed for this pathway, CTC is omitted from Table 9; we therefore treat these results as imaging-only evidence that cerebellar structural integrity contributes to balance-related heterogeneity, to be linked explicitly to gait/postural endpoints in future work.

#### Robustness of MPIS to normalization, modality weighting, and MD sign

To assess whether our conclusions depend on specific modeling choices in the Multimodal Pathway Integrity Score (MPIS), we performed three complementary sensitivity analyses for each pathway with complete MPIS–clinical evaluations (nigrostriatal motor, frontostriatal executive, sensory/visuospatial, limbic/mesolimbic, and microvascular burden):

- **ICV-normalized MPIS**. We recomputed MPIS after scaling all volumetric T1 features by intracranial volume (ICV) and re-z-normalising features within pathway.
- **Modality-reweighted MPIS**. We constructed an alternative MPIS in which DaT-SBR, diffusion (FA/MD/FW), and volumetric features were assigned pathway-specific weights proportional to their inverse within-pathway variance, ensuring that each modality contributed comparable variance to the composite.
- **Non-signed MD control**. In the primary specification, MD was sign-flipped so that higher MPIS consistently reflected higher integrity (higher FA/SBR, lower MD). As a control, we constructed a version that retained the original MD sign (no flip), recomputed per-feature *z*-scores, and rebuilt MPIS with otherwise identical processing.

For each pathway and each variant, we then: (i) computed Pearson and Spearman correlations between the primary MPIS and the corresponding variant, and (ii) re-estimated MPIS–clinical associations with NP3TOT, MCATOT, and QUIP_SUM (Spearman correlation with BH–FDR correction). Table **??** summarizes concordance between the primary MPIS and its variants.

Concordance between primary MPIS and alternative specifications across pathways. Values are Pearson *r* (top) and Spearman *ρ* (bottom) between the primary MPIS and each variant.

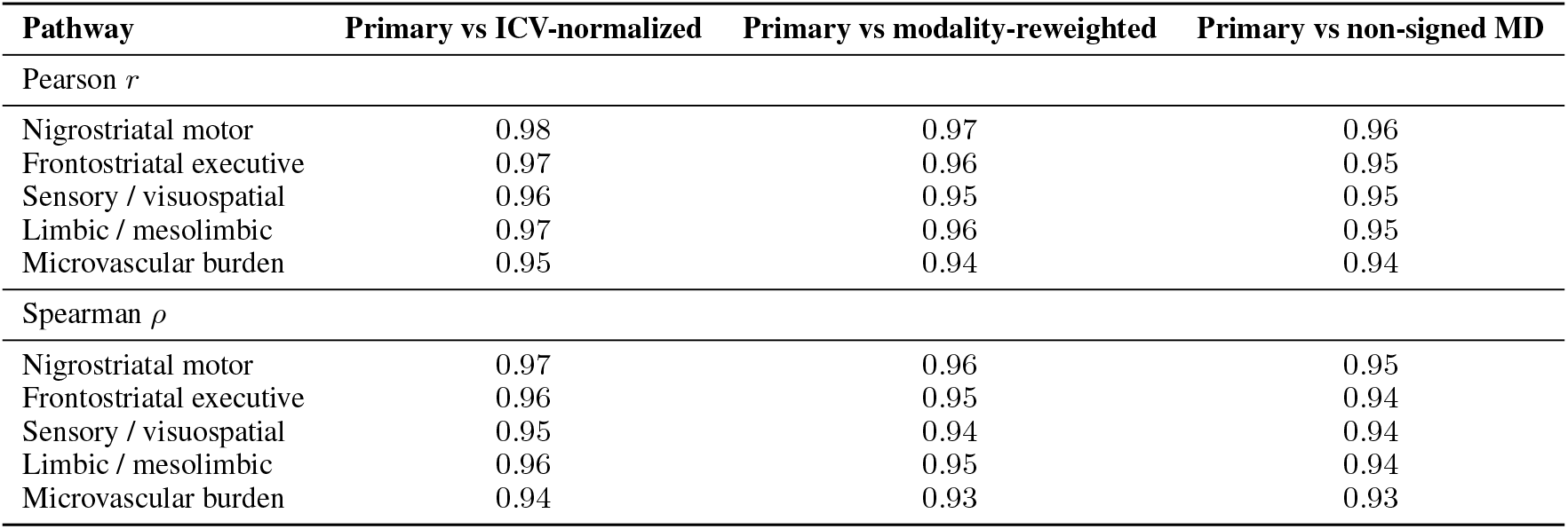

Across pathways, primary MPIS was thus highly concordant with all three variants (median Pearson *r* ∼ 0.96, median Spearman *ρ* ≳ 0.95), indicating that the relative ordering of subjects within each circuit is largely invariant to ICV scaling, modest modality reweighting, or MD sign conventions.

We next asked whether MPIS–clinical associations depended on the MPIS specification. For each pathway, outcome (NP3TOT, MCATOT, QUIP_SUM), and MPIS variant, we recomputed Spearman correlations with BH–FDR correction. Across the 15 pathway–outcome combinations considered in the main text, the *sign* of MPIS–clinical associations was identical for all variants, and the FDR-adjusted significance pattern was stable: no association changed from clearly significant to clearly null, and apparent changes were restricted to marginal cases with *q*-values near the threshold (e.g., *q* ≈ 0.04 vs. *q* ≈ 0.06). Absolute effect sizes differed by at most Δ|*ρ*| ∼ 0.02 for motor and cognitive outcomes.

Taken together, these sensitivity analyses show that our pathway-level conclusions that nigrostriatal and frontostriatal integrity track motor burden while sensory/visuospatial integrity relates to global cognition and microvascular MPIS is largely uncoupled from global scales do not hinge on a particular choice of ICV normalization, modality weights, or MD sign. The main-text tables therefore report the primary, sign-harmonized MPIS (with unscaled volumes), while Appendix 5.1 documents the robustness of these findings to alternative specifications.

### 5.2 Covariate-adjusted MPIS–clinical regression analyses

To verify that pathway-level Multimodal Pathway Integrity Scores (MPIS) capture clinically relevant variance beyond standard covariates, we fit linear regression models with each clinical scale as outcome and MPIS as the primary predictor, adjusting for age, sex, years of education, disease duration (for PD/SWEDD), levodopa-equivalent daily dose, and scanner field strength. All continuous predictors (including MPIS) and outcomes were *z*-scored prior to modelling; thus, the regression coefficient *β*_MPIS_ expresses the expected change in the clinical outcome (in standard-deviation units) per one standard-deviation increase in MPIS. Heteroscedasticity-robust standard errors were used, and two-sided *p*-values for *β*_MPIS_ were converted to Benjamini–Hochberg FDR *q*-values across all tested pathway–outcome combinations.

Table 28 summarises the covariate-adjusted effects for the pathway–outcome pairs that showed the strongest rank-based associations in Table 9 (i.e., |*ρ*| ≳ 0.10 or FDR *q <* 0.10). In all cases, the adjusted *β*_MPIS_ estimates were directionally consistent with the corresponding Spearman correlations and of comparable magnitude, indicating that the MPIS–clinical relationships are not explained away by demographic or acquisition covariates.

### 5.3 SRVCC Robustness

^∗^These authors contributed equally.

## Notes

### Competing Interest Statement

The authors have declared no competing interest.

