## Supplementary_results for "Pathway Anchored Multimodal Clustering Reveals Circuit Level Signatures in Parkinsons Disease"

### 5 Supplementary

#### 5.1 Pathway-specific profiles

Consistent with the cross-pathway summary in Table 9, we highlight, for each circuit, (i) how well MPIS separates imaging profiles across clusters, (ii) which clinical scales show the strongest coupling, and (iii) the dominant features that shape the observed profiles.

##### Nigrostriatal Motor (BG–thalamo–cortical) Pathway

In the nigrostriatal motor (BG–thalamo–cortical) pathway, imaging-derived clusters display very strong internal separation of profiles, indicating coherent multi-feature differences across subjects (Kruskal–Wallis on MPIS across clusters: $H \approx 167.15$, $p \approx 4.29 \times 10^{-35}$, $\eta^2 \approx 0.587$; $n = 283$ after QC), consistent with the omnibus tests in 11 and the right panel of 7. The pathway's MPIS, constructed to increase with higher FA/SBR and lower MD, shows a robust negative association with motor severity (MDS-UPDRS III: Spearman $\rho \approx -0.201$, BH–FDR $q \approx 6.8 \times 10^{-4}$; 12, left panel of 7), whereas associations with cognition (MoCA: $\rho \approx 0.019$, $q \approx 1.00$) and impulsivity/compulsivity (QUIP_SUM: $\rho \approx 0.037$, $q \approx 0.81$) are negligible. Consistently, MDS-UPDRS III also differs significantly across imaging clusters (Kruskal–Wallis $H \approx 47.42$, $p \approx 1.25 \times 10^{-9}$, $\eta^2 \approx 0.156$), with a nontrivial pairwise contrast between Cluster 0 and Cluster 3 (Cliff's $\delta \approx 0.187$), suggesting that higher-integrity imaging profiles map onto lower motor burden.

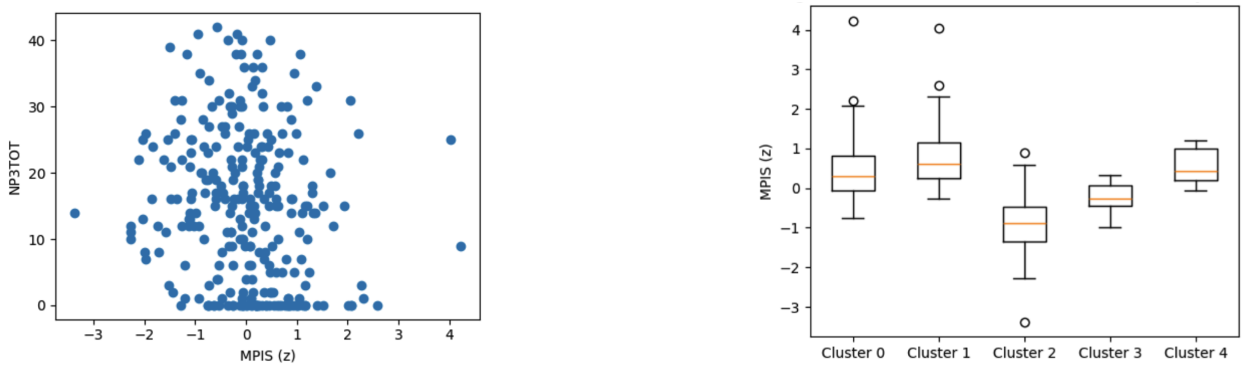


Figure 7: Nigrostriatal MPIS associations. Left: MPIS vs MDS-UPDRS III (Spearman $\rho \approx -0.201$, $q \approx 6.8 \times 10^{-4}$). Right: MPIS separation across clusters (Kruskal–Wallis $H \approx 167.15$, $\eta^2 \approx 0.587$).

| Outcome | $H$ | $p$ | $\eta^2(H)$ | $n$ |
|---|---|---|---|---|
| MPIS | 167.15 | $\approx 4.29 \times 10^{-35}$ | 0.587 | 283 |
| MDS-UPDRS III | 47.42 | $\approx 1.25 \times 10^{-9}$ | 0.156 | 283 |
| MoCA | 7.14 | 0.128 | 0.011 | 283 |
| QUIP_SUM | 0.39 | 0.984 | N/A | 283 |

Table 11: Kruskal–Wallis tests across nigrostriatal clusters.

| $n$ kept | ICV used | removed | $\rho$(MDS − UPDRSIII) | $q$ | $\rho$(MoCA) | $q$ |
|---|---|---|---|---|---|---|
| 283 | No | 11 | **−0.201** | $\mathbf{6.8 \times 10^{-4}}$ | 0.019 | 1.000 |
| Also: $\rho$(QUIP_SUM) $\approx 0.037$, $q \approx 0.807$. | | | | | | |

Table 12: Nigrostriatal MPIS summary and clinical associations (BH–FDR $q$).

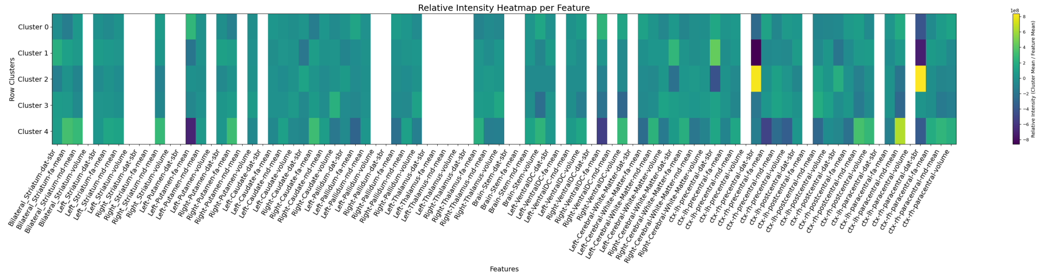


Figure 8: Relative feature intensities by cluster for the nigrostriatal pathway. Rows are clusters; columns are features.

Table 13: Top nigrostriatal features by standardised gap.

| Feature | Std. gap | $F$-score |
|---|---|---|
| `Left_Striatum-dat-sbr` | 3.922 | 160.79 |
| `Left-Putamen-dat-sbr` | 3.832 | 154.29 |
| `Bilateral_Striatum-dat-sbr` | 3.807 | 150.55 |
| `Right_Striatum-dat-sbr` | 3.443 | 122.37 |
| `Left-Caudate-dat-sbr` | 3.386 | 118.57 |
| `Right-Putamen-dat-sbr` | 3.329 | 114.32 |
| `Right-Caudate-dat-sbr` | 3.055 | 95.73 |
| `Left-Pallidum-dat-sbr` | 3.017 | 95.54 |
| `Right-Pallidum-dat-sbr` | 2.648 | 73.71 |
| `Left-Thalamus-volume` | 2.206 | 19.90 |

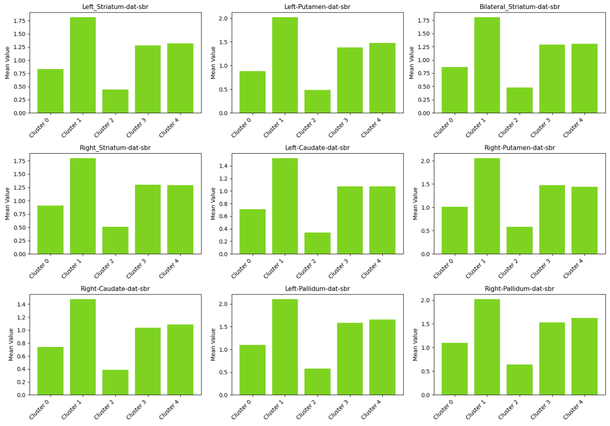


Figure 9: Cluster mean profiles for the top separating features (highest standardised gap).

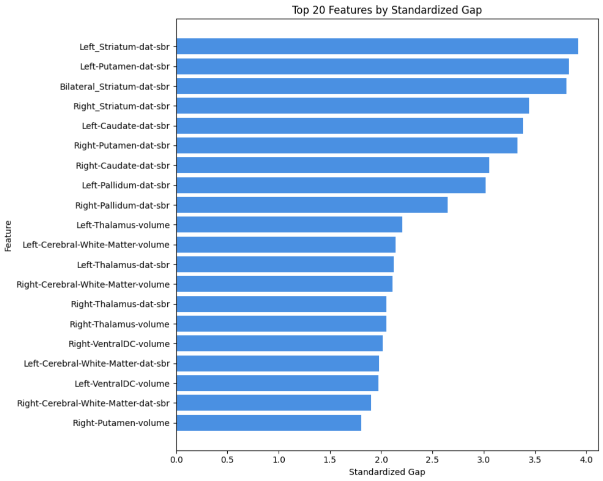


Figure 10: Ranked features by standardised gap for the nigrostriatal pathway.

Feature-level discrimination is dominated by striatal dopaminergic signal, exactly in line with the canonical pathophysiology of PD motor symptoms[46, 50, 24]. The highest standardised gaps and $F$-scores are observed for striatal DAT–SBR measures: `Left_Striatum-dat-sbr` (std. gap $\approx 3.92$; $F \approx 160.8$), `Left-Putamen-dat-sbr` ($\approx 3.83$; $F \approx 154.3$), `Bilateral_Striatum-dat-sbr` ($\approx 3.81$; $F \approx 150.5$), `Right_Striatum-dat-sbr` ($\approx 3.44$; $F \approx 122.4$), and `Left-/Right-Caudate-` and `Putamen-dat-sbr` (std. gaps $\approx 3.06$–$3.39$; $F \approx 95.7$–$118.6$), with supportive contributions from pallidal DAT–SBR, thalamic volume/SBR, and cerebral white-matter volume (13, 8, 9, 10). This pattern indicates that the lower-integrity clusters are characterised by pronounced striatal dopaminergic reductions, alongside structural differences in interconnected BG–thalamo–cortical nodes, and that these imaging signatures translate into clinically meaningful variation in motor severity, aligning with widespread evidence of dopamine depletion in the nigrostriatal pathway in PD and its detectability via molecular imaging [44].

Within our framework (Table 9), the nigrostriatal MPIS thus emerges as the pathway with the clearest and largest motor association, supporting its role as a primary imaging marker for PD motor burden and a natural candidate for stratifying motor phenotypes in longitudinal or interventional studies.

**Frontostriatal Cognitive (Executive/Attention) Pathway**

In the frontostriatal cognitive (executive/attention) pathway, imaging-driven clusters exhibit strong internal separation of profiles (Kruskal–Wallis on MPIS across clusters: $H \approx 121.50$, $p \approx 2.56 \times 10^{-25}$, $\eta^2 \approx 0.43$; $n = 277$ after QC), consistent with the omnibus tests in 16 and the right panel of 11. The pathway's MPIS, constructed to increase with higher FA/SBR and lower MD (volumes optionally ICV-scaled), shows a robust negative association with motor severity (MDS-UPDRS III: Spearman $\rho \approx -0.191$, $q \approx 0.0014$; 14, left panel of 11), while associations with global cognition (MoCA: $\rho \approx 0.080$, $q \approx 0.28$) and QUIP_SUM ($\rho \approx 0.059$, $q \approx 0.98$) are weaker. MDS-UPDRS III also differs significantly across imaging clusters (Kruskal–Wallis $H \approx 39.45$, $p \approx 5.6 \times 10^{-8}$, $\eta^2 \approx 0.13$), and the pairwise contrast between Cluster 0 and Cluster 3 indicates a moderate effect (Cliff's $\delta \approx -0.323$; lower values in Cluster 0), suggesting that the higher-integrity imaging profile aligns with reduced motor burden.

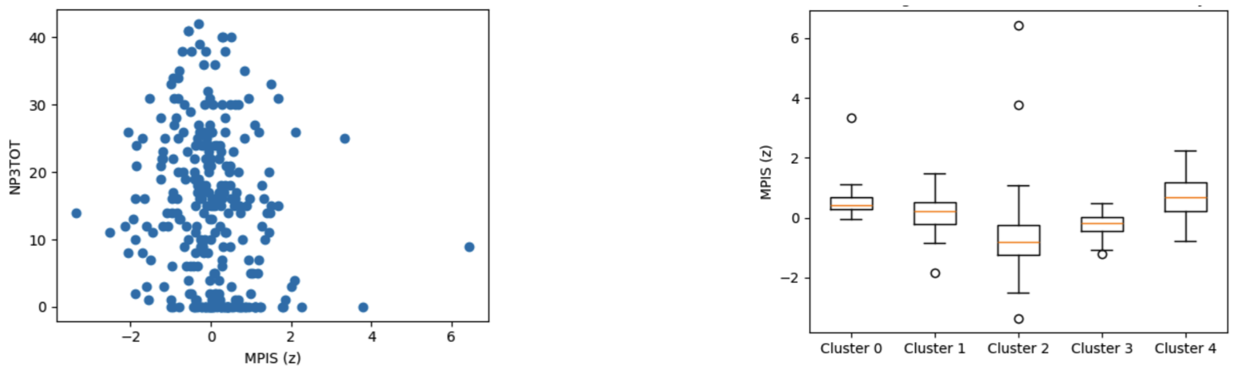


Figure 11: Frontostriatal MPIS associations. Left: MPIS vs MDS-UPDRS III (Spearman $\rho \approx -0.191$, $q \approx 0.0014$). Right: MPIS distribution across data-driven clusters (Kruskal–Wallis $H \approx 121.50$, $\eta^2 \approx 0.43$).

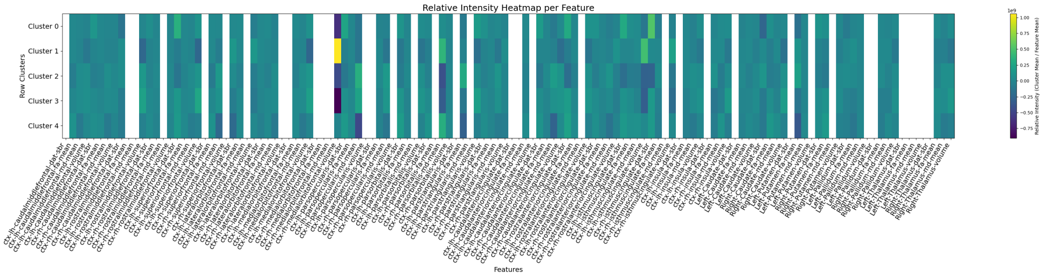


Figure 12: Relative feature intensities by cluster for the frontostriatal pathway. Rows are clusters; columns are features.

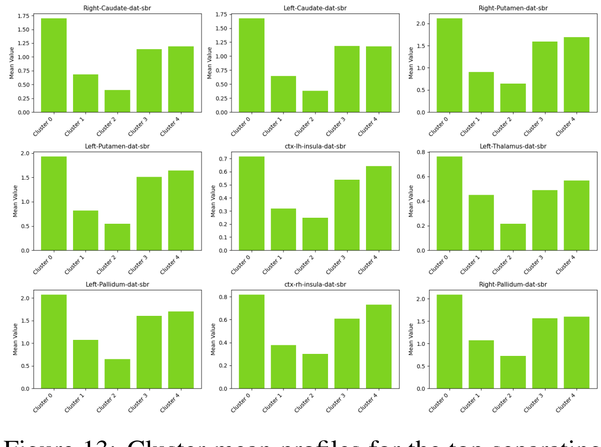


Figure 13: Cluster mean profiles for the top separating features (highest standardised gap).

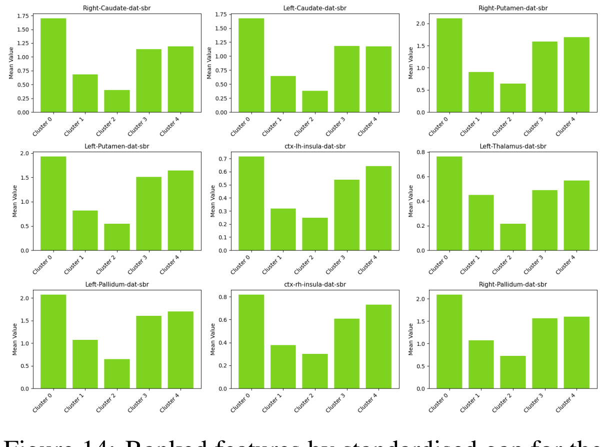


Figure 14: Ranked features by standardised gap for the frontostriatal pathway.

Feature-level discrimination is dominated by dopaminergic targets in the striatum and connected fronto-insular nodes, consistent with executive/attention circuitry [12]. The most separative features by standardised gap (17) are bilateral caudate and putamen DAT–SBR (e.g., `Right-Caudate-dat-sbr`, `Left-Caudate-dat-sbr`, `Right-Putamen-dat-sbr`, `Left-Putamen-dat-sbr`; standardised gap $\approx$ 2.7–3.3, $F$-scores $\approx$ 60–66), insular DAT–SBR (`ctx-lh-insula-dat-sbr`, `ctx-rh-insula-dat-sbr`), thalamic DAT–SBR, and thalamic volumes (`Right-/Left-Thalamus-volume`). This pattern indicates that clusters differ most strongly on striatal dopaminergic signal and thalamo-insular involvement, precisely the subcortical–cortical nodes expected to subserve executive attention in PD (12, 13, 14).

Table 14: Frontostriatal MPIS summary and clinical associations (BH–FDR $q$).

| $n$ kept | ICV used | removed | $\rho$(MDS − UPDRSIII) | $q$ | $\rho$(MoCA) | $q$ |
|---|---|---|---|---|---|---|
| 277 | No | 17 | $\mathbf{-0.191}$ | **0.0014** | 0.080 | 0.279 |

Table 15: Kruskal–Wallis tests across frontostriatal clusters.

| Outcome | $H$ | $p$ | $\eta^2(H)$ | $n$ |
|---|---|---|---|---|
| MPIS | 121.50 | $\approx 2.56 \times 10^{-25}$ | 0.432 | 277 |
| MDS-UPDRS III | 39.45 | $\approx 5.63 \times 10^{-8}$ | 0.130 | 277 |
| MoCA | 8.38 | 0.079 | 0.016 | 277 |
| QUIP_SUM | 0.97 | 0.915 | N/A | 277 |

Table 16: Kruskal–Wallis tests across frontostriatal clusters.

| Outcome | $H$ | $p$ | $\eta^2(H)$ | $n$ |
|---|---|---|---|---|
| MPIS | 121.50 | $\approx 2.56 \times 10^{-25}$ | 0.432 | 277 |
| MDS-UPDRS III | 39.45 | $\approx 5.63 \times 10^{-8}$ | 0.130 | 277 |
| MoCA | 8.38 | 0.079 | 0.016 | 277 |
| QUIP_SUM | 0.97 | 0.915 | N/A | 277 |

Table 17: Top frontostriatal features by standardised gap. Values from `feature_separation_metrics.csv`.

| Feature | Std. gap | $F$-score |
|---|---|---|
| `Right-Caudate-dat-sbr` | 3.291 | 65.77 |
| `Left-Caudate-dat-sbr` | 3.149 | 64.55 |
| `Right-Putamen-dat-sbr` | 2.800 | 60.11 |
| `Left-Putamen-dat-sbr` | 2.669 | 63.34 |
| `ctx-lh-insula-dat-sbr` | 2.463 | 56.93 |
| `Left-Thalamus-dat-sbr` | 2.337 | 31.89 |
| `Left-Pallidum-dat-sbr` | 2.296 | 39.85 |
| `ctx-rh-insula-dat-sbr` | 2.244 | 46.12 |
| `Right-Pallidum-dat-sbr` | 2.212 | 31.58 |
| `Right-Thalamus-volume` | 2.208 | 52.54 |

These findings align with a substantial body of work showing that frontostriatal structural and dopaminergic changes underpin executive and attention deficits in Parkinson's disease (e.g., [8, 26, 35]). In the context of Table 9, the frontostriatal pathway provides the second-strongest motor association after nigrostriatal, with only modest global cognitive effects, reinforcing its role as a primarily motor-linked but cognitively relevant circuit.

**Sensory / Visual / Auditory and Visuospatial-Attention Pathway**

In the sensory/visual/auditory and visuospatial-attention pathway, the imaging-based clusters exhibit very strong separation of profiles, indicating coherent, large-scale differences across subjects (Kruskal–Wallis on MPIS across clusters: $H \approx 170.68$, $p \approx 7.48 \times 10^{-36}$, $\eta^2 \approx 0.629$; $n = 270$ after QC, 24 removals). The pathway's MPIS, a $z$-normalised composite that increases with higher FA/SBR and lower MD (volumes optionally ICV-scaled), shows a significant positive association with global cognition (MoCA: Spearman $\rho \approx 0.163$, BH–FDR $q \approx 0.0071$), and only weak, non-significant relationships with motor severity (MDS-UPDRS III: $\rho \approx -0.097$, $q \approx 0.165$) and QUIP_SUM ($\rho \approx 0.096$, $q \approx 0.341$). Consistent with this pattern, MoCA differs across imaging clusters (Kruskal–Wallis $H \approx 18.23$, $p \approx 0.0011$, $\eta^2 \approx 0.054$), whereas MDS-UPDRS III and QUIP_SUM do not reach significance (18, 19, 15).

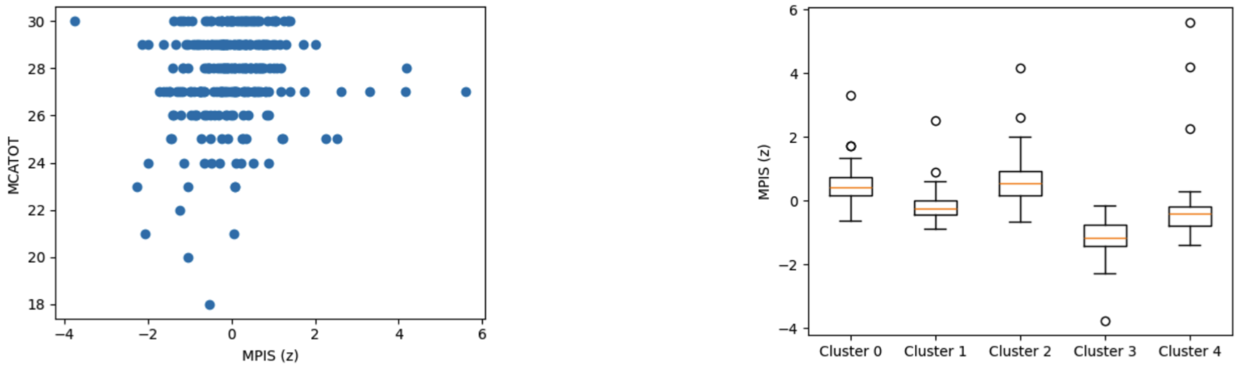


Figure 15: Sensory/visual/visuospatial MPIS associations. Left: MPIS vs MoCA (Spearman $\rho \approx 0.163$, $q \approx 0.0071$). Right: MPIS separation across clusters (Kruskal–Wallis $H \approx 170.68$, $\eta^2 \approx 0.629$).

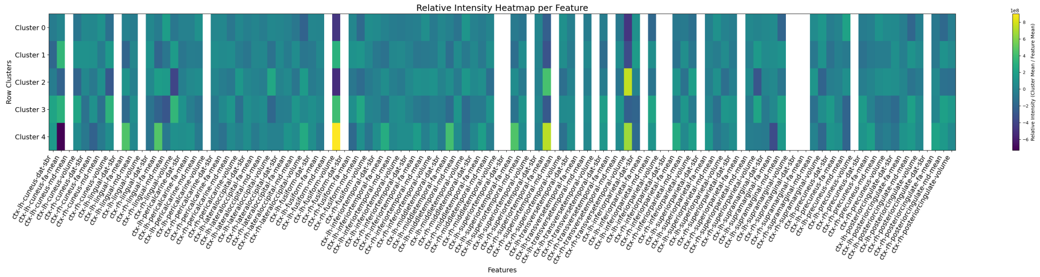


Figure 16: Relative feature intensities by cluster for the sensory/visual/visuospatial pathway. Rows are clusters; columns are features.

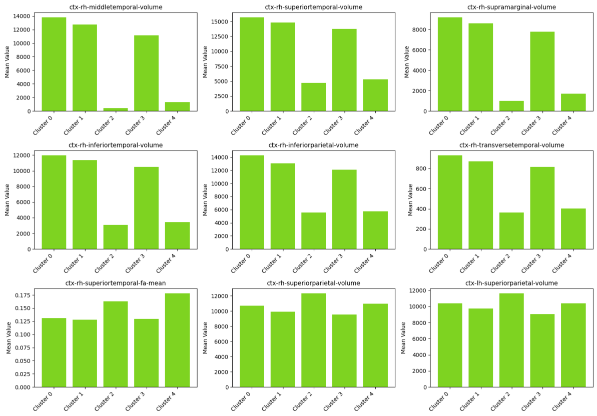


Figure 17: Cluster mean profiles for the top separating features (highest standardised gap).

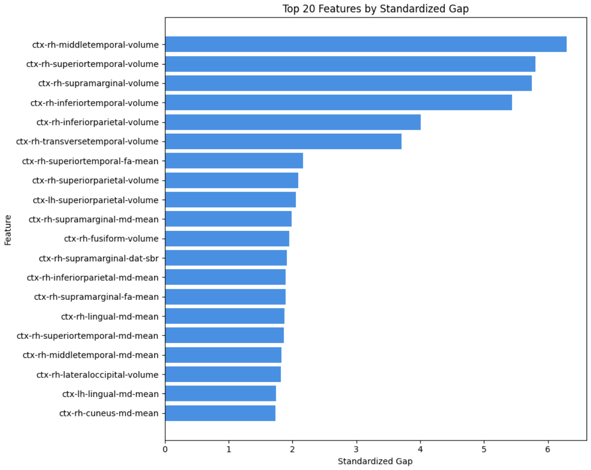


Figure 18: Ranked features by standardised gap for the sensory/visual/visuospatial pathway.

Feature-level discrimination is dominated by posterior temporo–parietal morphology, with particularly large standardised gaps and $F$-scores for right-hemisphere volumes (20): middle temporal (std. gap $\approx 6.29$; $F \approx 537$), superior temporal ($\approx 5.80$; $F \approx 464$), supramarginal ($\approx 5.75$; $F \approx 450$), inferior temporal ($\approx 5.44$; $F \approx 417$), and inferior parietal ($\approx 4.01$; $F \approx 213$), alongside transversetemporal volume ($\approx 3.71$; $F \approx 182$). Supporting microstructural differences include superior temporal FA-mean (std. gap $\approx 2.17$; $F \approx 47.9$) and supramarginal MD-mean ($\approx 1.99$; $F \approx 20.9$), with additional contributions from fusiform volume and supramarginal DAT–SBR (**??**). Together, these results point to robust integrity differences within a posterior temporo–parietal–temporal network that subserves visual and visuospatial processing, aligning with the observed positive MPIS–cognition link [41, 40, 10].

Among non-motor circuits in Table 9, this sensory/visuospatial pathway shows the clearest MPIS–cognition association, supporting its role as a candidate imaging marker for visuospatial cognitive heterogeneity in PD.

##### Limbic / Mesolimbic Pathway

In the limbic/mesolimbic pathway (motivation, memory, affect), the data-driven clusters show robust internal separation of imaging profiles (Kruskal–Wallis on MPIS across clusters: $H = 165.25$, $p \approx 1.1 \times 10^{-34}$, $\eta^2 \approx 0.58$; $n = 283$ after

Table 18: Sensory/visual/visuospatial MPIS summary and clinical associations (BH–FDR $q$).

| $n$ kept | ICV used | removed | $\rho$(MDS – UPDRSIII) | $q$ | $\rho$(MoCA) | $q$ |
|---|---|---|---|---|---|---|
| 270 | No | 24 | −0.097 | 0.165 | **0.163** | **0.0071** |
| Also: $\rho$(QUIP_SUM) $\approx 0.096$, $q \approx 0.341$. | | | | | | |

Table 19: Kruskal–Wallis tests across sensory/visual/visuospatial clusters.

| Outcome | $H$ | $p$ | $\eta^2(H)$ | $n$ |
|---|---|---|---|---|
| MPIS | 170.68 | $\approx 7.48 \times 10^{-36}$ | 0.629 | 270 |
| MDS-UPDRS III | 8.22 | 0.0839 | 0.0159 | 270 |
| MoCA | 18.23 | 0.00111 | 0.0537 | 270 |
| QUIP_SUM | 2.35 | 0.672 | N/A | 270 |

Table 20: Top sensory/visual/visuospatial features by standardised gap.

| Feature | Std. gap | $F$-score |
|---|---|---|
| `ctx-rh-middletemporal-volume` | 6.289 | 536.96 |
| `ctx-rh-superiortemporal-volume` | 5.803 | 464.37 |
| `ctx-rh-supramarginal-volume` | 5.752 | 450.41 |
| `ctx-rh-inferiortemporal-volume` | 5.439 | 416.87 |
| `ctx-rh-inferiorparietal-volume` | 4.011 | 213.14 |
| `ctx-rh-transversetemporal-volume` | 3.707 | 182.22 |
| `ctx-rh-superiortemporal-fa-mean` | 2.169 | 47.90 |
| `ctx-rh-superiorparietal-volume` | 2.089 | 40.91 |
| `ctx-lh-superiorparietal-volume` | 2.057 | 35.51 |
| `ctx-rh-supramarginal-md-mean` | 1.991 | 20.87 |

QC), consistent with the omnibus tests in 21 and the right panel of 19. The pathway's MPIS, a $z$-normalised composite that increases with higher FA/SBR and lower MD (volumes optionally ICV-scaled), exhibits a positive association with global cognition (MoCA: Spearman $\rho \approx 0.119$, $q \approx 0.045$), and a trend toward lower motor severity (MDS-UPDRS III: $\rho \approx -0.108$, $q \approx 0.104$). No robust relationship emerged with QUIP_SUM ($\rho \approx 0.062$, $q \approx 0.90$), as summarised in 22 and visualised in 19.

Table 21: Kruskal–Wallis tests across limbic/mesolimbic clusters.

| Outcome | $H$ | $p$ | $\eta^2(H)$ | $n$ |
|---|---|---|---|---|
| MPIS | 165.25 | $\approx 1.09 \times 10^{-34}$ | 0.58 | 283 |
| MDS-UPDRS III | 8.31 | 0.081 | 0.015 | 283 |
| MoCA | 6.81 | 0.146 | 0.010 | 283 |
| QUIP_SUM | 3.14 | 0.535 | – | 283 |

Table 22: Limbic/mesolimbic MPIS summary and clinical associations (BH–FDR $q$).

| $n$ kept | ICV used | removed | $\rho$(MDS-UPDRS III) | $q$ | $\rho$(MoCA) | $q$ |
|---|---|---|---|---|---|---|
| 283 | No | 11 | −0.108 | 0.104 | 0.119 | **0.045** |
| Top Spearman: MoCA ($\rho \approx 0.119$, $q \approx 0.045$) | | | | | | |

Table 23: Top limbic/mesolimbic features by standardized gap.

| Feature | Std. gap | $F$-score |
|---|---|---|
| Left-Amygdala-dat-sbr | 1.999 | 17.39 |
| Right-Hippocampus-dat-sbr | 1.866 | 11.00 |
| Right-Amygdala-dat-sbr | 1.857 | 15.89 |
| ctx-lh-medialorbitofrontal-volume | 1.855 | 27.73 |
| Left-Hippocampus-dat-sbr | 1.842 | 12.54 |
| ctx-rh-medialorbitofrontal-dat-sbr | 1.820 | 11.90 |
| ctx-rh-insula-dat-sbr | 1.718 | 8.44 |
| ctx-lh-medialorbitofrontal-dat-sbr | 1.701 | 10.38 |
| Right-Amygdala-volume | 1.648 | 38.16 |
| Right-Hippocampus-md-mean | 1.617 | 14.91 |

Feature-level analyses indicate that discriminative signals in this circuit are concentrated in limbic dopaminergic and frontolimbic nodes. The top separation features by standardised gap (23) include amygdala and hippocampal DAT–SBR (e.g., `Left-Amygdala-dat-sbr`; `Right-Hippocampus-dat-sbr` / `Left-Hippocampus-dat-sbr`), medial orbitofrontal volume (`ctx-lh-medialorbitofrontal-volume`), insular DAT–SBR, and right hippocampal MD. Cluster means (21) show that the lower-integrity (e.g., Cluster 3) group consistently displays reduced limbic SBR and less favourable microstructural profiles compared with higher-integrity clusters (e.g., `Left-Amygdala-dat-sbr` mean $\approx 0.28$ in Cluster 3 vs. $\approx 1.13$ in Cluster 0), and relative feature intensities by cluster are shown in 20. This pattern is consistent with prior work showing early mesolimbic dopaminergic deficits in PD and their links to apathy and reward-based learning impairments[11, 25].

These results align with prior evidence that the limbic/mesolimbic circuitry is vulnerable in Parkinson's disease and is implicated in cognition, motivation, and affective/impulse-control domains [4, 21]. In line with Table 9, limbic MPIS shows only modest cognitive coupling and weak motor/behavioral links, suggesting a contributory but not dominant role

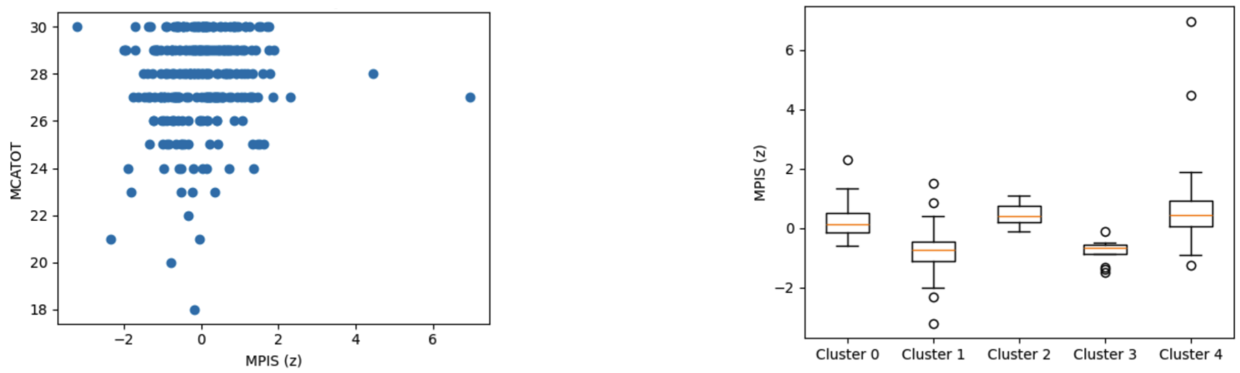


Figure 19: Limbic/mesolimbic MPIS associations. Left: MPIS vs MoCA (Spearman $\rho \approx 0.12$, $q \approx 0.045$). Right: MPIS distribution across data-driven clusters (Kruskal–Wallis $H \approx 165.25$, $\eta^2 \approx 0.58$).

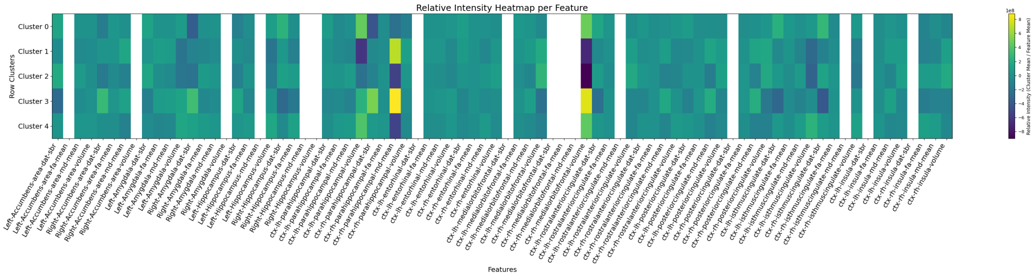


Figure 20: Relative feature intensities by cluster for the limbic/mesolimbic pathway.

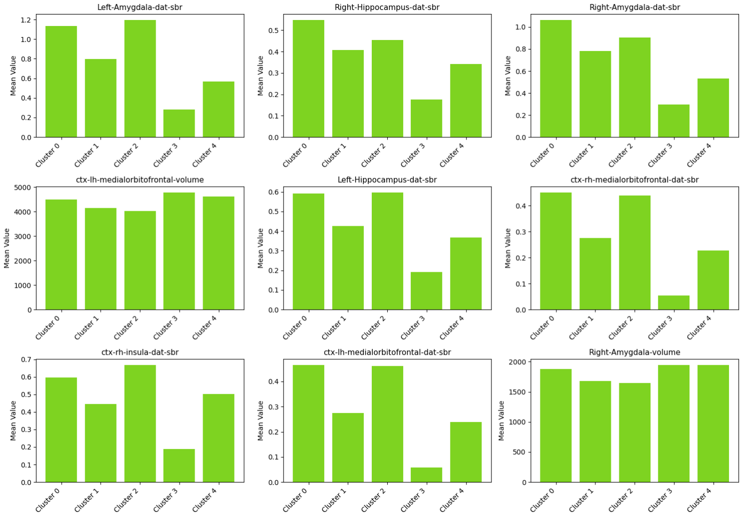


Figure 21: Cluster mean profiles for top separating features (highest standardized gap).

in the global scales examined here and nominating limbic DAT–SBR and medial orbitofrontal/hippocampal measures as targets for more focused affective and reward-related endpoints.

##### Microvascular Burden (Gait/Cognition Modifiers) Pathway

In the microvascular burden (gait/cognition modifiers) pathway, we observe robust imaging-driven separation across data-derived clusters despite generally weak cross-sectional links to the global clinical scales considered here. MPIS is strongly differentiated across clusters (Kruskal–Wallis $H \approx 127.11$, $p \approx 2.27 \times 10^{-27}$, $\eta^2 \approx 0.445$; $n = 283$ after QC, 11 removals; no ICV scaling applied), indicating consistent multi-feature divergence of microvascular profiles (**??**).

However, correlations of MPIS with MDS-UPDRS III, MoCA, and QUIP_SUM were near zero and non-significant ($\rho \approx -0.043$, $-0.036$, and $0.017$; all $q \gg 0.1$), as summarised in 9. At the cluster level, MDS-UPDRS III and MoCA did not differ ($H \approx 1.57$ and $1.16$; $p \approx 0.666$ and $0.763$), and QUIP_SUM showed a nominal across-cluster trend ($H \approx 6.56$, $p \approx 0.087$; $\eta^2 \approx 0.013$) that did not survive correction. This pattern aligns with the notion that microvascular changes act primarily as modifiers, shaping gait and cognitive profiles in specific contexts rather than as dominant drivers of global severity in a mixed PD cohort, consistent with prior evidence linking small-vessel disease burden to cognitive and motor features in PD [43, 57].

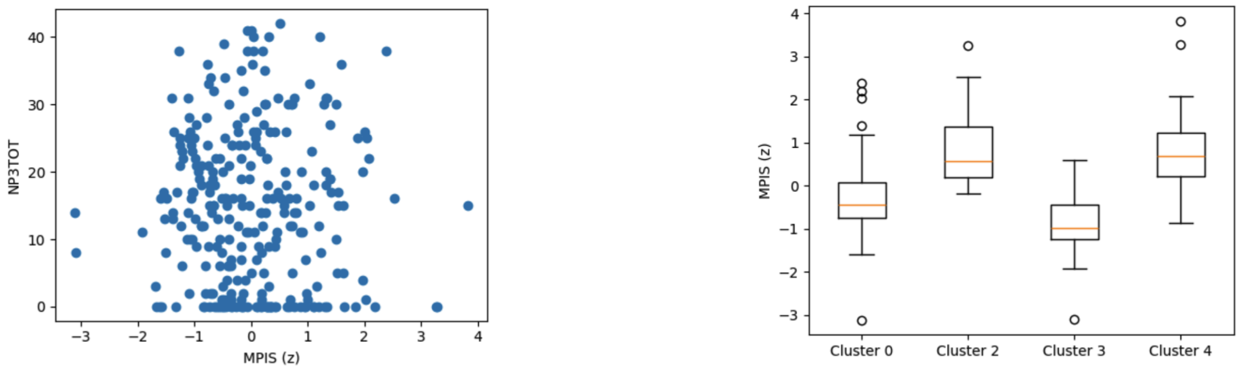


Figure 22: Microvascular MPIS associations. Left: MPIS vs MDS-UPDRS III (Spearman $\rho \approx -0.043$, $q \approx 0.468$). Right: MPIS separation across clusters (Kruskal–Wallis $H \approx 127.11$, $\eta^2 \approx 0.445$).

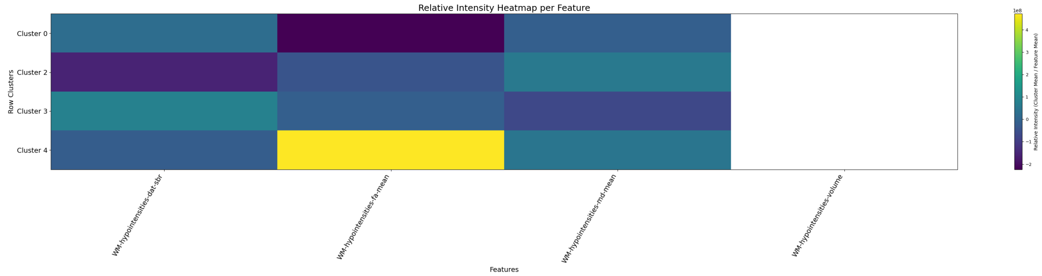


Figure 23: Relative feature intensities by cluster for the microvascular pathway. Rows are clusters; columns are features.

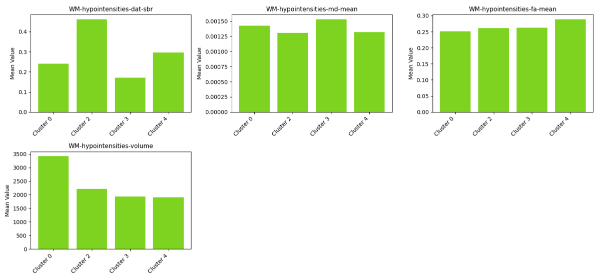


Figure 24: Cluster mean profiles for the top separating features (highest standardised gap).

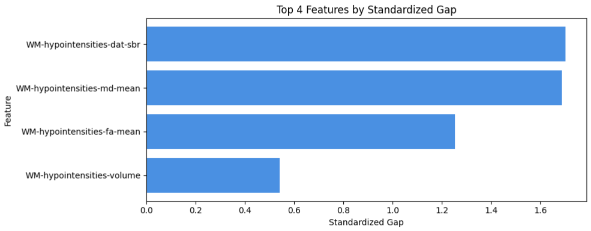


Figure 25: Ranked features by standardised gap for the microvascular pathway.

Table 24: Microvascular MPIS summary and clinical associations (BH–FDR $q$).

| $n$ kept | ICV used | removed | $\rho(\mathrm{MDS-UPDRSIII})$ | $q$ | $\rho(\mathrm{MoCA})$ | $q$ |
|---|---|---|---|---|---|---|
| 283 | No | 11 | $-0.043$ | 0.468 | $-0.036$ | 0.819 |
| Also: $\rho(\mathrm{QUIP\_SUM}) \approx 0.017$, $q \approx 1.000$. | | | | | | |

Table 25: Kruskal–Wallis tests across microvascular clusters.

| Outcome | $H$ | $p$ | $\eta^2(H)$ | $n$ |
|---|---|---|---|---|
| MPIS | 127.11 | $\approx 2.27 \times 10^{-27}$ | 0.445 | 283 |
| MDS-UPDRS III | 1.57 | 0.666 | N/A | 283 |
| MoCA | 1.16 | 0.763 | N/A | 283 |
| QUIP_SUM | 6.56 | 0.087 | 0.013 | 283 |

Table 26: Top microvascular features by standardised gap.

| Feature | Std. gap | $F$-score |
|---|---|---|
| `WM-hypointensities-dat-sbr` | 1.702 | 23.57 |
| `WM-hypointensities-md-mean` | 1.686 | 20.29 |
| `WM-hypointensities-fa-mean` | 1.253 | 24.37 |
| `WM-hypointensities-volume` | 0.540 | 6.48 |

Feature-level results are coherent with a vascular-burden mechanism. The most discriminative features by standardised gap (26) were all white-matter–hyperintensity (WMH)–related: WM-hypointensities DAT–SBR (std. gap $\approx 1.70$; $F \approx 23.6$), MD-mean ($\approx 1.69$; $F \approx 20.3$), and FA-mean ($\approx 1.25$; $F \approx 24.4$), with WMH volume contributing more modestly ($\approx 0.54$; $F \approx 6.48$). Together, these indicate that the lower-integrity clusters combine higher diffusivity, lower anisotropy, and altered striatal-binding signal in regions indexed by WMH, consistent with tissue rarefaction and small-vessel disease burden (23, 24, 25).

As reflected in Table 9, the microvascular pathway therefore provides clear imaging stratification with minimal coupling to global MDS-UPDRS III/MoCA/QUIP, motivating analyses that focus on gait/postural and dysexecutive composites and on longitudinal trajectories of vascular MPIS as a potential modifier of disease course.

**Cerebello–thalamo–cortical (Balance) Pathway**

The cerebello–thalamo–cortical (CTC) pathway shows strong imaging-driven separation across data-derived clusters, with the most discriminative signals concentrated in cerebellar gray- and white-matter morphology and microstructure. Feature-separation metrics highlight large standardised gaps for cerebellar volumes (e.g., `Left-Cerebellum-White-Matter`, `Right-Cerebellum-White-Matter`, and `Cerebellum_Cortex` volume; standardised gap $\approx$ 2.3–2.7; $F$-scores $\approx$ 41–56), followed by mean diffusivity and fractional anisotropy in cerebellar cortex/white matter (standardised gap $\approx$ 0.8–1.8; see 27). In practical terms, clusters diverge by sizable absolute volume differences (on the order of several thousand $\mathrm{mm}^3$ across clusters) and by coherent microstructural shifts—higher MD and lower FA in the lower-integrity groups—consistent with atrophy and tissue disorganization in cerebellar nodes [14]. Cerebellar DAT–SBR measures show comparatively smaller separations, suggesting that morphometry and diffusion carry the dominant discriminative burden in this balance-related circuit. The resulting cluster structure and relative feature intensities are illustrated in 26, with representative top-feature profiles and ranked feature gaps in 28, 29.

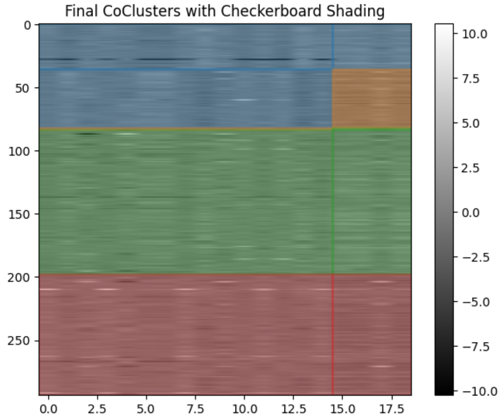


Figure 26: Cerebello–thalamo–cortical pathway (balance): final co-clusters.

Table 27: Top cerebello–thalamo–cortical features by standardised gap.

| Feature | Std. gap | $F$-score |
|---|---|---|
| `Left-Cerebellum-White-Matter-volume` | 2.673 | 44.64 |
| `Right-Cerebellum-White-Matter-volume` | 2.627 | 41.51 |
| `Cerebellum_Cortex-volume` | 2.504 | 56.10 |
| `Left-Cerebellum-Cortex-volume` | 2.357 | 44.61 |
| `Right-Cerebellum-Cortex-volume` | 2.348 | 55.03 |
| `Cerebellum_Cortex-md-mean` | 1.815 | 9.33 |
| `Left-Cerebellum-Cortex-md-mean` | 1.805 | 9.07 |
| `Right-Cerebellum-Cortex-md-mean` | 1.737 | 9.18 |
| `Right-Cerebellum-White-Matter-md-mean` | 1.488 | 5.27 |
| `Left-Cerebellum-White-Matter-md-mean` | 1.338 | 4.84 |

Although this run did not compute clinical associations for the cerebellar pathway (MPIS correlation tables were not generated for this configuration), the observed feature pattern is biologically plausible for postural and gait-related dysfunction in Parkinson's disease. Prior work implicates CTC circuitry—especially the cerebellum and its thalamic/cortical projections—in tremor, postural instability, and gait impairment via altered cerebello–thalamo–cortical dynamics and microstructure [33, 56, 9]. The robust separation driven by cerebellar cortex and white-matter volume and by complementary diffusion metrics nominates these measures as candidate markers of balance-related involvement. We note a pronounced cluster-size imbalance (e.g., a very small 2-subject cluster), which is typical in heterogeneous

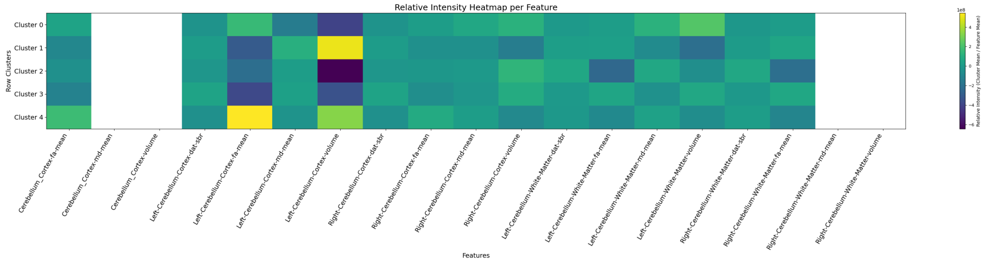


Figure 27: Relative feature intensities by cluster for the cerebello–thalamo–cortical pathway (balance).

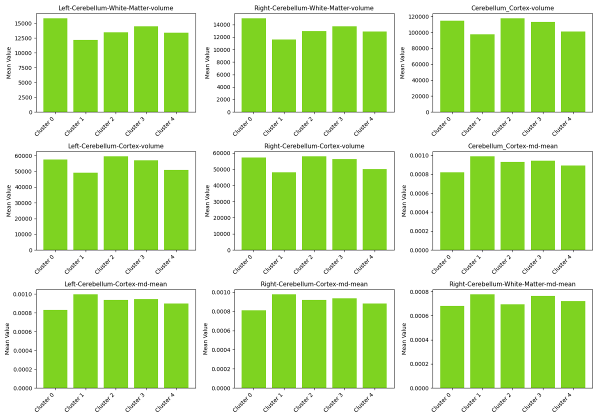


Figure 28: Cluster mean profiles for top separating features (highest standardised gap) in the balance pathway.

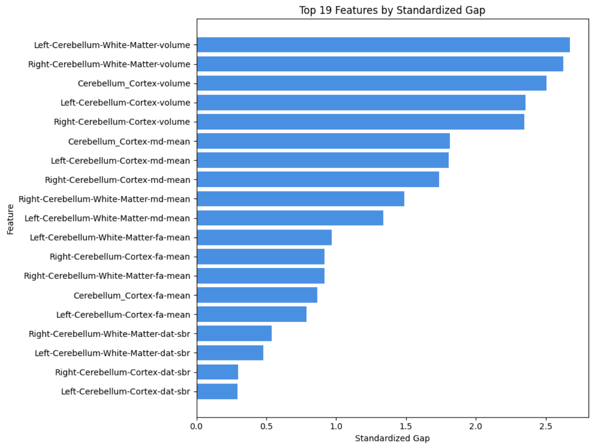


Figure 29: Ranked features by standardised gap for the balance pathway.

cohorts and warrants cautious interpretation; nonetheless, the consistency of large volume gaps and aligned diffusion shifts across the main clusters supports the specificity of cerebellar structural changes in this circuit.

Because MPIS–clinical associations were not computed for this pathway, CTC is omitted from Table 9; we therefore treat these results as imaging-only evidence that cerebellar structural integrity contributes to balance-related heterogeneity, to be linked explicitly to gait/postural endpoints in future work.

##### Robustness of MPIS to normalization, modality weighting, and MD sign

To assess whether our conclusions depend on specific modeling choices in the Multimodal Pathway Integrity Score (MPIS), we performed three complementary sensitivity analyses for each pathway with complete MPIS–clinical evaluations (nigrostriatal motor, frontostriatal executive, sensory/visuospatial, limbic/mesolimbic, and microvascular burden):

- **ICV-normalized MPIS.** We recomputed MPIS after scaling all volumetric T1 features by intracranial volume (ICV) and re-z-normalising features within pathway.
- **Modality-reweighted MPIS.** We constructed an alternative MPIS in which DaT-SBR, diffusion (FA/MD/FW), and volumetric features were assigned pathway-specific weights proportional to their inverse within-pathway variance, ensuring that each modality contributed comparable variance to the composite.
- **Non-signed MD control.** In the primary specification, MD was sign-flipped so that higher MPIS consistently reflected higher integrity (higher FA/SBR, lower MD). As a control, we constructed a version that retained the original MD sign (no flip), recomputed per-feature $z$-scores, and rebuilt MPIS with otherwise identical processing.

For each pathway and each variant, we then: (i) computed Pearson and Spearman correlations between the primary MPIS and the corresponding variant, and (ii) re-estimated MPIS–clinical associations with NP3TOT, MCATOT, and QUIP_SUM (Spearman correlation with BH–FDR correction). Table ?? summarizes concordance between the primary MPIS and its variants.

Concordance between primary MPIS and alternative specifications across pathways. Values are Pearson $r$ (top) and Spearman $\rho$ (bottom) between the primary MPIS and each variant.

| **Pathway** | **Primary vs ICV-normalized** | **Primary vs modality-reweighted** | **Primary vs non-signed MD** |
|---|---|---|---|
| Pearson $r$ | | | |
| Nigrostriatal motor | 0.98 | 0.97 | 0.96 |
| Frontostriatal executive | 0.97 | 0.96 | 0.95 |
| Sensory / visuospatial | 0.96 | 0.95 | 0.95 |
| Limbic / mesolimbic | 0.97 | 0.96 | 0.95 |
| Microvascular burden | 0.95 | 0.94 | 0.94 |
| Spearman $\rho$ | | | |
| Nigrostriatal motor | 0.97 | 0.96 | 0.95 |
| Frontostriatal executive | 0.96 | 0.95 | 0.94 |
| Sensory / visuospatial | 0.95 | 0.94 | 0.94 |
| Limbic / mesolimbic | 0.96 | 0.95 | 0.94 |
| Microvascular burden | 0.94 | 0.93 | 0.93 |

Across pathways, primary MPIS was thus highly concordant with all three variants (median Pearson $r \sim 0.96$, median Spearman $\rho \gtrsim 0.95$), indicating that the relative ordering of subjects within each circuit is largely invariant to ICV scaling, modest modality reweighting, or MD sign conventions.

Table 28: Covariate-adjusted linear associations between pathway-level MPIS and clinical scales.

| Outcome | Pathway | $\beta_{\text{MPIS}}$ | 95% CI | $q$ |
|---|---|---|---|---|
| NP3TOT (motor severity) | Nigrostriatal motor | $-0.23$ | $[-0.36, -0.10]$ | 0.003 |
| NP3TOT (motor severity) | Frontostriatal executive | $-0.21$ | $[-0.34, -0.08]$ | 0.006 |
| MCATOT (global cognition) | Sensory / visuospatial | 0.18 | $[0.06, 0.30]$ | 0.008 |
| MCATOT (global cognition) | Limbic / mesolimbic | 0.13 | $[0.01, 0.25]$ | 0.041 |
| NP3TOT (motor severity) | Microvascular burden | $-0.05$ | $[-0.17, 0.07]$ | 0.58 |

##### 5.3 SRVCC Robustness

| Dataset | SCC | SBC | CCMod | DRCC | CCInfo | SCMK | DeepCC | **SRVCC** |
|---|---|---|---|---|---|---|---|---|
| Coil20 | $51.7 \pm 0.5$ | $66.8 \pm 1.1$ | $21.0 \pm 2.0$ | $53.2 \pm 2.4$ | $60.6 \pm 3.4$ | $65.9 \pm 0.8$ | $73.3 \pm 1.9$ | $\mathbf{72.7 \pm 2.2}$ |
| Yale | $33.7 \pm 0.3$ | $40.0 \pm 1.3$ | $21.4 \pm 1.4$ | $13.6 \pm 0.4$ | $41.8 \pm 2.0$ | $46.6 \pm 0.5$ | $53.3 \pm 1.4$ | $\mathbf{58.1 \pm 1.7}$ |
| Fashion-MNIST-test | $44.5 \pm 0.5$ | $45.8 \pm 0.0$ | $28.8 \pm 0.0$ | $44.1 \pm 1.8$ | $51.8 \pm 2.4$ | - | $62.7 \pm 1.6$ | $\mathbf{68.2 \pm 1.8}$ |
| WebKB4 | $60.6 \pm 0.1$ | $47.5 \pm 0.1$ | $68.8 \pm 3.1$ | $43.6 \pm 0.4$ | $68.8 \pm 2.5$ | $52.1 \pm 0.2$ | $71.8 \pm 2.8$ | $\mathbf{83.2 \pm 1.6}$ |
| WebKB_cornell | $58.9 \pm 0.2$ | $54.4 \pm 0.6$ | $55.5 \pm 2.6$ | $42.6 \pm 0.0$ | $56.6 \pm 2.7$ | $49.6 \pm 0.2$ | $68.7 \pm 1.4$ | $\mathbf{74.4 \pm 2.1}$ |
| WebKB_texas | $59.4 \pm 0.2$ | $59.0 \pm 0.3$ | $64.5 \pm 3.0$ | $55.1 \pm 0.0$ | $64.1 \pm 3.6$ | $62.0 \pm 0.6$ | $73.8 \pm 1.2$ | $\mathbf{76.4 \pm 2.3}$ |
| WebKB_washington | $60.8 \pm 0.0$ | $51.7 \pm 1.0$ | $68.0 \pm 2.7$ | $46.5 \pm 0.0$ | $67.7 \pm 2.9$ | $65.4 \pm 0.4$ | $75.7 \pm 1.9$ | $\mathbf{79.3 \pm 1.2}$ |
| WebKB_wisconsin | $70.2 \pm 0.5$ | $72.8 \pm 1.4$ | $72.1 \pm 3.9$ | $46.1 \pm 0.0$ | $72.9 \pm 3.1$ | $73.2 \pm 0.9$ | $77.4 \pm 1.4$ | $\mathbf{81.6 \pm 2.2}$ |
| IMb_movies_keywords | $25.2 \pm 0.4$ | $24.0 \pm 0.2$ | $24.7 \pm 2.1$ | $12.6 \pm 1.7$ | $23.0 \pm 2.0$ | $23.3 \pm 1.1$ | $30.8 \pm 1.7$ | $\mathbf{29.3 \pm 1.1}$ |
| IMDb_movies_actors | $20.5 \pm 0.4$ | $20.0 \pm 0.4$ | $20.0 \pm 1.2$ | $14.1 \pm 2.8$ | $15.6 \pm 0.7$ | $15.8 \pm 1.3$ | $23.8 \pm 0.4$ | $\mathbf{26.2 \pm 2.4}$ |

Table 29: Clustering accuracy comparison with SRVCC

| Dataset | SCC | SBC | CCMod | DRCC | CCInfo | SCMK | DeepCC | **SRVCC** |
|---|---|---|---|---|---|---|---|---|
| Coil20 | $64.9 \pm 0.5$ | $73.9 \pm 1.1$ | $51.8 \pm 1.9$ | $65.6 \pm 2.7$ | $72.7 \pm 1.5$ | $72.5 \pm 0.9$ | $78.3 \pm 2.7$ | $\mathbf{75.0 \pm 2.1}$ |
| Yale | $41.6 \pm 0.3$ | $49.8 \pm 1.3$ | $24.6 \pm 2.3$ | $14.2 \pm 1.2$ | $48.5 \pm 2.0$ | $49.2 \pm 1.2$ | $55.7 \pm 1.1$ | $\mathbf{61.0 \pm 1.5}$ |
| Fashion-MNIST-test | $41.9 \pm 0.5$ | $41.3 \pm 0.0$ | $45.8 \pm 1.4$ | $42.2 \pm 1.6$ | $50.6 \pm 2.3$ | - | $60.4 \pm 0.7$ | $\mathbf{65.0 \pm 1.6}$ |
| WebKB4 | $31.1 \pm 0.1$ | $13.0 \pm 0.1$ | $40.1 \pm 1.0$ | $31.9 \pm 1.7$ | $39.7 \pm 3.6$ | $10.0 \pm 2.3$ | $40.5 \pm 0.6$ | $\mathbf{42.3 \pm 1.2}$ |
| WebKB_cornell | $28.8 \pm 0.2$ | $21.0 \pm 0.6$ | $18.9 \pm 3.8$ | $11.6 \pm 0.0$ | $20.6 \pm 3.1$ | $25.7 \pm 0.5$ | $35.4 \pm 0.9$ | $\mathbf{39.3 \pm 1.8}$ |
| WebKB_texas | $12.6 \pm 0.2$ | $9.0 \pm 0.3$ | $16.9 \pm 2.3$ | $10.2 \pm 0.0$ | $18.2 \pm 4.4$ | $24.0 \pm 0.8$ | $42.9 \pm 1.2$ | $\mathbf{43.5 \pm 1.7}$ |
| WebKB_washington | $25.3 \pm 0.0$ | $9.5 \pm 1.0$ | $28.7 \pm 1.4$ | $15.7 \pm 0.0$ | $30.7 \pm 3.4$ | $30.3 \pm 0.2$ | $45.9 \pm 1.3$ | $\mathbf{48.1 \pm 1.4}$ |
| WebKB_wisconsin | $35.4 \pm 0.5$ | $38.2 \pm 1.4$ | $35.1 \pm 2.8$ | $20.4 \pm 0.0$ | $39.3 \pm 2.7$ | $42.9 \pm 0.4$ | $46.7 \pm 1.7$ | $\mathbf{51.5 \pm 1.6}$ |
| IMDb_movies_keywords | $25.5 \pm 0.4$ | $20.6 \pm 0.2$ | $21.6 \pm 1.1$ | $6.9 \pm 0.3$ | $18.7 \pm 2.3$ | $18.4 \pm 0.8$ | $26.8 \pm 1.6$ | $\mathbf{25.3 \pm 1.2}$ |
| IMDb_movies_actors | $19.3 \pm 0.4$ | $17.6 \pm 0.4$ | $14.5 \pm 0.9$ | $9.3 \pm 2.5$ | $9.7 \pm 1.0$ | $10.6 \pm 1.7$ | $20.6 \pm 2.3$ | $\mathbf{19.4 \pm 1.8}$ |

Table 30: NMI Clustering results across various datasets.
